# Identifying non-coding variant effects at scale via machine learning models of cis-regulatory reporter assays

**DOI:** 10.1101/2025.04.16.648420

**Authors:** John C. Butts, Stephen Rong, Sager J. Gosai, Rodrigo I. Castro, Mackenzie Noon, Kehinde Adeniran, Rohit Ghosh, Pardis C. Sabeti, Ryan Tewhey, Steven K. Reilly

## Abstract

The inability to interpret the functional impact of non-coding variants has been a major impediment in the promise of precision medicine. While high-throughput experimental approaches such as Massively Parallel Reporter Assays (MPRAs) have made major progress in identifying causal variants and their underlying molecular mechanisms, these tools cannot exhaustively measure variant effects genome-wide. Here we present MPAC, an ensemble of machine-learning models trained on MPRA data that provides accurate and scalable prediction of the cis-regulatory impact of non-coding variants. Using MPAC we predict allelic effects for 575M single nucleotide variants (SNVs) across diverse applications, including complex trait genetics, clinical and tumor sequencing, evolutionary analyses, and saturation mutagenesis. We find MPAC predictions match the performance of empirical MPRAs in identifying causal complex trait-associated alleles. We demonstrate the utility of MPAC by applying it to ClinVar, identifying non-coding pathogenic variation with higher accuracy than other sequence-to-function models. We also nominate 1,892 candidate non-coding cancer drivers by predicting the functional effects of somatic SNVs in the COSMIC database. Next, we evaluate population-level genetic variation by predicting effects for all 514M non-coding SNVs in gnomAD, quantifying the relationship between regulatory function and evolutionary constraint. Finally, we generate prospective functional maps using *in-silico* saturation mutagenesis across 18,658 human promoters, observing widespread selection against variants predicted to disrupt promoter activity. Collectively, this study establishes the value of non-coding functional predictions and provides a comprehensive, publicly available resource for variant interpretation.

## Introduction

Over 1 billion ^1^ human single nucleotide variants (SNVs) have been identified to date from rapidly growing genomic sequencing efforts ^2–4^. As the vast majority of these variants reside in non-coding regions ^5–7^, a major unsolved task in human genomics has been to effectively identify variants with molecular functions that ultimately impact phenotype ^8,9^. Key examples include the over 700,000 significant associations between genomic loci and complex human diseases or traits discovered by genome-wide association studies (GWAS) ^10^, where identifying individual causal variants remains challenging due to linkage disequilibrium (LD) with non-functional variants. Similarly, studies of the genetic underpinnings of disease driven by rare ^11–15^, *de novo* ^16–19^, or somatic ^20–23^ mutations have identified more variants than are tractable to experimentally study. Evolutionary constraint has been used as a proxy for function to help prioritize a subset of putatively causal variants without needing to know the underlying mechanism ^24–29^. However, the relationship between constraint and function has not been comprehensively explored, especially in the non-coding genome.

Non-coding variants are highly enriched within *cis*-regulatory elements (CREs) such as promoters and enhancers, which precisely regulate gene expression in a spatio-temporal manner across cell types during development, homeostasis, and environmental response ^30,31^. However, our ability to interpret single-nucleotide changes is hindered by an incomplete understanding of regulatory grammar—the complex non-linear interactions between transcription factor (TF) sequence motifs, spacing, orientation, and context that define CREs ^6,7,32–36^. Large-scale efforts like ENCODE and others ^37–42^ have made substantial progress towards mapping the genomic locations of CREs across hundreds of cellular contexts. Association-based approaches, including molecular quantitative trait loci, have been used to map variants to function ^43–48^, but these approaches are limited to studying naturally occurring variation. Additionally, while models trained on epigenetic state including chromatin accessibility have emerged as powerful tools for predicting CRE activity ^49–57^, most lack training data that includes single-nucleotide changes, limiting their applicability to predicting variant effects.Massively parallel reporter assays (MPRAs) enable high-throughput, quantitative assessment of how DNA sequences regulate transcription, with the sensitivity to detect individual variants with small effects ^58–60^. MPRAs have been widely used to study the function of common variants from GWAS ^61–67^, rare and somatic mutations from disease cohorts ^21,68^, and putatively adaptive variants in human evolution ^69–73^. However, even the largest MPRAs, which analyze hundreds of thousands of naturally occurring variants ^66^, are outpaced by sequencing efforts that have identified over 1 billion ^1^ of the 9 billion SNVs possible in the human genome. To bridge this gap, machine learning models trained on large-scale MPRA data provide the opportunity to develop sequence-to-function models ^74^ and infer the *cis*-regulatory effects of untested variants ^56,75–80^. We recently developed a highly accurate convolutional neural network model of CRE activity, Malinois, trained on MPRA data from 776,474 sequences, the vast majority of which represent variant alleles assayed in three cell lines ^78^.

Here we introduce MPAC (Malinois with Parallel Aggregated Cross-validation), a scalable framework based on Malinois that is adapted for predicting changes in MPRA-measured CRE activity caused by SNVs. MPAC matches or exceeds the performance of existing variant effect prediction tools trained on larger or orthogonal datasets ^81–83^. We demonstrate MPAC’s scalability by generating hundreds of millions of predictions to identify causal and pathogenic variants, non-coding drivers of cancer, and the relationship between non-coding and coding constraint genome-wide.

## Results

### MPAC accurately captures variant effects of regulatory features

To develop MPAC, we introduced several essential modifications to Malinois’ CRE prediction framework, enabling genome-wide predictions of variant effects (**Figure 1A**) (**Methods**). First, we ensured unbiased CRE activity predictions for any human genomic sequence by avoiding overfitting to data seen during training. Autosomes were split into eleven pairs and each pair was used as the test set to train ten models (based on the Malinois architecture) per pair, for a total of 110 separate models (**Methods**). For any query sequence, the ten models not trained on the query’s source chromosome were used for prediction and ensembled by averaging to give a final *cis*-regulatory activity (log_2_FC) score. Across three cell types, MPAC predicts empirical MPRA activity with high accuracy (K562: Pearson’s *r* = 0.89, *n* = 495,180 sequences; HepG2: *r* = 0.89, *n* = 499,820; and SK-N-SH: *r* = 0.88, *n* = 485,034) (**Fig. 1B**), and high consistency across all chromosomes: (K562: *r* = 0.88-0.91, HepG2: *r* = 0.88-0.91, and SK-N-SH: *r* = 0.87-0.90) (**Supplementary Fig. 1**) (**Supplementary Dataset 1**). We further adapted MPAC for variant effect prediction by incorporating a larger sequence context, which has been shown to capture additional signals contributing to activity in reporter assays ^84,85^. Allelic skew of a variant is calculated as the difference in predicted activity between the alternate and reference sequences, based on averaging predictions for 18 sliding 200-bp windows upstream and downstream of the variant in 10-bp increments (**Supplemental Fig. 2**) (**Methods)**. MPAC allelic-skew predictions are highly correlated with empirical allelic skews for expression modulating variants (emVars) from MPRA in all three cell types (K562: Pearson’s *r* = 0.71, *n* = 16,187 sequence pairs; HepG2: *r* = 0.76, *n* = 14,029; SK-N-SH: *r* = 0.69, *n* = 14,723) (**Fig. 1C**). While empirical MPRA emVars can be determined using a *p*-value threshold, we define MPAC emVars using an absolute skew threshold (|allelic-skew log_2_FC| > 0.5), a comparable threshold to previous studies that also yields similar emVar rates ^66^.

**Fig. 1:**
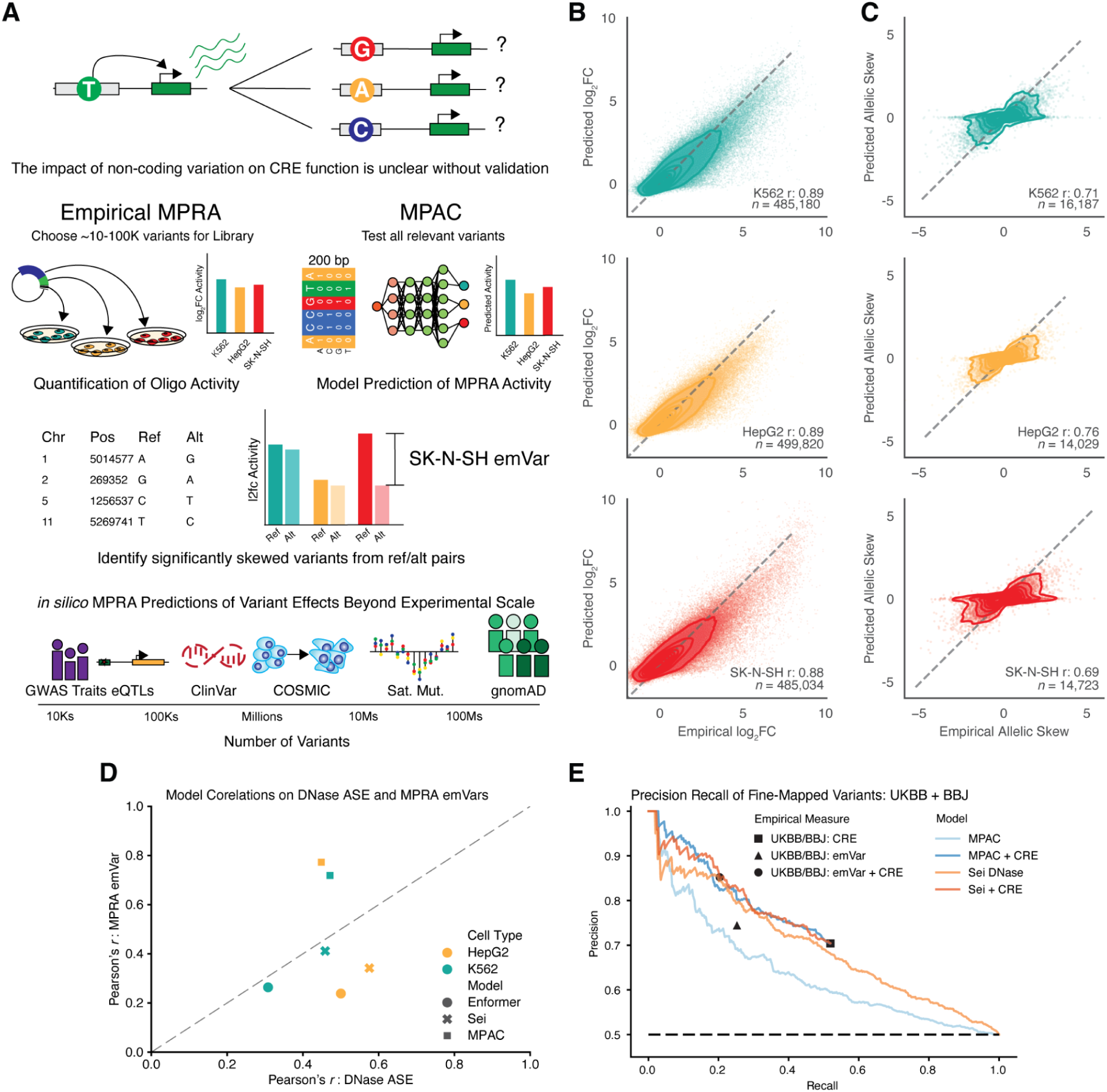
MPAC is a highly accurate model of MPRA activity for scalable prediction of allelic skew. **A)** Schematic illustrating the difficulty in interpreting non-coding regulatory variants and how empirical MPRAs or the predictive MPAC model can be used to identify expression modulating variants (emVars). **B)** Scatterplot of MPAC activity predictions and empirical MPRA activity in K562 (*n* = 485,180), HepG2 (*n* = 499,820), and SK-N-SH (*n* = 485,034). **C)** Scatterplot of MPAC allelic skew (log2FC alternate activity - reference activity) and empirical skew of MPRA emVars inK562 (*n* = 16,187), HepG2 (*n* = 14,029), and SK-N-SH (*n* = 14,723). **D)** Scatterplot of MPAC, Enformer, and Sei Pearson’s correlation to MPRA emVars (y-axis) and DNase ASE (x-axis). **E)** Precision-recall curves of causal complex trait-associated variation (high PIP > 0.9, *n* = 934 vs. low PIP <0.01, *n* = 935) for CREs (black square), empirical emVars (black triangle), emVar + CRE (black circle), MPAC (light blue), MPAC + CRE (dark blue), Sei (light orange), and Sei + CRE (dark orange).

For a fraction (13%) of variants, MPAC-predicted and empirical allelic-skew emVar calls were discordant, defined as being an emVar in one study and non-emVar with |log_2_FC| < 0.05 in the other. Discordancies were entirely due to false negative MPAC predictions. To assess if missed predictions were due to specific sequence features or motifs, we trained a gkm-SVM model ^86^ to identify k-mers that discriminate poorly predicted emVars (**Supplementary Fig. 3**). Top scoring k-mers identified by GkmExplain ^87,88^ were enriched for AT-homopolymers and those matching known PRDM6, FOXJ3, MEF2B, LHX3, FOXQ1, and ZN770 motifs (**Supplementary Fig. 4**). However, *de novo* motifs with matches to known motifs were infrequently identified in emVars, collectively overlapping just 0.9%, 1.81%, and 4.32% of K562, HepG2, and SK-N-SH sequences respectively (**Methods**), suggesting missed MPAC predictions are largely due to sequence features infrequently seen in training.

We next compared MPAC predictions to orthogonal measures of regulatory variant function (**Supplementary Dataset 2**). Although MPRA variant effects on reporter expressionand DNase allele-specific effects (ASE) on chromatin accessibility capture distinct features of gene regulation, we found MPAC-predicted allelic skew and DNase ASE were significantly correlated for variants with significant DNase ASE (K562: Pearson’s *r* = 0.42, *p =* 3.41 x 10^−54^; HepG2: *r* = 0.44, *p =* 6.82 x 10^−24^) (**Fig. 1D**) (**Methods**). Subsetting to variants confidently detected in both assays (MPAC emVar and DNase ASE) greatly increased correlation (K562: *r* =. 0.71; HepG2: *r* = 0.73) (**Supplementary Fig. 5A, C**) likely by removing variants that alter chromatin accessibility independently of CRE activity ^89^. Notably, these predicted results were comparable to correlations between empirically determined emVar allelic skew and DNase ASE (K562: *r* = 0.7 and HepG2: *r* = 0.73) (**Supplementary Fig. 5B, D**). MPAC approached or outperformed two leading sequence-to-function deep learning models on this task despite being trained on considerably less data overall and no DNaseI hypersensitivity data ^81,82^ (Sei: K562 *r* = 0.46, HepG2 *r* = 0.58; Enformer: K562 *r* = 0.31, HepG2 *r* = 0.50) (**Fig. 1D, Supplementary Fig. 6 and 7**) (**Methods**). Lastly, Enformer DNase ASE predictions showed low correlation to MPRA allelic skews in both K562 (*r* = 0.26, *p* = 8.13 x 10^−190^, *n* = 11,994) and HepG2 (*r* = 0.24, *p* = 1.55 x 10^−134^, *n* = 10,407) (**Fig. 1D**, **Supplementary Fig. 8**) (**Methods**) while Sei DNase ASE predictions showed modest correlation in both K562 (*r* = 0.41, *p* < 10^−300^, *n* = 11,994) and HepG2 (*r* = 0.34, *p* = 1.99 x 10^−283^, *n* = 10,407) (**Fig. 1D**, **Supplementary Fig. 9**).

To assess how effectively MPAC prioritizes trait-associated causal variants, we compared its predictions against empirical MPRA measurements for 217,577 statistically fine-mapped complex trait-associated variants from UK BioBank, BioBank Japan, and eQTLs from the Genotype-Tissue Expression project (GTEx) ^66^ (**Supplementary Dataset 1**). For complex trait associations, MPAC performed comparably to empirical MPRA at identifying causal variants (posterior inclusion probability (PIP) > 0.9) vs. non-causal variants (PIP < 0.1) within CREs (precision = 0.82 vs. 0.85 at recall = 0.20) (**Fig. 1E**). MPAC matches Sei (AUPRCs of 0.83 for both models) (**Fig. 1E**) and outperformed the *k-*mer based deltaSVM model ^83^ (AUPRC 0.83 vs. 0.74) (**Supplementary Fig. 10**). For molecular traits, MPAC identified causal eQTL variants within CREs on par with empirical MPRA (precision 0.81 vs. 0.82 at empirical recall of 0.15). MPAC was slightly outperformed by Sei (AUPRC 0.78 vs. 0.81) but outperformed deltaSVM (AUPRC 0.78 vs. 0.72) (**Supplementary Fig. 11**). Overall, MPAC is able to prioritize candidate causal variants with precision comparable to empirical MPRA, at vastly reduced cost, time, and labor.

### MPAC discerns clinically relevant germline and somatic variants

The inability to quickly categorize non-coding variants as pathogenic or benign limits our understanding of their contribution to rare or Mendelian diseases ^90,91^. Since CREs, as determined by DNase I Hypersensitivity Sites (DHS), show weak enrichment for pathogenic variants compared to non-CRE regions (*OR* = 1.22, *p* = 0.02), we sought to determine whether MPAC predictions provide better discrimination. We applied MPAC to all 180,032 non-coding variants in the ClinVar database (**Figure 2A**) (**Methods**) (**Supplementary Dataset 3**) and found that MPAC emVars exhibited greater enrichment for pathogenic variants when considering all non-coding ClinVar variants (*OR* = 3.48, *p* = 2.25 x 10^−11^) (**Supplementary Fig. 12A**), increasing further when restricting to CREs (*OR* = 12.09, *p* = 4.77 x 10^−15^, Fisher’s exact test) (**Supplementary Fig. 12A**). We next compared the precision and recall performance of MPAC with Sei’s DHS allelic-effect predictions using matched sets of pathogenic and benign variants (**Methods**). Across all non-coding ClinVar variants, both MPAC and Sei performed well, with MPAC achieving a mean AUPRC of 0.65 and Sei 0.61. Performance improved to 0.68 (MPAC) and 0.64 (Sei) when restricting variants to DHS regions, and further increased within promoters (MPAC: 0.82; Sei 0.78) (**Fig. 2B**, **Supplementary Fig. 12B**).

**Fig. 2:**
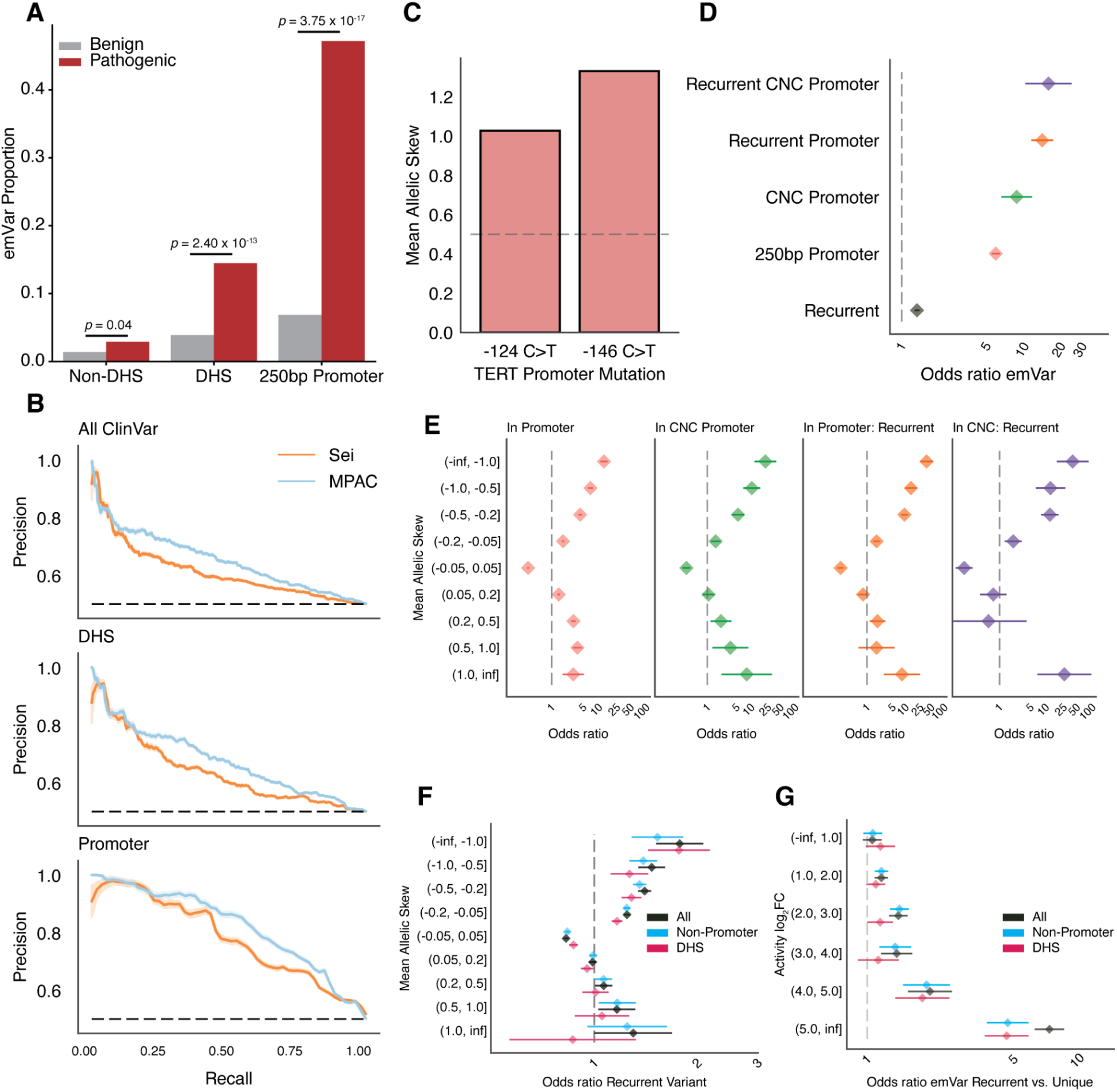
MPAC identifies non-coding pathogenic variants and recurrent cancer drivers. **A)** Bar plot showing the proportion of ClinVar pathogenic and benign emVars in different genomic contexts. DHS: DNase I hypersensitive site. *p*-values from Chi-Square Test. **B)** Precision-recall curves of ClinVar pathogenic vs. matched downsampled benign non-coding variants (All ClinVar: *n* = 573; DHS: *n* = 228 pathogenic, 228 benign; Promoter: *n* = 53 pathogenic). **C)** Mean allelic skew of TERT promoter mutations. **D)** Odds ratio (*OR*) that a variant is an emVar in COSMIC subsets. CNC: Cornell Non-Coding Cancer Driver Database. **E)** *OR* that a variant is in promoter annotations by mean skew bin. **F)** *OR* that a variant is recurrent compared to unique binned by mean skew. **G)** *OR* that a recurrent variant is an emVar compared to unique variants binned by sequence activity. Error bars in panels D-G represent 95% CIs.

Individual tumors can harbor tens to hundreds of thousands of somatic mutations ^92,93^, the majority of which are likely non-functional passenger mutations, making the identification of driver mutations experimentally intractable. Large sequencing cohorts can infer variant function by identifying mutations that recur across patients, as such recurrence is unlikely to occur by chance. However, non-coding mutations have proven difficult to identify ^94^ with only a few identified as important for tumorigenesis ^22,93,95,96^. A well-characterized example of recurrent, non-coding driver mutations are those reactivating TERT by creating ETS-family TF binding sites in its promoter ^95–97^. These two known mutations, −124 (C>T) and −146 (C>T) were predicted by MPAC to be emVars, increasing expression (−124 C>T: mean allelic skew = 1.03 and −146 C>T: mean allelic skew = 1.33) (**Fig. 2C**). Globally, ETS-family binding sites have been shown to be enriched for somatic mutations at promoters using a TF-aware burden test and functional validation ^98^. Using MPAC, we detected 61% of the ETS-family binding putative non-coding driver mutations as emVars, supporting our *in silico* approach to nominate variants associated with tumorigenesis (**Methods**).

Encouraged by our validation of known TF binding sites, we assessed emVar enrichments for somatic mutations more broadly in the Catalog of Somatic Mutations in Cancer (COSMIC v98) (**Supplementary Dataset 4**). We observed that across all promoters, variants within 250 bp of a transcription start site (TSS) were more likely to be emVars than those in distal regions (*OR* = 5.95, *p* = 2.34 x 10^−288^) (**Fig. 2D**). When subsetting to the promoters of genes previously associated with cancers from the Cornell Non-Coding Cancer Driver Database (CNC) (**Methods**) ^99^ this enrichment increased (*OR* = 8.86, *p* = 4.13 x 10^−31^). Notably, regardless of inclusion in CNC, variants in promoters showed the greatest enrichment for strong effect mutations (allelic skew ≤ −1 or ≥ 1) (**Fig. 2E**), with a bias towards loss-of-function (LoF) mutations when considering all promoters (**Fig. 2E**). Finally, when comparing mutations that occur across multiple individuals to those appearing once, we observed that recurrent variants in promoters were significantly more likely to be emVars (*OR* = 14.35, *p* = 6.81 x 10^−84^). Recurrent promoter variants for CNC genes displayed the highest emVar enrichment (*OR* = 16.18, *p* = 9.18 x 10^−22^) (**Fig. 2D**). We next considered all 2,432,725 variants in COSMIC to determine whether MPAC identified functional enrichments beyond promoters. Across the entire dataset, recurrent variants were enriched for MPAC-emVars compared to variants observed in only one individual (*OR* = 1.34, *p* = 9.76 x 10^−32^) (**Fig. 2D**). When recurrent and unique variants were partitioned by mean allelic skew, we identified enrichments for large negative skew variants for all recurrent variants, variants outside of promoters, and those within DHS elements (allelic skew ≤ −1) (**Fig. 2F**). Furthermore, we expect recurrent driver mutations to be enriched in sequences with high predicted CRE activity. Activity predictions from MPAC correlated with DHS element likelihood; notably, 98% of emVars with a mean reference log_2_FC > 5 overlapped DHS elements. We observed enrichment between active sequences (all bins log_2_FC > 1, All COSMIC variants) and recurrence (*OR* = 1.17, *p* = 4.41 x 10^−5^), which is greatest at highly active sequences (log_2_FC > 5, *OR* = 7.39, *p* = 8.62 x 10^−85^) (**Fig. 2G**). This enrichment persisted after excluding promoters both genome-wide and within DHS peaks (**Fig. 2G**). In total, we nominated 117 promoter and 1,775 non-promoter recurrent emVars as candidate non-coding drivers across COSMIC.

### Scaling MPAC predictions to common and rare human genetic variation reveals patterns of functional constraint

To systematically evaluate the functional consequences of rare and common variation from globally diverse human populations, we first used MPAC to assess the regulatory activity of 514 million non-coding SNVs from the 76,156 genomes in The Genome Aggregation Database (gnomAD v3.1.2) (**Supplementary Fig. 13) (Methods) (Supplementary Dataset 5**). We observed that 10.0% (51.5M) of all variants were predicted to be in active sequences(log_2_FC > 1). We intersected all gnomAD variants with ENCODE candidate CREs (cCREs) which cover 7.9% of the genome ^37^, identifying 0.4% (1.9M) in promoter-like signatures (PLS), 2.1% (10.9M) in proximal enhancer-like signatures (pELS), 14.3% (73.6M) in distal enhancer-like signatures (dELS), 5.7% (29.4M) in additional cCREs such as TF binding sites (other cCREs), and 77.5% (398M) in regions outside of cCREs (non-cCRE) (**Fig. 3A**). As expected, variants overlapping PLS showed the highest average predicted activity (log_2_FC = 2.26), followed by pELS (log_2_FC = 0.65) and dELS (log_2_FC = 0.48), while variants outside of cCREs showed minimal activity (log_2_FC = 0.25), suggesting that MPAC precisely captures the activity difference of cCRE annotations (**Fig. 3B**).

**Fig. 3:**
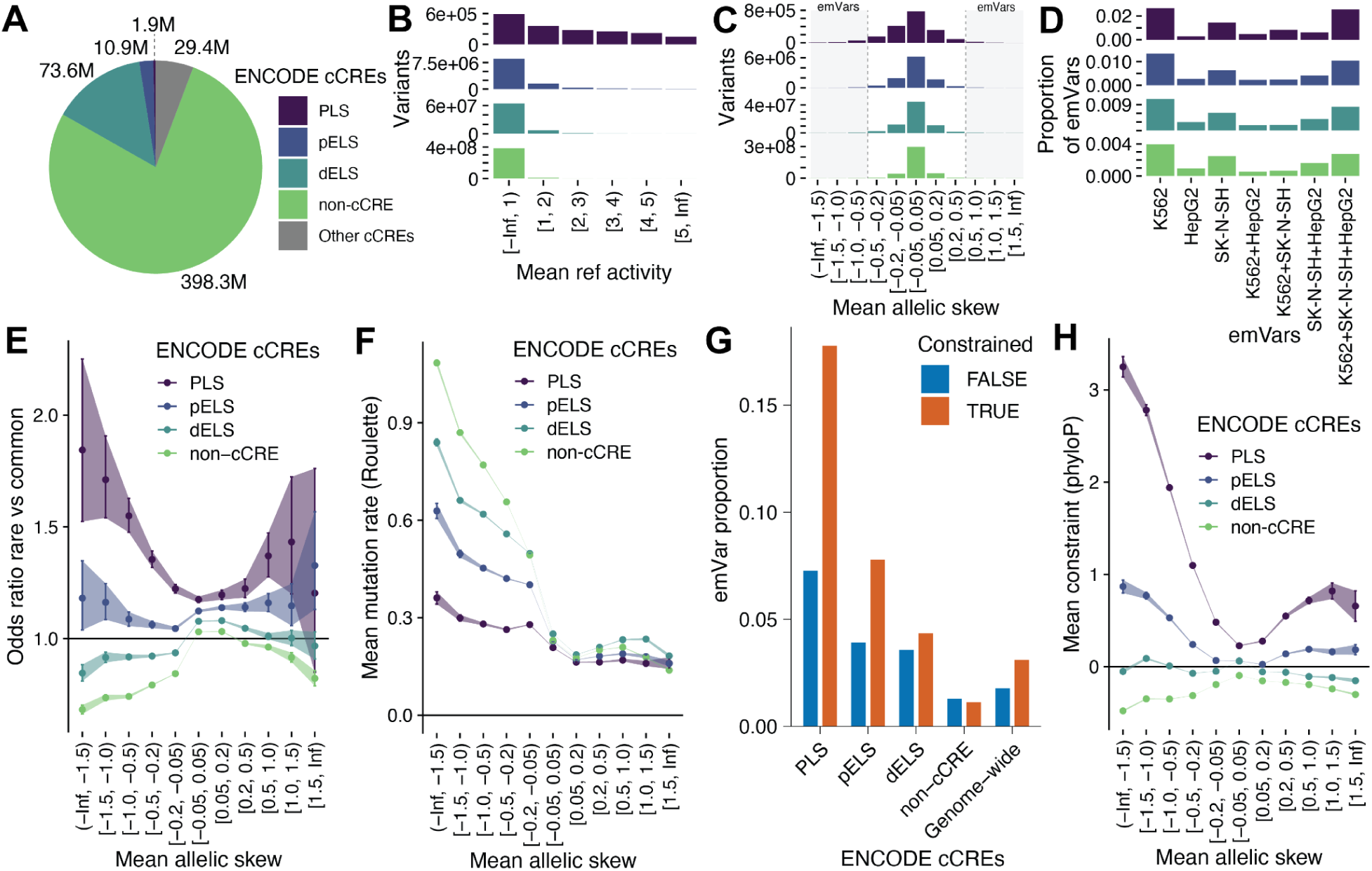
MPAC-predicted emVars in gnomAD are under strong purifying selection. **A)** Overlap of non-coding gnomAD SNVs with ENCODE cCRE classes including PLS, dELS, dELS, other cCREs, and non-cCRE regions. Distribution of gnomAD SNVs across cCRE classes based on **B)** mean MPAC activity (log_2_FC) predictions and **C)** mean MPAC allelic-skew (log_2_FC) predictions. **D)** Proportion of expression-modulating variants (emVars) in one or more cell lines by cCRE classes. **E)** Recent purifying selection, quantified by the OR of rare (MAF < 0.1%) vs. common (MAF > 0.1%) SNVs relative to genome-wide baseline, across cCRE classes with different levels of mean MPAC allelic-skew predictions. **F)** Mean mutation rates, as predicted by Roulette. **G)** Proportion of variants called emVars at constrained and unconstrained bases in different cCRE classes (phyloP > 2.27, FDR < 0.05). **H)** Evolutionary constraint of gnomAD SNVs, measured by mean Zoonomia phyloP score. Error bars in panels E-G represent 95% CIs.

Next, we used MPAC to predict allelic skew for all gnomAD variants ^65,66,80,100^. MPAC predicted 9.4M (1.8%) of all variants to be emVars, consistent with rates in empirical MPRA studies ^61–63^. As expected due to their overall higher constraint, PLS contained the highest proportion of emVars (8.83%) (**Fig. 3C**) and displayed the largest absolute allelic skews (PLS vs. genome-wide mean |log_2_FC| = 0.139 vs. 0.057) (**Fig. 3C**). Consistent with promoters’ more ubiquitous function, cell type specificity of emVars was lowest in PLS (49.5%), but increased in contexts farther from the TSS (pELS: 53.7%, dELS: 55.5%) (**Fig. 3D**), consistent with the established role of distal CREs in driving cell type-specific gene expression (**Supplementary Fig. 14**)^101,102^. As a resource, we also provide allelic-skew predictions for lead SNVs and variants in high LD (*r*^2^ ≥ 0.7) for SNVs found in the 5,781 studies in the GWAS Catalog ^10^ (**Supplementary Dataset 6**) (**Methods**). Across all trait-associated SNVs assessed, 1.4% (33,808 out of 2,415,078) are predicted to be emVars and 24.6% (184,663 out of 750,574) of trait-associations have at least one predicted emVar. Lead SNVs were more likely to be emVars than SNVs in LD (*OR* = 1.31-1.43 across populations, *p* < 10^−31^).

As CRE activity is largely defined by the combinatorial binding of transcription factors (TF) to accessible chromatin ^35,103^, we hypothesized emVars should be enriched at accessible and TF-bound regions. gnomAD variants with large predicted effect sizes, defined as having an allelic skew |log_2_FC| > 1.5, were highly enriched in DNase I TF footprints ^104^ (negative: *OR* = 8.70, positive: *OR* = 3.11) and TF ChIP-seq peaks ^105,106^(negative: *OR* = 2.31, positive: *OR* = 2.12) (**Supplementary Fig. 15A,B**). Conversely, variants predicted to have little to no impact (allelic skew |log_2_FC| < 0.05) were depleted for TF binding (TF footprints: *OR* = 0.37, TF ChIP-seq: *OR* = 0.62). However, the vast majority of emVars were located outside of annotated

TF footprints (85.9%, 3.77M vs. 617k) or TF ChIP-seq peaks (92.1%, 4.04M vs. 346k) (**Supplementary Fig. 15C,D**), highlighting the limitations of current TF binding annotations. Non-coding functional variants are likely to be deleterious and thus under constraint within-species ^107–109^. However, this relationship has only been explored with MPRAs for a small subset of CREs ^110^, as most past studies ascertained variants biased by their trait associations. We stratified all observed gnomAD variants by MPAC allelic-skew predictions and used the odds ratio (*OR*) of rare SNVs (allele frequency (AF) ≤ 0.1%) vs. common SNVs (AF > 0.1%) as a proxy for the overall strength of recent purifying selection. Starting with variants in PLS, we observed higher constraint at extremes of allelic skew compared to variants with less or no allelic skew (**Fig. 3E, Supplementary Fig. 16A-C)**. This U-shaped relationship was asymmetric, with large negative allelic-skew variants displaying the strongest purifying selection (*OR* = 1.84) compared to those with minimal allelic skew (|log_2_FC| < 0.05, *OR* = 1.17). While this pattern is consistent with purifying selection against deleterious promoter loss-of-function mutations, mutations with large positive allelic skews also displayed purifying selection (*OR* = 1.20). We see a similar pattern for variants overlapping pELS, but not dELS or non-cCRE contexts (**Fig. 3E**). Lastly, we found that the predicted *de novo* mutation rate increases with the magnitude of negative allelic skews (**Fig. 3F, Supplementary Fig. 16D-F**). This trend was strongest for variants outside cCREs, and within cCREs was strongest for more promoter-distal cCREs.

### MPAC enables a comprehensive exploration of the evolutionary constraint-function relationship

Variant effect predictors have used evolutionary constraint across species as an indirect measure to assess potential pathogenicity, but this relationship has not been comprehensively evaluated for non-coding regions ^111,112^. To evaluate if MPAC could effectively identify constrained, functional regulatory variants, we compared predicted emVars to evolutionary constraint maps at base-level resolution from the Zoonomia 241-way mammalian alignment which defines constrained bases as having phyloP > 2.27 (FDR < 0.05) ^26^. As expected, within known cCREs, gnomAD variants overlapping constrained bases were significantly more likely to be emVars than those overlapping unconstrained bases (*OR* = 1.77, *p* < 2.2 x 10^−16^, Fisher’s exact test) (**Fig. 3G**). Across cCRE classes, the highest enrichment was found in promoters (PLS: *OR* = 2.76, pELS: *OR* = 2.07, dELS: *OR* = 1.22, and non-CRE: *OR* = 0.87, all *p* < 2.2 x 10^−16^) (**Fig. 3G**). However, among the 322,644 emVars with evidence of constraint, the majority were found in pELS and dELS (65.2%) compared to PLS (15.1%), reflecting their larger portion of the genome (**Supplementary Fig. 17**).

MPAC’s quantitative predictions further allowed us to examine how the magnitude and directionality of variant function relates to evolutionary constraint. We compared phyloP scores to MPAC allelic-skew predictions across cCRE classes. Similar to our findings using within-species constraint, variants in PLS exhibited a U-shaped relationship between cross-species constraint and allelic skew (**Fig. 3H**, **Supplementary Fig. 18A-C**). Variants with large negative allelic skew displayed very strong levels of cross-species constraint (mean phyloP = 3.25), much higher than those with no allelic skew (mean phyloP = 0.23). Comparatively, variants with large positive allelic skew also showed increased constraint, albeit more modest (mean phyloP = 0.66). The constraint of the most negatively impactful variants was slightly higher than that of start loss variants, splice donor, and splice acceptor variants (mean phyloP = 3.14, 2.87, 2.85, respectively) and almost as high as that of missense and stop gained variants (mean phyloP = 3.61, 3.45, respectively) (**Supplementary Fig. 18D**). Thus, MPAC can prioritize non-coding variants with functional impacts as deleterious as coding mutations.

We next looked at non-promoter regions, which prior studies have shown display a weaker relationship between function and constraint ^113–115^. Consistent with these findings, we detected a similar but weaker U-shaped relationship between constraint and function in pELS regions, with large negative and positive allelic skews under more constraint (phyloP = 0.87, 0.18) than those with minimal effects (phyloP = 0.07). Comparatively, we did not observe this pattern for variants in dELS (**Fig. 3H**). Similarly, we observed that pleiotropy, as measured by sharing of emVars across cell types, increased with constraint in PLS and pELS, but not dELS (**Supplementary Fig. 19A**). Notably, MPAC activity of the element harboring the variant did not correlate strongly with constraint (**Supplementary Fig. 19B**), underscoring the importance of variant-level functional predictions. The overall weaker relationship between function and constraint at non-promoter CREs is consistent with their greater evolutionary turnover ^116^ and functional redundancy resulting in phenotypic robustness ^117^.

While purifying selection prevents deleterious variants from reaching high allele frequencies, most variants are expected to be non-functional and the majority of variants in gnomAD are rare (AF < 0.1%). Evolutionary constraint has a moderate ability to distinguish rare from common variation (phyloP = −0.30 vs −0.08) (**Supplementary Fig. 20A**), but the constraint threshold distinguishing functional from non-functional regulatory variants is unknown. We hypothesized that stratifying variants by their regulatory impacts could decompose this relationship. As expected, across all allelic-skew bins, constraint increased with allele rarity; however this relationship varied substantially across different regulatory contexts (**Supplementary Fig. 20B**). Variants in PLS showed the strongest signals of purifying selection, with the highest constraint observed for singleton variants with large negative allelic skew (phyloP = 3.59) compared to either singletons with minimal predicted skew (phyloP = 0.31) or all singletons without functional stratification (phyloP = 0.56). For ultra-rare PLS variants, constraint was similarly much higher at functional positions (phyloP = 3.09) compared to non-functional (phyloP = 0.23). Variants in pELS displayed a similar but more modest pattern, whereas variants in dELS and those outside of cCREs consistently showed the lowest constraint and weakest relationship with allele frequency.

### Prospective in-silico saturation mutagenesis of all promoters

We deployed MPAC to prospectively and comprehensively map the regulatory impact of all possible SNVs in human promoters genome-wide (**Supplementary Dataset 7**). We generated MPAC predictions for all three non-reference single nucleotide mutations within the promoter of 18,658 protein-coding genes, defined as positions −1000 bp to −1 bp upstream of the canonical TSS (**Methods**). Most (53.4%) emVars are clustered within the proximal 250 bp upstream of the TSS, where they displayed both higher mean activity and greater mean allelic skew than emVars found in more distal regions (**Supplementary Fig. 21A,B**). Interestingly, two-thirds (67.2%) of emVars are predicted to reduce promoter activity (mean allelic skew = −0.64) instead of increasing activity (mean allelic skew = 0.34), suggesting functional promoter mutations are more likely to be disruptive. Activity and allelic skew were most correlated across cell types in the 250 bps proximal to the TSS (activity: Spearman’s *ρ* = 0.93-0.97; allelic skew: *ρ* = 0.74-0.83) (**Supplementary Fig. 22A-D**), and decreased with distance. Moreover, constraint (phyloP score) (**Supplementary Fig. 21C**) and correlation between constraint and allelic skew (**Supplementary Fig. 22E-H**) were highest proximal to the TSS, consistent with the central role of the core initiation site.Focusing on the core promoter (proximal 250 bp to TSS), we next sought to understand if MPAC predictions captured known features of promoter initiation. Across all three cell types, we observed high predicted regulatory activity broadly from −150 to 0 bp from the TSS (average log_2_FC across all promoters and cell lines) **(Fig. 4A)**, suggesting sequences in this region may be particularly sensitive to mutations. Within this range, we found two regions particularly concentrated in positions of large allelic skew (average log_2_FC of three possible mutations across all promoters and cell lines) **(Fig. 4B)** and coinciding with elevated evolutionary constraint, suggesting underlying sequence features drive these patterns **(Fig. 4C)**. We next asked if known promoter motifs^118^ might be disrupted at these regions, identifying broadly negative allelic skew extending from the TSS to −26. As expected, this region harbors core promoter elements for the TATA box, NRF1, and ELK1 **(Fig. 4D, Supplementary Fig. 23)**. Further upstream, we observed a strong skew peak centered at −45 that co-localised with the promoter-associated TF SP1 motif **(Fig. 4D, Supplementary Fig. 23)**. We also found that predicted activity of promoters was significantly correlated with gene expression levels in all three cell types (Pearson’s *r* = 0.59-0.53, all *p* < 2.2 x 10^−16^) (**Supplementary Fig. 24**). Together, these findings show the ability of MPAC saturation mutagenesis to recapitulate known aspects of promoter regulatory grammar.

**Fig. 4:**
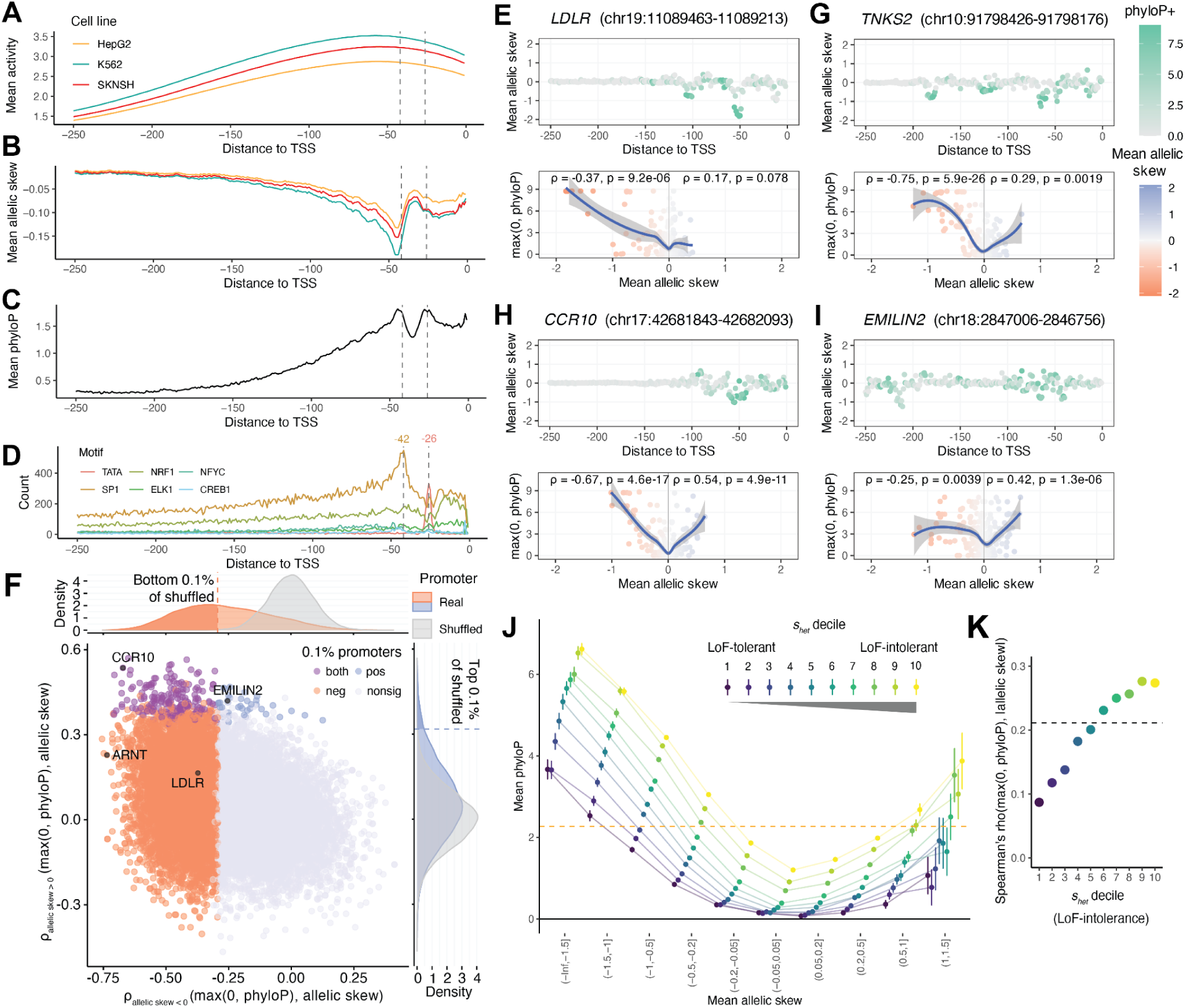
Saturation mutagenesis of 18,658 promoters reveals interplay between variant function, LoF tolerance, and constraint. Meta-promoter plots of mean MPAC-predicted **A)** activity (log2FC) and **B)** allelic skew (log2FC) from in silico saturation mutagenesis of the 250-bp promoter region upstream of the TSS of 18,658 canonical protein-coding genes. **C)** Meta-promoter plots of evolutionary constraint (mean phyloP). **D)** Motif counts by distance to TSS of representative upstream promoter-binding transcription factors TATA-binding protein, NRF1, NFYC, SP1, ELK1, and CREB1. Vertical lines correspond to modes of TATA and SP1 motif counts. **E)** Promoter-level plots of mean allelic skew by distance to the TSS colored by max(0, phyloP) (top) and the correlation (ρ and p-values) of max(0, phyloP) and allelic skew for mutations with negative (bottom left) or positive (bottom right) allelic skew for *LDLR*. **F)** Promoter-level correlations (ρ) between constraint max(0, phyloP) and MPAC allelic-skew predictions for the subset of SNVs with negative (x-axis) or positive (y-axis) allelic skew. Colors highlight promoters with ρ in bottom 0.1% for negative allelic-skew mutations (red) or top 0.1% for positive allelic-skew mutations (blue) relative to permutation-based nulls. Plotting the subset of 16,995 promoters for which both correlations are well-defined (see **Methods**). Example promoters with strong correlations between phyloP and allelic skew for **G)** negative allelic-skew mutations only, *TNKS2*; **H)** both negative and positive allelic-skew mutations, *CCR10*; and **I)** positive allelic-skew mutations only, *EMILIN2*. **J)** Relationship between promoter variant function (mean allelic skew) and constraint (mean phyloP) stratified by gene-level loss-of-function (LoF) intolerance (shet decile). Error bars represent 95% CIs. Constraint cutoff (phyloP = 2.27, FDR < 0.05) shown as orange line. **K)** Spearman’s ρ between variant function (|allelic skew|) and constraint (max(0, phyloP)) by shet decile and genome-wide (dashed line).

Saturation mutagenesis MPRAs have previously observed variable correlations between constraint (phyloP) and variant function (empirical allelic skew) from studies of a limited number of promoters ^59,66,119,120^ ^27,119^, so we explored the strength and generality of this relationship across all promoters. At the previously examined *LDLR* (low-density lipoprotein receptor) promoter ^27,119^, we separately compared negative LoF and positive gain of function (GoF) allelic-skew mutations to constraint and confirmed the previously observed correlation. Notably, the correlation was driven by negative allelic skew (Spearman’s *ρ* = −0.37, *p* = 9.2 x 10^−6^) (**Fig. 4E**) with no significant correlation between phyloP and positive allelic-skew mutations (*ρ* = 0.17, *p* = 0.078) (**Fig. 4E**), attributable to the lack of possible SNVs with appreciably positive allelic skew (no SNVs with allelic skew > 0.5).

Analyzing all promoters (**Supplementary Dataset 8)**, 27% had correlations between phyloP and allelic skew more extreme than *LDLR* (**Fig. 4F; Supplementary Fig. 25**). The most highly correlated example was the promoter of the broadly expressed *TNKS2*, which encodes a polymerase involved in various processes including Wnt/β-catenin signaling, telomere maintenance, and DNA repair ^121^ (*ρ* = −0.75, *p* = 5.9 x 10^−26^) (**Fig. 4G**). We found that 1.1% (*n* = 192) of promoters exhibited significant correlation between allelic skew and phyloP for both LoF (*ρ* = −0.54, *p* = 4.9 x 10^−11^) and GoF (*ρ* = −0.67, *p* = 4.7 x 10^−17^) mutations, suggestive of stabilizing selection (**Fig. 4H**; **Supplementary Fig. 26**). Comparably, 42.5% (*n* = 7,645) of promoters had a significant correlation only at LoF mutations, indicating the commonality of strong purifying selection against reductions in promoter activity (permutation *p* < 0.001). Only 0.2% (*n* = 38) of promoters showed significant correlation only at GoF mutations, including *EMILIN2* (*ρ* = 0.42, *p* = 1.3 x 10^−6^) (**Fig. 4I**), an extracellular matrix protein ^122^ (**Supplementary Fig. 27**). Genes with significant correlation between only negative allelic skew and phyloP had higher gene-level coding constraint (*p* < 0.001, Wilcoxon signed rank test) (**Supplementary Fig. 28**) and were enriched in pathways related to RNA or protein metabolism, cell cycle regulation, signal transduction, neurodevelopment, cellular stress responses, and embryonic lethality (**Supplementary Dataset 9**). MPAC thus enables improved regulatory variant interpretation and novel gene prioritization strategies, accounting for both the magnitude and direction of expression effects in the context of evolutionary constraint.

Constraint in promoter regions can reflect both *cis*-regulatory and downstream gene-level fitness effects. We explored this relationship using MPAC allelic skew to assign *cis*-regulatory impact, *s_het_* to quantify gene-level loss-of-function (LoF) intolerance, and phyloP to measure constraint ^123–125^ (**Fig. 4J**). We observed that genes with higher *s_het_* had higher correlations between allelic skew and phyloP (top 10%: *ρ* = 0.27 vs. bottom 10%: *ρ* = 0.087) (**Fig. 4K**), suggesting that the importance of the gene strongly influences the relationship between non-coding effect and constraint. For large negative skew mutations in promoters of the most LoF-intolerant genes, we observed extremely elevated constraint (phyloP = 6.52) (**Fig. 4J**). Expression-increasing mutations in the same promoters also showed significant, albeit reduced, levels of constraint (mean phyloP = 3.87). Surprisingly, large negative effect mutations for the most LoF-tolerant genes were highly constrained (bottom 10%: mean phyloP = 3.67) comparable to large positive effect functional mutations for the most LoF-intolerant genes (top 10%: mean phyloP = 3.88). Thus, even LoF-tolerant genes show constraint against likely deleterious promoter mutations, a finding consistent across alternative measures of gene constraint, including LOEUF, MOEUF, and AlphaMissense ^124,126^ (**Supplementary Fig. 29)**.

## Discussion

Identifying causal non-coding variants is challenging due to the complex grammar underlying CRE function. MPRAs have shown promise in quantitatively measuring non-coding variants’ allelic skew to identify those likely underlying human traits and disease; however, oligo-based assays can only assay a maximum of a few hundred thousand variants at a time ^66^. To address this, we developed MPAC, an extension of our previous Malinois model of MPRA activity ^78^. MPAC provides a scalable framework for genome-wide non-coding variant effect prediction, and requires significantly less training data than other models. We leveraged MPAC’s scalability, which exceeds the largest empirical MPRAs by orders of magnitude^100^, to predict non-coding functional effects of large databases of hundreds of millions of common, rare, *de novo*, and somatic variation genome-wide, and to perform deep mutational scans across all human promoters.

While we demonstrate MPAC’s functional predictions reliably identify clinically relevant non-coding variation, there are important biological and methodological limitations to consider. First, our predictions are based on an episomal MPRA using a minimal promoter designed to interrogate CRE function that is best suited for a subset of non-coding sequences ^66,84^. MPAC is unable to predict functional variants in UTRs, some repressors, large structural or copy number variants, or elements involved in chromatin looping. Reporter assays are also unable to link CREs with their target genes, requiring methods such as pooled CRISPRi screens or Hi-C to contextualize the effect of the CRE. From a modeling perspective, MPAC is trained on MPRA data from three cell lines which limits the TFs identified by the model to those expressed in K562, HepG2, and SK-N-SH. We also identify a small number of motifs that contribute to missed emVar predictions, which could be tested via an active learning approach to improve model accuracy. We expect that advances in pre-training, multi-modal model architectures, expanded MPRA training size ^76,85^ and cellular contexts will improve the model, especially improving cell type-specific, non-promoter regulatory elements.

As genomic medicine advances towards personalized treatments, the availability of predictions for every possible mutation across the human genome will have many applications. For example, deep mutational scanning has demonstrated clinical utility in coding regions ^127–131^, yet the scale and complexity of the non-coding genome makes regulatory variant effect modeling difficult. We expect MPAC predictions to complement existing maps of CREs, providing nucleotide-level predictions for functionally informed GWAS fine-mapping and rare disease clinical interpretation ^111,132,133^. We functionally nominate a set of 1,892 recurrent somatic, putatively tumorigenic mutations in COSMIC for deeper mechanistic study. Our comprehensive analysis of gnomAD SNVs quantifies the evolutionary forces acting on regulatory function and enables better clinical interpretation of rare, likely deleterious, non-coding variants. Lastly, our promoter saturation mutagenesis map facilitates exploration of the relationships between constraint and function at individual genes and equips disease researchers with functional predictions for all possible promoter SNVs. MPAC predictions are rapidly deployable on DNA sequence alone, ancestry-agnostic, independent of linkage disequilibrium, and do not rely on large sequencing cohorts to identify putatively causal functional variants.

Overall, the advent of highly performant non-coding variant effect predictors like MPAC to create prospective maps of variant function at the scale of the entire genome will enable genome interpretation to match the speed of variant discovery. As whole-genome sequencing becomes increasingly available across common, rare, and somatic disease cohorts, the need for effective variant interpretation will only grow. While MPAC focuses on non-coding regulatory variation, we expect its integration with advanced models of other genomic functions ^52,55,81,82,112,134–139^ will enhance personal genome interpretation and deepen our molecular understanding of human disease.

## Data availability

All MPAC predictions, additional datasets and model hyperparameters are available at Zenodo (https://doi.org/10.5281/zenodo.15178434) organized as follows:

Supplementary Dataset 1: MPAC_UKBB_BBJ_GTEx_MPRA_predictions

Supplementary Dataset 2: MPAC_DNase_ASE_predictions

Supplementary Dataset 3: MPAC_ClinVar_predictions

Supplementary Dataset 4: MPAC_COSMIC_predictions

Supplementary Dataset 5: MPAC_gnomAD_predictions

Supplementary Dataset 6: MPAC_GWAS_tag_SNVs+LD_predictions

Supplementary Dataset 7: MPAC_promoter_saturation_mutagenesis

Supplementary Dataset 8: Promoter_function_constraint_correlations

Supplementary Dataset 9: Promoter_gene_set_enrichment_results

Supplementary Dataset 10: MPAC_model_hyperparameters

Supplementary Dataset 11: MPAC_model_artifacts

## Code availability

All code and documentation are deposited online as follows. Code used for MPAC variant effect predictions and to analyze and visualize data related to the benchmarking, ClinVar, and COSMIC work is available at https://github.com/john-c-butts/MPAC/. Code used to analyze and visualize data related to the gnomAD and promoter saturation mutagenesis work is available at https://github.com/Reilly-Lab-Yale/MPAC_gnomAD_and_satmut.

## Contributions

J.C.B., S.R., S.J.G., R.I.C., R.T., and S.K.R. designed the study and contributed to the development of MPAC. J.C.B., S.R., S.J.G., M.N., K.A., R.G., R.I.C., R.T., and S.K.R. performed data analysis and interpreted results. J.C.B., S.R., M.N., R.T., and S.K.R. drafted the manuscript. P.C.S., R.T., and S.K.R. secured funding. R.T. and S.K.R. supervised the study. All the authors revised the manuscript and accepted its final version.

## Acknowledgements

We thank Jared Akers, Erin Gilbertson, Ira Hall, Takeshi Iwasaki, Diyendo Massilani, Catherine McGuinness, Kousuke Mouri, Niketa Nerurkar, Thanh Nguyen, Arya Rao, and the IGVF Evolution group for advice and comments during the preparation of this manuscript. We thank J. Ulirsch for advice, input on design of the study, and unpublished results. We thank Jeff Vierstra and members of his lab for sharing unpublished DNase ASE data. SR was supported by the Boehringer Ingelheim Biomedical Data Science Fellowship. MN was supported by a Gruber Foundation Fellowship. This work was directly supported by National Institutes of Health grants UM1HG009435, R00HG008179, R00HG010669, R01HG012872, and R35HG011329. The content is solely the responsibility of the authors and does not necessarily represent the official views of the National Institutes of Health.

## Methods

### Data preprocessing and model training

We partitioned a previously generated MPRA dataset ^66^ into 11 roughly even sized splits based on the chromosome pairs from which the oligo sequences were derived. Pairs of source chromosomes were selected such that the sums of the chromosome identifiers were 23 (e.g., chr3 and chr20 form a pair). Next, we organized the data into 110 combinations of training, validation, and test sequences composed of 10, 1, and 1 pairs of source chromosomes, respectively. The training and validation groups were used to fit and select model parameters, respectively. Therefore, for every pair of source chromosomes, 10 models are fitted using unique training/validation combinations to use for benchmarking performance and making predictions. We then trained 110 models as previously described using these data set combinations using the code base available at https://github.com/sjgosai/boda2 and hyperparameter settings defined in (**Supplementary Data 10**).

### Predicting variant effects using MPAC

Command line tools, the original Malinois model, and code base is available at (https://github.com/sjgosai/boda2). The 110 MPAC models are available at Zenodo (https://doi.org/10.5281/zenodo.15178434). Predictions are generated with the ‘vcf_predict.py’ script as shown below:

> *python /opt/boda2/src/vcf_predict.py --artifact_path {10X $MODEL} --vcf_file ${VCF} --fasta_file ${FASTA} –output ${OUTPUT} --relative_start 9 --relative_end 181 --step_size 10 --strand_reduction mean --window_reduction mean*

The above example generates 18 predictions per variant, tested in positions 9 to 181 (0-based, 10 bp intervals) which are reduced to a single reference and alternate activity (log_2_FC) by averaging. Allelic skew is then calculated as the difference between alternate and reference activity. GRCh38_no_alt_analysis_set_GCA_000001405.15.fasta was downloaded from https://www.encodeproject.org/files/GRCh38_no_alt_analysis_set_GCA_000001405.15/ and used as the reference genome for all analyses. All predictions were generated with the above prediction schema except for correlation with empirical MPRA activity (**Figure 1B**) which used the following:

> *python /opt/boda2/src/vcf_predict.py --artifact_path {10X $MODEL} --vcf_file ${VCF} --fasta_file ${FASTA} –output ${OUTPUT} --relative_start 99 --relative_end 100 --step_size 1 --strand_reduction mean --window_reduction mean*

Which generates a single prediction with the variant centered at position 99 (0-based) to exactly reproduce the sequences tested by the empirical MPRA used for comparison ^66^.

### MPAC correlation with empirical MPRA

Single, variant centered predictions were generated for all empirical MPRA sequences from ^66^ that passed QC, K562 (n=485,180), HepG2 (n=499,820), and SK-N-SH (n=485,034). Empirical log_2_FC was compared to predicted log_2_FC for computing Pearson’s correlation. Next, MPAC MPAC allelic skew and empirical Log2Skew were compared for empirically validated emVars ((logPadj_BF ≳ −log10(0.01) & abs(log2FC) >= 1) & Skew_logPadj >= −log10(0.1)) in K562 (*n* = 16,187), HepG2 (*n* = 14,029), and SK-N-SH (*n* = 14,723) from the same experimental dataset. Predictions and empirical measurements were correlated using Pearson’s method.

### Correlation to DNase allele-specific effects variants

DNase allele specific effects (ASE) data were obtained for the three MPAC cell types (K562, HepG2, and SK-N-SH) from Jeff Vierstra (unpublished). Significant ASE variants were called using a minimum FDR threshold of <= 0.05. As SK-N-SH samples had no significant hits they were dropped from subsequent analysis. ASE mean effect sizes were multiplied by −1 to match directionality of MPAC allelic skew (Alt – Ref). MPAC allelic skew was compared to DNase skew for all K562 and HepG2 ASE variants as well as the subset of MPAC emVars (|allelic skew| > 0.5). Significant ASE variants were identified in previously performed MPRA experiments ^66^ and Log2Skew was compared to ASE effect size for both all ASE variants and those called emVars by MPRA ((logPadj_BF ≳ −log10(0.01) & abs(log2FC) >= 1) & Skew_logPadj >= −log10(0.1)). All correlations were performed using Pearson’s method.

For additional model comparisons precomputed Enformer 1KG predictions were downloaded from (https://console.cloud.google.com/storage/browser/dm-enformer/variant-scores/1000-genomes/enformer) and filtered for significant ASE variants (HepG2 *n* = 331, K562: *n* = 855). All Pearson’s correlations presented in **Fig. 1D** reflect this subset. Sei ^81^ predictions were generated for these same variants using the web interface (https://hb.flatironinstitute.org/deepsea/) and filtered for HepG2 and K562 DNase. For both Enformer and Sei the highest Pearson’s correlation to empirical DNase was reported.

### Correlation of Enformer and Sei DNase predictions to MPRA emVars

All empirical emVars ^66^ present in the precomputed Enformer 1KG variants predictions (HepG2 *n* = 10,407, K562 = 11,993) were downloaded from (https://console.cloud.google.com/storage/browser/dm-enformer/variant-scores/1000-genomes/enformer). Sei was downloaded and installed from (https://github.com/FunctionLab/sei-framework). Variant effect predictions for all emVars were generated using the following command ‘sh 1_variant_effect_prediction.sh../vcfs2sei/emvar_vcf.vcf hg38 ../sei_preds/mpra_emvars --cuda.’ For both Enformer and Sei predictions all HepG2 and K562 DNase predictions were compared to MPRA skew and the highest Pearson’s correlation was used.

### Precision-Recall of high PIP GWAS variants

We generated MPAC predictions for a set of statistically fine-mapped variants from the UK Biobank and Biobank Japan previously tested by empirical MPRA ^66^. After filtering for autosomal variants tested in K562, HepG2, or SK-N-SH, the test set contained 1,869 variants: 934 positive (PIP > 0.9) and 935 negative (PIP < 0.01). MPAC predictions were reduced to a single score using the max(|allelic skew|) of K562, HepG2, or SK-N-SH. We also generated predictions for these variants with Sei ^81^ and LS-GKM ^86,140^. Sei predictions were generated with the following command ‘sh 1_variant_effect_prediction.sh ../vcfs2sei/traits_prc_hg38_vars_for_SEI_preds.vcf hg38 ../sei_preds/ukbb_gtex --cuda’ and the max(|allelic skew|) prediction for DNase in all cell types was used. LS-GKM and model weights were downloaded from (https://www.beerlab.org/deltasvm/). HepG2 predictions were generated using the following command ‘perl deltasvm.pl ref_fasta alt_fasta SupplementaryTable_hepg2weights.txt hepg2_out’ and K562 predictions with ‘perl deltasvm.pl ref_fasta alt_fasta SupplementaryTable_k562weights.txt k562_out’. The max(|allelic skew|) prediction from K562 and HepG2 was used. Precision and recall curves, as well as AUPRC measurements were generated using scikit-learn 1.4.1.post1 ^141^.

### Precision-Recall of high PIP GTEx variants

MPAC predictions were generated for a set of previously described high (> 0.9) and low (< 0.01) PIP eQTLs ^66^. After filtering for autosomal variants empirically tested in K562, HepG2, and SK-N-SH, SNVs the test set contained 22,720 variants: 11,354 positive and 11,366 negative. MPAC predictions were reduced to a single score using the max(|allelic skew|) of K562, HepG2, and SK-N-SH. Sei ^81^ predictions were generated with ‘sh 1_variant_effect_prediction.sh {../vcfs2sei/gtex_prc_hg38_vars_for_SEI_preds.vcf} {hg38} {../sei_preds/ukbb_gtex} --cuda’ and the max(|allelic skew|) prediction for DNase in all cell types was used. LS-GKM ^86,140^ predictions were generated using the command ‘perl deltasvm.pl {ref_fasta} {alt_fasta} {SupplementaryTable_hepg2weights.txt} {hepg2_out}’ for HepG2 and ‘perl deltasvm.pl {ref_fasta} {alt_fasta} {SupplementaryTable_k562weights.txt} {k562_out}’ for K562.The max(|allelic skew|) prediction from K562 and HepG2 was used. As before LS-GKM and model weights were downloaded from (https://www.beerlab.org/deltasvm/). Precision and recall curves, and AUPRC measurements were generated using scikit-learn 1.4.1.post1 ^141^.

### Missed MPAC prediction analysis

To understand sources of MPAC missed predictions compared to empirical MPRA, we subset all MPRA emVars for those with an |allelic skew| > 0.5 and an MPAC predicted |allelic skew| < 0.05. These incorrectly predicted emVars were independently identified from each cell type, split into 80/20 training and test sets and matched with a randomly sampled set of correctly predicted sequences. An LS-GKM model ^86,140^ was trained for each cell type using the command ‘gkmtrain {positive_train_fasta} {negative_train_fasta} {output}.’

Using these models, importance scores were generated using gkmexplain ^88^ and passed to tfmodisco-lite (https://github.com/jmschrei/tfmodisco-lite) ^87^ for de-novo motif identification. tfmodisco-lite motifs were matched to known TF motifs using TOMTOM (MEME version 5.5.7, HOCOMOCOv11_core_HUMAN) ^142^. We next used FIMO ^143^ (‘fimo –no-pgc -o {outfile} {modisc_pwms} {emvar_fasta}’) to quantify the presence of motifs identified by tfmodisco-lite in correctly predicted emVar sequences. FIMO hits were considered if they overlapped the variant (oligo position 100) and had a q-value < 0.01. Lastly, to determine abundance of de-novo motifs in the correctly predicted emVars the total number of significant FIMO hits with a significant TOMTOM match (q-value < 0.05) to a known motif was divided by the total number of correctly predicted emVar sequences.

### ClinVar analysis

The ClinVar VCF was downloaded from (https://ftp.ncbi.nlm.nih.gov/pub/clinvar/vcf_GRCh38/archive_2.0/2023/clinvar_20230930.vcf.gz, fileDate: 2023-09-30) and MPAC predictions were generated for all ClinVar variants prior to filtering for non-coding variants. To focus on non-coding SNVs, we filtered transcripts from GENCODE v44 basic annotations ^144^ (downloaded from https://www.gencodegenes.org/human/release_44.html) that were located on chr1-22, had a transcript_type column = “protein_coding”, a non-missing hgnc_id, and tag column matching “Ensembl_canonical” but not matching “readthrough_gene”. We then removed variants intersecting any exons. To remove potential splice disrupting variations, we also removed SNVs in intronic regions up to −20 from 3’ splice sites or +6 from 5’ splice sites, consistent with splice region definitions from MaxEntScan ^144,145^. After filtering a total of 180,032 variants classified as pathogenic (ClinVar ‘Pathogenic’ or ‘Likely_pathogenic’) or benign (ClinVar ‘Benign’ or ‘Likely_benign’) were used for all downstream analyses. ClinVar variants were intersected with peaks of DNase I Hypersensitivity (DHS) downloaded from (https://www.encodeproject.org/files/ENCFF503GCK/) ^146^ and promoters. Promoters were defined as 250 bp immediately upstream of the TSS of filtered GENCODE transcripts. emVar proportions for pathogenic and benign variants are calculated as the number of emVars labeled non-DHS, DHS, or Promoter over the total number of variants in each category. Significance was tested by the Chi-Square Test. Enrichments for pathogenic and benign variants were calculated by odds ratio of emVars in each respective group and tested by Fisher’s exact test to calculate *p* values.

Sei ^81^ predictions were generated for all non-coding SNVs using the command ‘sh 1_variant_effect_prediction.sh {clinvar_exon_filtered_vars.vcf} {hg38} {/clinvar_preds} --cuda’ and filtered for K562, HepG2, and SK-N-SH ENCODE DHS predictions. For a given category (All ClinVar, DHS, Promoter) all pathogenic variants were compared to an equal random sample of benign variants 100 times. For both MPAC and Sei predictions the max(|allelic skew|) prediction from K562, HepG2, and SK-N-SH was used in precision recall calculations. Precision and recall curves, and AUPRC measures were generated using scikit-learn 1.4.1.post1 ^141^.

### COSMIC analysis

MPAC predictions were generated for all non-coding COSMIC ^23^ (v98, downloaded from (https://cancer.sanger.ac.uk/cosmic/download/cosmic/v98/noncodingvariantsvcf). Variants were filtered for only SNVs derived from whole genome sequencing (’Whole_Genome_Reseq’ == ‘y’ and ‘Whole_Exome’ == ‘n’) from (https://cancer.sanger.ac.uk/cosmic/download/cosmic/v98/noncodingvariantstsv). Variant recurrence was calculated as the number of COSMIC mutation IDs (’GENOMIC_MUTATION_ID’) present after filtering. MPAC predictions were generated for ETS-TF disrupting putative drivers previously tested by MPRA ^98^. We calculated percent emVar recovery as the number of MPAC emVars divided by empirically defined emVars.

Variants were subset into recurrent (*n* ≥ 2) or unique (*n* = 1) by ID count and feature annotations, promoters (as defined in ClinVar analysis), cancer promoters (CNC) ^99^, recurrent promoters, and recurrent CNC promoters were assigned by intersection. CNC promoters were downloaded from (https://cncdatabase.med.cornell.edu/) with the query ‘promoters’ and using all unique gene names for the final list. Variants were categorized into putative regulatory elements by intersection with DHS elements from (https://www.encodeproject.org/files/ENCFF503GCK/) ^146^. Odds ratios (OR) were calculated for the proportion of emVars in the category of interest versus the remainder of COSMIC variants. For allelic skew bin comparisons bins were defined as the mean allelic skew of all predictions. *OR*s were calculated for promoter variants in the allelic skew bin versus distal variants. For activity comparisons variants were binned by the max(ref | alt) log_2_FC and the *OR* that a recurrent or unique variant was an emVar was calculated. For all *OR*s Fisher’s exact test was used to calculate *p* values and 95% confidence intervals.

### MPAC gnomAD SNV predictions and analysis

We used MPAC to predict variant effects for all 646,033,065 SNVs reported in 76,156 whole genomes analyzed by gnomAD v3.1.2 on hg38 ^29^ (downloaded from https://gnomad.broadinstitute.org/downloads#v3-variants). We filtered to include SNVs that: 1) passed gnomAD QC, 2) were located on autosomes (chr1-22), 3) had non-zero minor allele frequency (MAF = min(AF, 1 - AF) > 0), 4) non-zero allele count (AC > 0), and 5) total number of alleles observed is at least half of gnomAD (AN ≥ 76,156).

We annotated SNVs by 1) overlapping Zoonomia 241-way base-level phyloP scores ^26^ (downloaded from https://cgl.gi.ucsc.edu/data/cactus/241-mammalian-2020v2-hub/Homo_sapiens/241-mammalian-2020v2.bigWig), and 2) annotations of mutation rate from Roulette predictions ^147^ (as annotated in CADD v1.6 ^111^ downloaded from https://krishna.gs.washington.edu/download/CADD/v1.6/GRCh38/whole_genome_SNVs.tsv.gz). We removed variants without a phyloP annotation (1.7% of variants).

To focus on non-coding SNVs, we removed variants intersecting our previously defined exons and splice regions on filtered GENCODE transcripts (see “*ClinVar analysis*”). This resulted in a final set of 513,886,257 SNVs for downstream analyses.

GWAS Catalog (accessed 2024-12-19) ^10^ SNVs were expanded to include all variants in high LD (*r*^2^ ≥ 0.7) in each of three superpopulations (ASN, AFR, EUR). This set was assigned MPAC predictions by subsetting our filtered GWAS prediction set to those with a MAF ≥ 1%, then joining by rsID.

### gnomAD SNV functional annotations

We categorized non-coding SNVs by intersecting with the ENCODE cCRE class annotations ^37,144,145^ (downloaded from https://downloads.wenglab.org/Registry-V4/GRCh38-cCREs.bed), specifically promoter-like signatures (PLS), proximal enhancer-like signatures (pELS), and distal enhancer-like signatures (dELS). SNVs not falling into the above categories were further classified as “Other cCREs” if they overlapped any remaining cCRE annotations, or labeled as “non-cCRE” if overlapping no annotations.

For the TF binding enrichment analysis, we labeled SNVs as overlapping DNase I TF footprints if they intersected at least one consensus footprint from Vierstra et al. ^104^ (downloaded from https://zenodo.org/records/3905306) and overlapping TF ChIP-seq peaks if they intersected at least one peak from ChIP-Atlas ^105,106^. Odds ratios (*OR*) were calculated using Fisher’s exact test with a +1 pseudocount correction to calculate the *OR*, *p*-value, and 95% CI.

To compare mean phyloP between non-coding and coding functional variants, we annotated SNVs by Ensembl VEP ^105,148^ based on gnomAD v3.1.2 annotations using the worst predicted consequence as reported in the sequence ontology ranking provided in the Ensembl VEP documentation (https://ensembl.org/info/genome/variation/prediction/predicted_data.html retrieved 2024-06-10) prior to removing SNVs overlapping exons or splice regions.

### gnomAD evolutionary constraint analyses

We binned SNVs by their MPAC predicted mean log_2_FC in activity across all cell lines or mean log_2_FC allelic skew. We used the enrichment of rare (AF < 0.1%) vs. common (AF ≥ 0.1%) SNVs as a measure of recent purifying selection and compared across MPAC allelic-skew bin for each ENCODE cCRE class. We omitted bins with less than 50 variants. We calculated the odds ratios (*OR*) based on counts of rare vs. common SNVs belonging to a focal subset of SNVs (MPAC allelic-skew bin x ENCODE cCRE class) relative to that of all other SNVs in the genome, using Fisher’s exact test with a +1 pseudocount correction to calculate the *OR*, *p*-value, and 95% CI.

### MPAC promoter saturation mutagenesis predictions

Starting from the filtered list of 18,761 GENCODE transcripts (see “*ClinVar analysis*”), we used MPAC to conduct *in silico* saturation mutagenesis of all 3,000 possible single nucleotide mutations across the 1 kb promoter regions located upstream of the TSS. We masked mutations intersecting any exons or splice regions of filtered GENCODE transcripts (see “*ClinVar analysis*”), resulting in 18,658 promoter regions with any mutations after masking. At each base position, the mean activity (log_2_FC) or allelic skew (log_2_FC) is that of all three non-reference mutations averaged across all three cell lines (nine values total). The 1 kb promoter regions were used for **Supplementary Fig. 21** and **22**. All subsequent analyses were conducted on the 250 bp promoter region upstream of the TSS. Means at each distance from the TSS were calculated using promoters that had unmasked values at that specific position.

### Motif analysis of promoter regions

DNA sequences for 250 bp promoter regions were scanned for TF binding motifs using FIMO from the memes R package ^143,149^ and position weight matrices from JASPAR 2024 ^150^ with accessions: TATA (MA0108.3), SP1 (MA0079.5), NRF1 (MA0506.1), ELK1 (MA0028.2), NFYC (MA1644.1), CREB1 (MA0018.3), YY1 (MA0095.2), CTCF (MA0139.1), and ZNF143 (MA0088.2). Significant motifs were called at default *p* < 4 x 10^−4^, which corresponds to rescaling the default FIMO value of 1 x 10^−4^ to 250 bp promoters as specified in https://meme-suite.org/meme/doc/fimo-tutorial.html. Motif count profiles were based on the positions of motif midpoints relative to the TSS.

### Gene-level expression annotations

Gene expression in transcripts per million (TPM) were downloaded from the Human Protein Atlas for K562, HepG2, and SK-N-SH cell lines ^151^. Using Pearson’s *r*, the log_10_TPMs were then correlated against the mean MPAC-predicted activity of each 250 bp promoter region, computed as the mean of activities centered on every position in the region. Gene expression annotations were joined to filtered GENCODE transcripts (see “*ClinVar analysis*”) by matching Ensembl gene IDs, Ensembl transcript IDs, or gene names.

### Gene-level coding constraint stratification

Promoters were annotated by four different scores of gene-level coding constraint: GeneBayes *s_het_*LoF-intolerance ^125^, LOEUF LoF-intolerance and MOEUF missense-intolerance from gnomAD v4.0 ^124^, and mean AlphaMissense missense pathogenicity ^126^. Each of these were binned into deciles for the coding constraint stratification analyses. Gene-level coding constraint annotations were joined to filtered GENCODE transcripts (see “*ClinVar analysis*”) by matching Ensembl gene IDs, Ensembl transcript IDs, or gene names.

### Promoter correlations between function and constraint

For each promoter, we calculated Spearman correlation (*ρ*) between MPAC-predicted allelic skew at each position (mean log_2_FC of the three possible mutations across three cell lines) and across-species base-level constraint (max(0, phyloP) scores). We performed this calculation separately for negative or positive allelic-skew variants, requiring 50 unmasked positions that had both a phyloP score and a non-zero skew prediction. Among analyzed promoters, 17,392 had defined correlations for at least one category (positive or negative allelic-skew variants), while 16,995 had both. We identified promoters with significant correlations by comparing them to a permutation-based null distribution created by shuffling base positions among 250 bp promoter regions, where we defined promoters as significant if their correlation values fell below the bottom 0.01% (for negative allelic-skew) or above the top 0.01% (for positive allelic-skew) of this null distribution.

Gene set enrichment analyses were conducted using the Enrichr web server ^152^ with the negative allelic-skew correlation, positive allelic-skew correlation, or both correlation gene sets using the background set of 17,392 genes where either the positive or negative allelic-skew correlations was well-defined, focusing on Reactome Pathways 2024, GO Biological Process 2025, and MGI Mammalian Phenotype Level 4 gene sets with FDR < 0.05.

## References

1. Phan, L., Zhang, H., Wang, Q., Villamarin, R., Hefferon, T., Ramanathan, A. & Kattman, B. The evolution of dbSNP: 25 years of impact in genomic research. Nucleic Acids Res. 53, D925–D931 (2025).

2. Saunders, G., Baudis, M., Becker, R., Beltran, S., Béroud, C., Birney, E., Brooksbank, C., Brunak, S., Van den Bulcke, M., Drysdale, R., Capella-Gutierrez, S., Flicek, P., Florindi, F., Goodhand, P., Gut, I., Heringa, J., Holub, P., Hooyberghs, J., Juty, N., Keane, T. M., Korbel, J. O., Lappalainen, I., Leskosek, B., Matthijs, G., Mayrhofer, M. T., Metspalu, A., Navarro, A., Newhouse, S., Nyrönen, T., Page, A., Persson, B., Palotie, A., Parkinson, H., Rambla, J., Salgado, D., Steinfelder, E., Swertz, M. A., Valencia, A., Varma, S., Blomberg, N. & Scollen, S. Leveraging European infrastructures to access 1 million human genomes by 2022. Nat Rev Genet 20, 693–701 (2019).

3. Gallagher, C. S., Ginsburg, G. S. & Musick, A. Biobanking with genetics shapes precision medicine and global health. Nat Rev Genet 26, 191–202 (2025).

4. Genomic data in the All of Us Research Program. Nature 627, 340–346 (2024).

5. Hindorff, L. A., Sethupathy, P., Junkins, H. A., Ramos, E. M., Mehta, J. P., Collins, F. S. & Manolio, T. A. Potential etiologic and functional implications of genome-wide association loci for human diseases and traits. Proc. Natl. Acad. Sci. U. S. A. 106, 9362–9367 (2009).

6. Maurano, M. T., Humbert, R., Rynes, E., Thurman, R. E., Haugen, E., Wang, H., Reynolds, A. P., Sandstrom, R., Qu, H., Brody, J., Shafer, A., Neri, F., Lee, K., Kutyavin, T., Stehling-Sun, S., Johnson, A. K., Canfield, T. K., Giste, E., Diegel, M., Bates, D., Hansen, R. S., Neph, S., Sabo, P. J., Heimfeld, S., Raubitschek, A., Ziegler, S., Cotsapas, C., Sotoodehnia, N., Glass, I., Sunyaev, S. R., Kaul, R. & Stamatoyannopoulos, J. A. Systematic localization of common disease-associated variation in regulatory DNA. Science 337, 1190–1195 (2012).

7. ENCODE Project Consortium. An integrated encyclopedia of DNA elements in the human genome. Nature 489, 57–74 (2012).

8. Gallagher, M. D. & Chen-Plotkin, A. S. The Post-GWAS Era: From Association to Function. Am J Hum Genet 102, 717–730 (2018).

9. Lappalainen, T. & MacArthur, D. G. From variant to function in human disease genetics. Science 373, 1464–1468 (2021).

10. Sollis, E., Mosaku, A., Abid, A., Buniello, A., Cerezo, M., Gil, L., Groza, T., Güneş, O., Hall, P., Hayhurst, J., Ibrahim, A., Ji, Y., John, S., Lewis, E., MacArthur, J. A. L., McMahon, A., Osumi-Sutherland, D., Panoutsopoulou, K., Pendlington, Z., Ramachandran, S., Stefancsik, R., Stewart, J., Whetzel, P., Wilson, R., Hindorff, L., Cunningham, F., Lambert, S. A., Inouye, M., Parkinson, H. & Harris, L. W. The NHGRI-EBI GWAS Catalog: knowledgebase and deposition resource. Nucleic Acids Res. 51, D977–D985 (2023).

11. Gazal, S., Loh, P.-R., Finucane, H. K., Ganna, A., Schoech, A., Sunyaev, S. & Price, A. L. Functional architecture of low-frequency variants highlights strength of negative selection across coding and non-coding annotations. Nat. Genet. 50, 1600–1607 (2018).

12. Wang, Q., Dhindsa, R. S., Carss, K., Harper, A. R., Nag, A., Tachmazidou, I., Vitsios, D., Deevi, S. V. V., Mackay, A., Muthas, D., Hühn, M., Monkley, S., Olsson, H., AstraZeneca Genomics Initiative, Wasilewski, S., Smith, K. R., March, R., Platt, A., Haefliger, C. & Petrovski, S. Rare variant contribution to human disease in 281,104 UK Biobank exomes. Nature 597, 527–532 (2021).

13. Hawkes, G., Beaumont, R. N., Li, Z., Mandla, R., Li, X., Albert, C. M., Arnett, D. K., Ashley-Koch, A. E., Ashrani, A. A., Barnes, K. C., Boerwinkle, E., Brody, J. A., Carson, A. P., Chami, N., Chen, Y.-D. I., Chung, M. K., Curran, J. E., Darbar, D., Ellinor, P. T., Fornage, M., Gordeuk, V. R., Guo, X., He, J., Hwu, C.-M., Kalyani, R. R., Kaplan, R., Kardia, S. L. R., Kooperberg, C., Loos, R. J. F., Lubitz, S. A., Minster, R. L., Naseri, T., Viali, S. ‘itea, Mitchell, B. D., Murabito, J. M., Palmer, N. D., Psaty, B. M., Redline, S., Shoemaker, M. B., Silverman, E. K., Telen, M. J., Weiss, S. T., Yanek, L. R., Zhou, H., NHLBI Trans-Omics for Precision Medicine (TOPMed) Consortium, Liu, C.-T., North, K. E., Justice, A. E., Locke, J. M., Owens, N., Murray, A., Patel, K., Frayling, T. M., Wright, C. F., Wood, A. R., Lin, X., Manning, A. & Weedon, M. N. Whole-genome sequencing in 333,100 individuals reveals rare non-coding single variant and aggregate associations with height. Nat. Commun. 15, 8549 (2024).

14. Hawkes, G., Chundru, K., Jackson, L., Patel, K. A., Murray, A., Wood, A. R., Wright, C. F., Weedon, M. N., Frayling, T. M. & Beaumont, R. N. Whole-genome sequencing analysis identifies rare, large-effect noncoding variants and regulatory regions associated with circulating protein levels. Nat. Genet. (2025). doi:10.1038/s41588-025-02095-4

15. Weiner, D. J., Nadig, A., Jagadeesh, K. A., Dey, K. K., Neale, B. M., Robinson, E. B., Karczewski, K. J. & O’Connor, L. J. Polygenic architecture of rare coding variation across 394,783 exomes. Nature 614, 492–499 (2023).

16. Samocha, K. E., Robinson, E. B., Sanders, S. J., Stevens, C., Sabo, A., McGrath, L. M., Kosmicki, J. A., Rehnström, K., Mallick, S., Kirby, A., Wall, D. P., MacArthur, D. G., Gabriel, S. B., DePristo, M., Purcell, S. M., Palotie, A., Boerwinkle, E., Buxbaum, J. D., Cook, E. H., Jr, Gibbs, R. A., Schellenberg, G. D., Sutcliffe, J. S., Devlin, B., Roeder, K., Neale, B. M. & Daly, M. J. A framework for the interpretation of de novo mutation in human disease. Nat. Genet. 46, 944–950 (2014).

17. Turner, T. N., Coe, B. P., Dickel, D. E., Hoekzema, K., Nelson, B. J., Zody, M. C., Kronenberg, Z. N., Hormozdiari, F., Raja, A., Pennacchio, L. A., Darnell, R. B. & Eichler, E. E. Genomic patterns of DE Novo mutation in simplex autism. Cell 171, 710–722.e12 (2017).

18. Short, P. J., McRae, J. F., Gallone, G., Sifrim, A., Won, H., Geschwind, D. H., Wright, C. F., Firth, H. V., FitzPatrick, D. R., Barrett, J. C. & Hurles, M. E. De novo mutations in regulatory elements in neurodevelopmental disorders. Nature 555, 611–616 (2018).

19. Wright, C. F., Campbell, P., Eberhardt, R. Y., Aitken, S., Perrett, D., Brent, S., Danecek, P., Gardner, E. J., Chundru, V. K., Lindsay, S. J., Andrews, K., Hampstead, J., Kaplanis, J., Samocha, K. E., Middleton, A., Foreman, J., Hobson, R. J., Parker, M. J., Martin, H. C., FitzPatrick, D. R., Hurles, M. E., Firth, H. V. & DDD Study. Genomic diagnosis of rare pediatric disease in the United Kingdom and Ireland. N. Engl. J. Med. 388, 1559–1571 (2023).

20. McConnell, M. J., Moran, J. V., Abyzov, A., Akbarian, S., Bae, T., Cortes-Ciriano, I., Erwin, J. A., Fasching, L., Flasch, D. A., Freed, D., Ganz, J., Jaffe, A. E., Kwan, K. Y., Kwon, M., Lodato, M. A., Mills, R. E., Paquola, A. C. M., Rodin, R. E., Rosenbluh, C., Sestan, N., Sherman, M. A., Shin, J. H., Song, S., Straub, R. E., Thorpe, J., Weinberger, D. R., Urban, A. E., Zhou, B., Gage, F. H., Lehner, T., Senthil, G., Walsh, C. A., Chess, A., Courchesne, E., Gleeson, J. G., Kidd, J. M., Park, P. J., Pevsner, J., Vaccarino, F. M. & Brain Somatic Mosaicism Network. Intersection of diverse neuronal genomes and neuropsychiatric disease: The Brain Somatic Mosaicism Network. Science 356, (2017).

21. Maury, E. A., Jones, A., Seplyarskiy, V., Nguyen, T. T. L., Rosenbluh, C., Bae, T., Wang, Y., Abyzov, A., Khoshkhoo, S., Chahine, Y., Zhao, S., Venkatesh, S., Root, E., Voloudakis, G., Roussos, P., Brain Somatic Mosaicism Network‡, Park, P. J., Akbarian, S., Brennand, K., Reilly, S., Lee, E. A., Sunyaev, S. R., Walsh, C. A. & Chess, A. Somatic mosaicism in schizophrenia brains reveals prenatal mutational processes. Science 386, 217–224 (2024).

22. Rheinbay, E., Nielsen, M. M., Abascal, F., Wala, J. A., Shapira, O., Tiao, G., Hornshøj, H., Hess, J. M., Juul, R. I., Lin, Z., Feuerbach, L., Sabarinathan, R., Madsen, T., Kim, J., Mularoni, L., Shuai, S., Lanzós, A., Herrmann, C., Maruvka, Y. E., Shen, C., Amin, S. B., Bandopadhayay, P., Bertl, J., Boroevich, K. A., Busanovich, J., Carlevaro-Fita, J., Chakravarty, D., Chan, C. W. Y., Craft, D., Dhingra, P., Diamanti, K., Fonseca, N. A., Gonzalez-Perez, A., Guo, Q., Hamilton, M. P., Haradhvala, N. J., Hong, C., Isaev, K., Johnson, T. A., Juul, M., Kahles, A., Kahraman, A., Kim, Y., Komorowski, J., Kumar, K., Kumar, S., Lee, D., Lehmann, K.-V., Li, Y., Liu, E. M., Lochovsky, L., Park, K., Pich, O., Roberts, N. D., Saksena, G., Schumacher, S. E., Sidiropoulos, N., Sieverling, L., Sinnott-Armstrong, N., Stewart, C., Tamborero, D., Tubio, J. M. C., Umer, H. M., Uusküla-Reimand, L., Wadelius, C., Wadi, L., Yao, X., Zhang, C.-Z., Zhang, J., Haber, J. E., Hobolth, A., Imielinski, M., Kellis, M., Lawrence, M. S., von Mering, C., Nakagawa, H., Raphael, B. J., Rubin, M. A., Sander, C., Stein, L. D., Stuart, J. M., Tsunoda, T., Wheeler, D. A., Johnson, R., Reimand, J., Gerstein, M., Khurana, E., Campbell, P. J., López-Bigas, N., Weischenfeldt, J., Beroukhim, R., Martincorena, I., Pedersen, J. S. & Getz, G. Analyses of non-coding somatic drivers in 2,658 cancer whole genomes. Nature 578, 102–111 (2020).

23. Sondka, Z., Dhir, N. B., Carvalho-Silva, D., Jupe, S., Madhumita, McLaren, K., Starkey, M., Ward, S., Wilding, J., Ahmed, M., Argasinska, J., Beare, D., Chawla, M. S., Duke, S., Fasanella, I., Neogi, A. G., Haller, S., Hetenyi, B., Hodges, L., Holmes, A., Lyne, R., Maurel, T., Nair, S., Pedro, H., Sangrador-Vegas, A., Schuilenburg, H., Sheard, Z., Yong, S. Y. & Teague, J. COSMIC: a curated database of somatic variants and clinical data for cancer. Nucleic Acids Res. 52, D1210–D1217 (2024).

24. Pollard, K. S., Hubisz, M. J., Rosenbloom, K. R. & Siepel, A. Detection of nonneutral substitution rates on mammalian phylogenies. Genome Res. 20, 110–121 (2010).

25. Zoonomia Consortium. A comparative genomics multitool for scientific discovery and conservation. Nature 587, 240–245 (2020).

26. Christmas, M. J., Kaplow, I. M., Genereux, D. P., Dong, M. X., Hughes, G. M., Li, X., Sullivan, P. F., Hindle, A. G., Andrews, G., Armstrong, J. C., Bianchi, M., Breit, A. M., Diekhans, M., Fanter, C., Foley, N. M., Goodman, D. B., Goodman, L., Keough, K. C., Kirilenko, B., Kowalczyk, A., Lawless, C., Lind, A. L., Meadows, J. R. S., Moreira, L. R., Redlich, R. W., Ryan, L., Swofford, R., Valenzuela, A., Wagner, F., Wallerman, O., Brown, A. R., Damas, J., Fan, K., Gatesy, J., Grimshaw, J., Johnson, J., Kozyrev, S. V., Lawler, A. J., Marinescu, V. D., Morrill, K. M., Osmanski, A., Paulat, N. S., Phan, B. N., Reilly, S. K., Schäffer, D. E., Steiner, C., Supple, M. A., Wilder, A. P., Wirthlin, M. E., Xue, J. R., Zoonomia Consortium§, Birren, B. W., Gazal, S., Hubley, R. M., Koepfli, K.-P., Marques-Bonet, T., Meyer, W. K., Nweeia, M., Sabeti, P. C., Shapiro, B., Smit, A. F. A., Springer, M. S., Teeling, E. C., Weng, Z., Hiller, M., Levesque, D. L., Lewin, H. A., Murphy, W. J., Navarro, A., Paten, B., Pollard, K. S., Ray, D. A., Ruf, I., Ryder, O. A., Pfenning, A. R., Lindblad-Toh, K. & Karlsson, E. K. Evolutionary constraint and innovation across hundreds of placental mammals. Science 380, eabn3943 (2023).

27. Sullivan, P. F., Meadows, J. R. S., Gazal, S., Phan, B. N., Li, X., Genereux, D. P., Dong, M. X., Bianchi, M., Andrews, G., Sakthikumar, S., Nordin, J., Roy, A., Christmas, M. J., Marinescu, V. D., Wang, C., Wallerman, O., Xue, J., Yao, S., Sun, Q., Szatkiewicz, J., Wen, J., Huckins, L. M., Lawler, A., Keough, K. C., Zheng, Z., Zeng, J., Wray, N. R., Li, Y., Johnson, J., Chen, J., Zoonomia Consortium§, Paten, B., Reilly, S. K., Hughes, G. M., Weng, Z., Pollard, K. S., Pfenning, A. R., Forsberg-Nilsson, K., Karlsson, E. K. & Lindblad-Toh, K. Leveraging base-pair mammalian constraint to understand genetic variation and human disease. Science 380, eabn2937 (2023).

28. Kuderna, L. F. K., Ulirsch, J. C., Rashid, S., Ameen, M., Sundaram, L., Hickey, G., Cox, A. J., Gao, H., Kumar, A., Aguet, F., Christmas, M. J., Clawson, H., Haeussler, M., Janiak, M. C., Kuhlwilm, M., Orkin, J. D., Bataillon, T., Manu, S., Valenzuela, A., Bergman, J., Rouselle, M., Silva, F. E., Agueda, L., Blanc, J., Gut, M., de Vries, D., Goodhead, I., Harris, R. A., Raveendran, M., Jensen, A., Chuma, I. S., Horvath, J. E., Hvilsom, C., Juan, D., Frandsen, P., Schraiber, J. G., de Melo, F. R., Bertuol, F., Byrne, H., Sampaio, I., Farias, I., Valsecchi, J., Messias, M., da Silva, M. N. F., Trivedi, M., Rossi, R., Hrbek, T., Andriaholinirina, N., Rabarivola, C. J., Zaramody, A., Jolly, C. J., Phillips-Conroy, J., Wilkerson, G., Abee, C., Simmons, J. H., Fernandez-Duque, E., Kanthaswamy, S., Shiferaw, F., Wu, D., Zhou, L., Shao, Y., Zhang, G., Keyyu, J. D., Knauf, S., Le, M. D., Lizano, E., Merker, S., Navarro, A., Nadler, T., Khor, C. C., Lee, J., Tan, P., Lim, W. K., Kitchener, A. C., Zinner, D., Gut, I., Melin, A. D., Guschanski, K., Schierup, M. H., Beck, R. M. D., Karakikes, I., Wang, K. C., Umapathy, G., Roos, C., Boubli, J. P., Siepel, A., Kundaje, A., Paten, B., Lindblad-Toh, K., Rogers, J., Marques Bonet, T. & Farh, K. K.-H. Identification of constrained sequence elements across 239 primate genomes. Nature 625, 735–742 (2024).

29. Chen, S., Francioli, L. C., Goodrich, J. K., Collins, R. L., Kanai, M., Wang, Q., Alföldi, J., Watts, N. A., Vittal, C., Gauthier, L. D., Poterba, T., Wilson, M. W., Tarasova, Y., Phu, W., Grant, R., Yohannes, M. T., Koenig, Z., Farjoun, Y., Banks, E., Donnelly, S., Gabriel, S., Gupta, N., Ferriera, S., Tolonen, C., Novod, S., Bergelson, L., Roazen, D., Ruano-Rubio, V., Covarrubias, M., Llanwarne, C., Petrillo, N., Wade, G., Jeandet, T., Munshi, R., Tibbetts, K., Genome Aggregation Database Consortium, O’Donnell-Luria, A., Solomonson, M., Seed, C., Martin, A. R., Talkowski, M. E., Rehm, H. L., Daly, M. J., Tiao, G., Neale, B. M., MacArthur, D. G. & Karczewski, K. J. A genomic mutational constraint map using variation in 76,156 human genomes. Nature 625, 92–100 (2024).

30. Wittkopp, P. J. & Kalay, G. Cis-regulatory elements: molecular mechanisms and evolutionary processes underlying divergence. Nat. Rev. Genet. 13, 59–69 (2011).

31. Preissl, S., Gaulton, K. J. & Ren, B. Characterizing cis-regulatory elements using single-cell epigenomics. Nat. Rev. Genet. 24, 21–43 (2023).

32. Yáñez-Cuna, J. O., Kvon, E. Z. & Stark, A. Deciphering the transcriptional cis-regulatory code. Trends Genet. 29, 11–22 (2013).

33. Zeitlinger, J. Seven myths of how transcription factors read the cis-regulatory code. Curr. Opin. Syst. Biol. 23, 22–31 (2020).

34. de Boer, C. G. & Taipale, J. Hold out the genome: a roadmap to solving the cis-regulatory code. Nature 625, 41–50 (2024).

35. Kim, S. & Wysocka, J. Deciphering the multi-scale, quantitative cis-regulatory code. Mol. Cell 83, 373–392 (2023).

36. Finucane, H. K., Bulik-Sullivan, B., Gusev, A., Trynka, G., Reshef, Y., Loh, P.-R., Anttila, V., Xu, H., Zang, C., Farh, K., Ripke, S., Day, F. R., ReproGen Consortium, Schizophrenia Working Group of the Psychiatric Genomics Consortium, RACI Consortium, Purcell, S., Stahl, E., Lindstrom, S., Perry, J. R. B., Okada, Y., Raychaudhuri, S., Daly, M. J., Patterson, N., Neale, B. M. & Price, A. L. Partitioning heritability by functional annotation using genome-wide association summary statistics. Nat. Genet. 47, 1228–1235 (2015).

37. ENCODE Project Consortium, Moore, J. E., Purcaro, M. J., Pratt, H. E., Epstein, C. B., Shoresh, N., Adrian, J., Kawli, T., Davis, C. A., Dobin, A., Kaul, R., Halow, J., Van Nostrand, E. L., Freese, P., Gorkin, D. U., Shen, Y., He, Y., Mackiewicz, M., Pauli-Behn, F., Williams, B. A., Mortazavi, A., Keller, C. A., Zhang, X.-O., Elhajjajy, S. I., Huey, J., Dickel, D. E., Snetkova, V., Wei, X., Wang, X., Rivera-Mulia, J. C., Rozowsky, J., Zhang, J., Chhetri, S. B., Zhang, J., Victorsen, A., White, K. P., Visel, A., Yeo, G. W., Burge, C. B., Lécuyer, E., Gilbert, D. M., Dekker, J., Rinn, J., Mendenhall, E. M., Ecker, J. R., Kellis, M., Klein, R. J., Noble, W. S., Kundaje, A., Guigó, R., Farnham, P. J., Cherry, J. M., Myers, R. M., Ren, B., Graveley, B. R., Gerstein, M. B., Pennacchio, L. A., Snyder, M. P., Bernstein, B. E., Wold, B., Hardison, R. C., Gingeras, T. R., Stamatoyannopoulos, J. A. & Weng, Z. Expanded encyclopaedias of DNA elements in the human and mouse genomes. Nature 583, 699–710 (2020).

38. Andersson, R., Gebhard, C., Miguel-Escalada, I., Hoof, I., Bornholdt, J., Boyd, M., Chen, Y., Zhao, X., Schmidl, C., Suzuki, T., Ntini, E., Arner, E., Valen, E., Li, K., Schwarzfischer, L., Glatz, D., Raithel, J., Lilje, B., Rapin, N., Bagger, F. O., Jørgensen, M., Andersen, P. R., Bertin, N., Rackham, O., Burroughs, A. M., Baillie, J. K., Ishizu, Y., Shimizu, Y., Furuhata, E., Maeda, S., Negishi, Y., Mungall, C. J., Meehan, T. F., Lassmann, T., Itoh, M., Kawaji, H., Kondo, N., Kawai, J., Lennartsson, A., Daub, C. O., Heutink, P., Hume, D. A., Jensen, T. H., Suzuki, H., Hayashizaki, Y., Müller, F., Forrest, A. R. R., Carninci, P., Rehli, M. & Sandelin, A. An atlas of active enhancers across human cell types and tissues. Nature 507, 455–461 (2014).

39. Roadmap Epigenomics Consortium, Kundaje, A., Meuleman, W., Ernst, J., Bilenky, M., Yen, A., Heravi-Moussavi, A., Kheradpour, P., Zhang, Z., Wang, J., Ziller, M. J., Amin, V., Whitaker, J. W., Schultz, M. D., Ward, L. D., Sarkar, A., Quon, G., Sandstrom, R. S., Eaton, M. L., Wu, Y.-C., Pfenning, A. R., Wang, X., Claussnitzer, M., Liu, Y., Coarfa, C., Harris, R. A., Shoresh, N., Epstein, C. B., Gjoneska, E., Leung, D., Xie, W., Hawkins, R. D., Lister, R., Hong, C., Gascard, P., Mungall, A. J., Moore, R., Chuah, E., Tam, A., Canfield, T. K., Hansen, R. S., Kaul, R., Sabo, P. J., Bansal, M. S., Carles, A., Dixon, J. R., Farh, K.-H., Feizi, S., Karlic, R., Kim, A.-R., Kulkarni, A., Li, D., Lowdon, R., Elliott, G., Mercer, T. R., Neph, S. J., Onuchic, V., Polak, P., Rajagopal, N., Ray, P., Sallari, R. C., Siebenthall, K. T., Sinnott-Armstrong, N. A., Stevens, M., Thurman, R. E., Wu, J., Zhang, B., Zhou, X., Beaudet, A. E., Boyer, L. A., De Jager, P. L., Farnham, P. J., Fisher, S. J., Haussler, D., Jones, S. J. M., Li, W., Marra, M. A., McManus, M. T., Sunyaev, S., Thomson, J. A., Tlsty, T. D., Tsai, L.-H., Wang, W., Waterland, R. A., Zhang, M. Q., Chadwick, L. H., Bernstein, B. E., Costello, J. F., Ecker, J. R., Hirst, M., Meissner, A., Milosavljevic, A., Ren, B., Stamatoyannopoulos, J. A., Wang, T. & Kellis, M. Integrative analysis of 111 reference human epigenomes. Nature 518, 317–330 (2015).

40. Stunnenberg, H. G., International Human Epigenome Consortium & Hirst, M. The International Human Epigenome Consortium: A blueprint for scientific collaboration and discovery. Cell 167, 1145–1149 (2016).

41. Zhang, K., Hocker, J. D., Miller, M., Hou, X., Chiou, J., Poirion, O. B., Qiu, Y., Li, Y. E., Gaulton, K. J., Wang, A., Preissl, S. & Ren, B. A single-cell atlas of chromatin accessibility in the human genome. Cell 184, 5985–6001.e19 (2021).

42. Boix, C. A., James, B. T., Park, Y. P., Meuleman, W. & Kellis, M. Regulatory genomic circuitry of human disease loci by integrative epigenomics. Nature 590, 300–307 (2021).

43. Göring, H. H. H., Curran, J. E., Johnson, M. P., Dyer, T. D., Charlesworth, J., Cole, S. A., Jowett, J. B. M., Abraham, L. J., Rainwater, D. L., Comuzzie, A. G., Mahaney, M. C., Almasy, L., MacCluer, J. W., Kissebah, A. H., Collier, G. R., Moses, E. K. & Blangero, J. Discovery of expression QTLs using large-scale transcriptional profiling in human lymphocytes. Nat. Genet. 39, 1208–1216 (2007).

44. Degner, J. F., Pai, A. A., Pique-Regi, R., Veyrieras, J.-B., Gaffney, D. J., Pickrell, J. K., De Leon, S., Michelini, K., Lewellen, N., Crawford, G. E., Stephens, M., Gilad, Y. & Pritchard, J. K. DNase I sensitivity QTLs are a major determinant of human expression variation. Nature 482, 390–394 (2012).

45. van der Wijst, M. G. P., Brugge, H., de Vries, D. H., Deelen, P., Swertz, M. A., LifeLines Cohort Study, BIOS Consortium & Franke, L. Single-cell RNA sequencing identifies celltype-specific cis-eQTLs and co-expression QTLs. Nat. Genet. 50, 493–497 (2018).

46. GTEx Consortium. The GTEx Consortium atlas of genetic regulatory effects across human tissues. Science 369, 1318–1330 (2020).

47. Võsa, U., Claringbould, A., Westra, H.-J., Bonder, M. J., Deelen, P., Zeng, B., Kirsten, H., Saha, A., Kreuzhuber, R., Yazar, S., Brugge, H., Oelen, R., de Vries, D. H., van der Wijst, M. G. P., Kasela, S., Pervjakova, N., Alves, I., Favé, M.-J., Agbessi, M., Christiansen, M. W., Jansen, R., Seppälä, I., Tong, L., Teumer, A., Schramm, K., Hemani, G., Verlouw, J., Yaghootkar, H., Sönmez Flitman, R., Brown, A., Kukushkina, V., Kalnapenkis, A., Rüeger, S., Porcu, E., Kronberg, J., Kettunen, J., Lee, B., Zhang, F., Qi, T., Hernandez, J. A., Arindrarto, W., Beutner, F., BIOS Consortium, i2QTL Consortium, Dmitrieva, J., Elansary, M., Fairfax, B. P., Georges, M., Heijmans, B. T., Hewitt, A. W., Kähönen, M., Kim, Y., Knight, J. C., Kovacs, P., Krohn, K., Li, S., Loeffler, M., Marigorta, U. M., Mei, H., Momozawa, Y., Müller-Nurasyid, M., Nauck, M., Nivard, M. G., Penninx, B. W. J. H., Pritchard, J. K., Raitakari, O. T., Rotzschke, O., Slagboom, E. P., Stehouwer, C. D. A., Stumvoll, M., Sullivan, P., ’t Hoen, P. A. C., Thiery, J., Tönjes, A., van Dongen, J., van Iterson, M., Veldink, J. H., Völker, U., Warmerdam, R., Wijmenga, C., Swertz, M., Andiappan, A., Montgomery, G. W., Ripatti, S., Perola, M., Kutalik, Z., Dermitzakis, E., Bergmann, S., Frayling, T., van Meurs, J., Prokisch, H., Ahsan, H., Pierce, B. L., Lehtimäki, T., Boomsma, D. I., Psaty, B. M., Gharib, S. A., Awadalla, P., Milani, L., Ouwehand, W. H., Downes, K., Stegle, O., Battle, A., Visscher, P. M., Yang, J., Scholz, M., Powell, J., Gibson, G., Esko, T. & Franke, L. Large-scale cis- and trans-eQTL analyses identify thousands of genetic loci and polygenic scores that regulate blood gene expression. Nat. Genet. 53, 1300–1310 (2021).

48. Yazar, S., Alquicira-Hernandez, J., Wing, K., Senabouth, A., Gordon, M. G., Andersen, S., Lu, Q., Rowson, A., Taylor, T. R. P., Clarke, L., Maccora, K., Chen, C., Cook, A. L., Ye, C. J., Fairfax, K. A., Hewitt, A. W. & Powell, J. E. Single-cell eQTL mapping identifies cell type-specific genetic control of autoimmune disease. Science 376, eabf3041 (2022).

49. Rosenberg, A. B., Patwardhan, R. P., Shendure, J. & Seelig, G. Learning the sequence determinants of alternative splicing from millions of random sequences. Cell 163, 698–711 (2015).

50. Bogard, N., Linder, J., Rosenberg, A. B. & Seelig, G. A Deep Neural Network for Predicting and Engineering Alternative Polyadenylation. Cell 178, 91–106.e23 (2019).

51. Sample, P. J., Wang, B., Reid, D. W., Presnyak, V., McFadyen, I. J., Morris, D. R. & Seelig, G. Human 5′ UTR design and variant effect prediction from a massively parallel translation assay. Nat. Biotechnol. 37, 803–809 (2019).

52. Kelley, D. R., Reshef, Y. A., Bileschi, M., Belanger, D., McLean, C. Y. & Snoek, J. Sequential regulatory activity prediction across chromosomes with convolutional neural networks. Genome Res. 28, 739–750 (2018).

53. Zhou, J. & Troyanskaya, O. G. Predicting effects of noncoding variants with deep learning-based sequence model. Nat. Methods 12, 931–934 (2015).

54. Quang, D. & Xie, X. DanQ: a hybrid convolutional and recurrent deep neural network for quantifying the function of DNA sequences. Nucleic Acids Res. 44, e107 (2016).

55. Jaganathan, K., Kyriazopoulou Panagiotopoulou, S., McRae, J. F., Darbandi, S. F., Knowles, D., Li, Y. I., Kosmicki, J. A., Arbelaez, J., Cui, W., Schwartz, G. B., Chow, E. D., Kanterakis, E., Gao, H., Kia, A., Batzoglou, S., Sanders, S. J. & Farh, K. K.-H. Predicting Splicing from Primary Sequence with Deep Learning. Cell 176, 535–548.e24 (2019).

56. de Almeida, B. P., Reiter, F., Pagani, M. & Stark, A. DeepSTARR predicts enhancer activity from DNA sequence and enables the de novo design of synthetic enhancers. Nat. Genet. 54, 613–624 (2022).

57. Penzar, D., Nogina, D., Noskova, E., Zinkevich, A., Meshcheryakov, G., Lando, A., Rafi, A. M., de Boer, C. & Kulakovskiy, I. V. LegNet: a best-in-class deep learning model for short DNA regulatory regions. Bioinformatics 39, (2023).

58. Melnikov, A., Murugan, A., Zhang, X., Tesileanu, T., Wang, L., Rogov, P., Feizi, S., Gnirke, A., Callan, C. G., Jr, Kinney, J. B., Kellis, M., Lander, E. S. & Mikkelsen, T. S. Systematic dissection and optimization of inducible enhancers in human cells using a massively parallel reporter assay. Nat. Biotechnol. 30, 271–277 (2012).

59. Patwardhan, R. P., Hiatt, J. B., Witten, D. M., Kim, M. J., Smith, R. P., May, D., Lee, C., Andrie, J. M., Lee, S.-I., Cooper, G. M., Ahituv, N., Pennacchio, L. A. & Shendure, J. Massively parallel functional dissection of mammalian enhancers in vivo. Nat. Biotechnol. 30, 265–270 (2012).

60. Arnold, C. D., Gerlach, D., Stelzer, C., Boryń, Ł. M., Rath, M. & Stark, A. Genome-wide quantitative enhancer activity maps identified by STARR-seq. Science 339, 1074–1077 (2013).

61. Tewhey, R., Kotliar, D., Park, D. S., Liu, B., Winnicki, S., Reilly, S. K., Andersen, K. G., Mikkelsen, T. S., Lander, E. S., Schaffner, S. F. & Sabeti, P. C. Direct Identification of Hundreds of Expression-Modulating Variants using a Multiplexed Reporter Assay. Cell 165, 1519–1529 (2016).

62. Ulirsch, J. C., Nandakumar, S. K., Wang, L., Giani, F. C., Zhang, X., Rogov, P., Melnikov, A., McDonel, P., Do, R., Mikkelsen, T. S. & Sankaran, V. G. Systematic functional dissection of common genetic variation affecting red blood cell traits. Cell 165, 1530–1545 (2016).

63. Abell, N. S., DeGorter, M. K., Gloudemans, M. J., Greenwald, E., Smith, K. S., He, Z. & Montgomery, S. B. Multiple causal variants underlie genetic associations in humans. Science 375, 1247–1254 (2022).

64. McAfee, J. C., Lee, S., Lee, J., Bell, J. L., Krupa, O., Davis, J., Insigne, K., Bond, M. L., Zhao, N., Boyle, A. P., Phanstiel, D. H., Love, M. I., Stein, J. L., Ruzicka, W. B., Davila-Velderrain, J., Kosuri, S. & Won, H. Systematic investigation of allelic regulatory activity of schizophrenia-associated common variants. Cell Genom. 3, 100404 (2023).

65. Deng, C., Whalen, S., Steyert, M., Ziffra, R., Przytycki, P. F., Inoue, F., Pereira, D. A., Capauto, D., Norton, S., Vaccarino, F. M., PsychENCODE Consortium‡, Pollen, A. A., Nowakowski, T. J., Ahituv, N., Pollard, K. S. & PsychENCODE Consortium. Massively parallel characterization of regulatory elements in the developing human cortex. Science 384, eadh0559 (2024).

66. Siraj, L., Castro, R. I., Dewey, H., Kales, S., Nguyen, T. T. L., Kanai, M., Berenzy, D., Mouri, K., Wang, Q., McCaw, Z. R., Gosai, S. J., Aguet, F., Cui, R., Vockley, C. M., Lareau, C. A., Okada, Y., Gusev, A., Jones, T. R., Lander, E. S., Sabeti, P. C., Finucane, H. K., Reilly, S. K., Ulirsch, J. C. & Tewhey, R. Functional dissection of complex and molecular trait variants at single nucleotide resolution. bioRxiv 2024.05.05.592437 (2024). doi:10.1101/2024.05.05.592437

67. Feng, Y., Xie, N., Inoue, F., Fan, S., Saskin, J., Zhang, C., Zhang, F., Hansen, M. E. B., Nyambo, T., Mpoloka, S. W., Mokone, G. G., Fokunang, C., Belay, G., Njamnshi, A. K., Marks, M. S., Oancea, E., Ahituv, N. & Tishkoff, S. A. Integrative functional genomic analyses identify genetic variants influencing skin pigmentation in Africans. Nat. Genet. 56, 258–272 (2024).

68. Shin, T., Song, J. H. T., Kosicki, M., Kenny, C., Beck, S. G., Kelley, L., Antony, I., Qian, X., Bonacina, J., Papandile, F., Gonzalez, D., Scotellaro, J., Bushinsky, E. M., Andersen, R. E., Maury, E., Pennacchio, L. A., Doan, R. N. & Walsh, C. A. Rare variation in non-coding regions with evolutionary signatures contributes to autism spectrum disorder risk. Cell Genom. 4, 100609 (2024).

69. Uebbing, S., Gockley, J., Reilly, S. K., Kocher, A. A., Geller, E., Gandotra, N., Scharfe, C., Cotney, J. & Noonan, J. P. Massively parallel discovery of human-specific substitutions that alter enhancer activity. Proc. Natl. Acad. Sci. U. S. A. 118, e2007049118 (2021).

70. Weiss, C. V., Harshman, L., Inoue, F., Fraser, H. B., Petrov, D. A., Ahituv, N. & Gokhman, D. The cis-regulatory effects of modern human-specific variants. Elife 10, (2021).

71. Jagoda, E., Xue, J. R., Reilly, S. K., Dannemann, M., Racimo, F., Huerta-Sanchez, E., Sankararaman, S., Kelso, J., Pagani, L., Sabeti, P. C. & Capellini, T. D. Detection of Neanderthal adaptively introgressed genetic variants that modulate reporter gene expression in human immune cells. Mol. Biol. Evol. 39, (2022).

72. Mangan, R. J., Alsina, F. C., Mosti, F., Sotelo-Fonseca, J. E., Snellings, D. A., Au, E. H., Carvalho, J., Sathyan, L., Johnson, G. D., Reddy, T. E., Silver, D. L. & Lowe, C. B. Adaptive sequence divergence forged new neurodevelopmental enhancers in humans. Cell 185, 4587–4603.e23 (2022).

73. Xue, J. R., Mackay-Smith, A., Mouri, K., Garcia, M. F., Dong, M. X., Akers, J. F., Noble, M., Li, X., Zoonomia Consortium†, Lindblad-Toh, K., Karlsson, E. K., Noonan, J. P., Capellini, T. D., Brennand, K. J., Tewhey, R., Sabeti, P. C. & Reilly, S. K. The functional and evolutionary impacts of human-specific deletions in conserved elements. Science 380, eabn2253 (2023).

74. Sasse, A., Chikina, M. & Mostafavi, S. Unlocking gene regulation with sequence-to-function models. Nat. Methods 21, 1374–1377 (2024).

75. Movva, R., Greenside, P., Marinov, G. K., Nair, S., Shrikumar, A. & Kundaje, A. Deciphering regulatory DNA sequences and noncoding genetic variants using neural network models of massively parallel reporter assays. PLoS One 14, e0218073 (2019).

76. Vaishnav, E. D., de Boer, C. G., Molinet, J., Yassour, M., Fan, L., Adiconis, X., Thompson, D. A., Levin, J. Z., Cubillos, F. A. & Regev, A. The evolution, evolvability and engineering of gene regulatory DNA. Nature 603, 455–463 (2022).

77. Tareen, A., Kooshkbaghi, M., Posfai, A., Ireland, W. T., McCandlish, D. M. & Kinney, J. B. MAVE-NN: learning genotype-phenotype maps from multiplex assays of variant effect. Genome Biol. 23, 98 (2022).

78. Gosai, S. J., Castro, R. I., Fuentes, N., Butts, J. C., Mouri, K., Alasoadura, M., Kales, S., Nguyen, T. T. L., Noche, R. R., Rao, A. S., Joy, M. T., Sabeti, P. C., Reilly, S. K. & Tewhey, R. Machine-guided design of cell-type-targeting cis-regulatory elements. Nature 634, 1211–1220 (2024).

79. Rafi, A. M., Nogina, D., Penzar, D., Lee, D., Lee, D., Kim, N., Kim, S., Kim, D., Shin, Y., Kwak, I.-Y., Meshcheryakov, G., Lando, A., Zinkevich, A., Kim, B.-C., Lee, J., Kang, T., Vaishnav, E. D., Yadollahpour, P., Random Promoter DREAM Challenge Consortium, Kim, S., Albrecht, J., Regev, A., Gong, W., Kulakovskiy, I. V., Meyer, P. & de Boer, C. G. A community effort to optimize sequence-based deep learning models of gene regulation. Nat. Biotechnol. (2024). doi:10.1038/s41587-024-02414-w

80. Agarwal, V., Inoue, F., Schubach, M., Penzar, D., Martin, B. K., Dash, P. M., Keukeleire, P., Zhang, Z., Sohota, A., Zhao, J., Georgakopoulos-Soares, I., Noble, W. S., Yardımcı, G. G., Kulakovskiy, I. V., Kircher, M., Shendure, J. & Ahituv, N. Massively parallel characterization of transcriptional regulatory elements. Nature 1–10 (2025).

81. Chen, K. M., Wong, A. K., Troyanskaya, O. G. & Zhou, J. A sequence-based global map of regulatory activity for deciphering human genetics. Nat Genet 54, 940–949 (2022).

82. Avsec, Ž., Agarwal, V., Visentin, D., Ledsam, J. R., Grabska-Barwinska, A., Taylor, K. R., Assael, Y., Jumper, J., Kohli, P. & Kelley, D. R. Effective gene expression prediction from sequence by integrating long-range interactions. Nat Methods 18, 1196–1203 (2021).

83. Lee, D., Gorkin, D. U., Baker, M., Strober, B. J., Asoni, A. L., McCallion, A. S. & Beer, M. A. A method to predict the impact of regulatory variants from DNA sequence. Nat Genet 47, 955–961 (2015).

84. Klein, J. C., Agarwal, V., Inoue, F., Keith, A., Martin, B., Kircher, M., Ahituv, N. & Shendure, J. A systematic evaluation of the design and context dependencies of massively parallel reporter assays. Nat. Methods 17, 1083–1091 (2020).

85. Sahu, B., Hartonen, T., Pihlajamaa, P., Wei, B., Dave, K., Zhu, F., Kaasinen, E., Lidschreiber, K., Lidschreiber, M., Daub, C. O., Cramer, P., Kivioja, T. & Taipale, J. Sequence determinants of human gene regulatory elements. Nat. Genet. 54, 283–294 (2022).

86. Ghandi, M., Lee, D., Mohammad-Noori, M. & Beer, M. A. Enhanced Regulatory Sequence Prediction Using Gapped k-mer Features. PLOS Computational Biology 10, e1003711 (2014).

87. Shrikumar, A., Tian, K., Avsec, Ž., Shcherbina, A., Banerjee, A., Sharmin, M., Nair, S. & Kundaje, A. Technical note on Transcription Factor Motif Discovery from Importance Scores (TF-MoDISco) version 0.5.6.5. arXiv [cs.LG] (2018). at <http://arxiv.org/abs/1811.00416>

88. Shrikumar, A., Prakash, E. & Kundaje, A. GkmExplain: fast and accurate interpretation of nonlinear gapped k-mer SVMs. Bioinformatics 35, i173–i182 (2019).

89. Kribelbauer-Swietek, J. F., Gardeux, V., Llimos-Aubach, G., Faltejskova, K., Russeil, J., Grenningloh, N., Levassor, L., Steiner, C., Vondrasek, J. & Deplancke, B. EXTRA-seq: a genome-integrated extended massively parallel reporter assay to quantify enhancer-promoter communication. bioRxiv 2024.12.08.627402 (2024). doi:10.1101/2024.12.08.627402

90. Landrum, M. J., Lee, J. M., Riley, G. R., Jang, W., Rubinstein, W. S., Church, D. M. & Maglott, D. R. ClinVar: public archive of relationships among sequence variation and human phenotype. Nucleic Acids Res. 42, D980–5 (2014).

91. Rehm, H. L., Berg, J. S., Brooks, L. D., Bustamante, C. D., Evans, J. P., Landrum, M. J., Ledbetter, D. H., Maglott, D. R., Martin, C. L., Nussbaum, R. L., Plon, S. E., Ramos, E. M., Sherry, S. T., Watson, M. S. & ClinGen. ClinGen--the clinical genome resource. N. Engl. J. Med. 372, 2235–2242 (2015).

92. ICGC/TCGA Pan-Cancer Analysis of Whole Genomes Consortium. Pan-cancer analysis of whole genomes. Nature 578, 82–93 (2020).

93. Fredriksson, N. J., Ny, L., Nilsson, J. A. & Larsson, E. Systematic analysis of noncoding somatic mutations and gene expression alterations across 14 tumor types. Nat. Genet. 46, 1258–1263 (2014).

94. Wang, Y. & Hon, G. C. Towards functional maps of non-coding variants in cancer. Front. Genome Ed. 6, 1481443 (2024).

95. Huang, F. W., Hodis, E., Xu, M. J., Kryukov, G. V., Chin, L. & Garraway, L. A. Highly recurrent TERT promoter mutations in human melanoma. Science 339, 957–959 (2013).

96. Horn, S., Figl, A., Rachakonda, P. S., Fischer, C., Sucker, A., Gast, A., Kadel, S., Moll, I., Nagore, E., Hemminki, K., Schadendorf, D. & Kumar, R. TERT promoter mutations in familial and sporadic melanoma. Science 339, 959–961 (2013).

97. Dratwa, M., Wysoczańska, B., Łacina, P., Kubik, T. & Bogunia-Kubik, K. TERT-Regulation and Roles in Cancer Formation. Front. Immunol. 11, 589929 (2020).

98. Carrasco Pro, S., Hook, H., Bray, D., Berenzy, D., Moyer, D., Yin, M., Labadorf, A. T., Tewhey, R., Siggers, T. & Fuxman Bass, J. I. Widespread perturbation of ETS factor binding sites in cancer. Nat. Commun. 14, 913 (2023).

99. Liu, E. M., Martinez-Fundichely, A., Bollapragada, R., Spiewack, M. & Khurana, E. CNCDatabase: a database of non-coding cancer drivers. Nucleic Acids Res. 49, D1094–D1101 (2021).

100. van Arensbergen, J., Pagie, L., FitzPatrick, V. D., de Haas, M., Baltissen, M. P., Comoglio, F., van der Weide, R. H., Teunissen, H., Võsa, U., Franke, L., de Wit, E., Vermeulen, M., Bussemaker, H. J. & van Steensel, B. High-throughput identification of human SNPs affecting regulatory element activity. Nat. Genet. 51, 1160–1169 (2019).

101. Visel, A., Blow, M. J., Li, Z., Zhang, T., Akiyama, J. A., Holt, A., Plajzer-Frick, I., Shoukry, M., Wright, C., Chen, F., Afzal, V., Ren, B., Rubin, E. M. & Pennacchio, L. A. ChIP-seq accurately predicts tissue-specific activity of enhancers. Nature 457, 854–858 (2009).

102. Heintzman, N. D., Hon, G. C., Hawkins, R. D., Kheradpour, P., Stark, A., Harp, L. F., Ye, Z., Lee, L. K., Stuart, R. K., Ching, C. W., Ching, K. A., Antosiewicz-Bourget, J. E., Liu, H., Zhang, X., Green, R. D., Lobanenkov, V. V., Stewart, R., Thomson, J. A., Crawford, G. E., Kellis, M. & Ren, B. Histone modifications at human enhancers reflect global cell-type-specific gene expression. Nature 459, 108–112 (2009).

103. Szutorisz, H., Dillon, N. & Tora, L. The role of enhancers as centres for general transcription factor recruitment. Trends Biochem. Sci. 30, 593–599 (2005).

104. Vierstra, J., Lazar, J., Sandstrom, R., Halow, J., Lee, K., Bates, D., Diegel, M., Dunn, D., Neri, F., Haugen, E., Rynes, E., Reynolds, A., Nelson, J., Johnson, A., Frerker, M., Buckley, M., Kaul, R., Meuleman, W. & Stamatoyannopoulos, J. A. Global reference mapping of human transcription factor footprints. Nature 583, 729–736 (2020).

105. Zou, Z., Ohta, T., Miura, F. & Oki, S. ChIP-Atlas 2021 update: a data-mining suite for exploring epigenomic landscapes by fully integrating ChIP-seq, ATAC-seq and Bisulfite-seq data. Nucleic Acids Res 50, W175–W182 (2022).

106. Oki, S., Ohta, T., Shioi, G., Hatanaka, H., Ogasawara, O., Okuda, Y., Kawaji, H., Nakaki, R., Sese, J. & Meno, C. ChIP-Atlas: a data-mining suite powered by full integration of public ChIP-seq data. EMBO Rep 19, (2018).

107. Andolfatto, P. Adaptive evolution of non-coding DNA in Drosophila. Nature 437, 1149–1152 (2005).

108. Ward, L. D. & Kellis, M. Evidence of abundant purifying selection in humans for recently acquired regulatory functions. Science 337, 1675–1678 (2012).

109. Khurana, E., Fu, Y., Colonna, V., Mu, X. J., Kang, H. M., Lappalainen, T., Sboner, A., Lochovsky, L., Chen, J., Harmanci, A., Das, J., Abyzov, A., Balasubramanian, S., Beal, K., Chakravarty, D., Challis, D., Chen, Y., Clarke, D., Clarke, L., Cunningham, F., Evani, U. S., Flicek, P., Fragoza, R., Garrison, E., Gibbs, R., Gümüş, Z. H., Herrero, J., Kitabayashi, N., Kong, Y., Lage, K., Liluashvili, V., Lipkin, S. M., MacArthur, D. G., Marth, G., Muzny, D., Pers, T. H., Ritchie, G. R. S., Rosenfeld, J. A., Sisu, C., Wei, X., Wilson, M., Xue, Y., Yu, F., 1000 Genomes Project Consortium, Dermitzakis, E. T., Yu, H., Rubin, M. A., Tyler-Smith, C. & Gerstein, M. Integrative annotation of variants from 1092 humans: application to cancer genomics. Science 342, 1235587 (2013).

110. Mouri, K., Dewey, H. B., Castro, R., Berenzy, D., Kales, S. & Tewhey, R. Whole-genome functional characterization of RE1 silencers using a modified massively parallel reporter assay. Cell Genom. 3, 100234 (2023).

111. Kircher, M., Witten, D. M., Jain, P., O’Roak, B. J., Cooper, G. M. & Shendure, J. A general framework for estimating the relative pathogenicity of human genetic variants. Nat Genet 46, 310–315 (2014).

112. Schubach, M., Maass, T., Nazaretyan, L., Röner, S. & Kircher, M. CADD v1.7: using protein language models, regulatory CNNs and other nucleotide-level scores to improve genome-wide variant predictions. Nucleic Acids Res 52, D1143–D1154 (2024).

113. Margulies, E. H., Cooper, G. M., Asimenos, G., Thomas, D. J., Dewey, C. N., Siepel, A., Birney, E., Keefe, D., Schwartz, A. S., Hou, M., Taylor, J., Nikolaev, S., Montoya-Burgos, J. I., Löytynoja, A., Whelan, S., Pardi, F., Massingham, T., Brown, J. B., Bickel, P., Holmes, I., Mullikin, J. C., Ureta-Vidal, A., Paten, B., Stone, E. A., Rosenbloom, K. R., Kent, W. J., Bouffard, G. G., Guan, X., Hansen, N. F., Idol, J. R., Maduro, V. V. B., Maskeri, B., McDowell, J. C., Park, M., Thomas, P. J., Young, A. C., Blakesley, R. W., Muzny, D. M., Sodergren, E., Wheeler, D. A., Worley, K. C., Jiang, H., Weinstock, G. M., Gibbs, R. A., Graves, T., Fulton, R., Mardis, E. R., Wilson, R. K., Clamp, M., Cuff, J., Gnerre, S., Jaffe, D. B., Chang, J. L., Lindblad-Toh, K., Lander, E. S., Hinrichs, A., Trumbower, H., Clawson, H., Zweig, A., Kuhn, R. M., Barber, G., Harte, R., Karolchik, D., Field, M. A., Moore, R. A., Matthewson, C. A., Schein, J. E., Marra, M. A., Antonarakis, S. E., Batzoglou, S., Goldman, N., Hardison, R., Haussler, D., Miller, W., Pachter, L., Green, E. D. & Sidow, A. Analyses of deep mammalian sequence alignments and constraint predictions for 1% of the human genome. Genome Res 17, 760–774 (2007).

114. ENCODE Project Consortium, Birney, E., Stamatoyannopoulos, J. A., Dutta, A., Guigó, R., Gingeras, T. R., Margulies, E. H., Weng, Z., Snyder, M., Dermitzakis, E. T., Thurman, R. E., Kuehn, M. S., Taylor, C. M., Neph, S., Koch, C. M., Asthana, S., Malhotra, A., Adzhubei, I., Greenbaum, J. A., Andrews, R. M., Flicek, P., Boyle, P. J., Cao, H., Carter, N. P., Clelland, G. K., Davis, S., Day, N., Dhami, P., Dillon, S. C., Dorschner, M. O., Fiegler, H., Giresi, P. G., Goldy, J., Hawrylycz, M., Haydock, A., Humbert, R., James, K. D., Johnson, B. E., Johnson, E. M., Frum, T. T., Rosenzweig, E. R., Karnani, N., Lee, K., Lefebvre, G. C., Navas, P. A., Neri, F., Parker, S. C. J., Sabo, P. J., Sandstrom, R., Shafer, A., Vetrie, D., Weaver, M., Wilcox, S., Yu, M., Collins, F. S., Dekker, J., Lieb, J. D., Tullius, T. D., Crawford, G. E., Sunyaev, S., Noble, W. S., Dunham, I., Denoeud, F., Reymond, A., Kapranov, P., Rozowsky, J., Zheng, D., Castelo, R., Frankish, A., Harrow, J., Ghosh, S., Sandelin, A., Hofacker, I. L., Baertsch, R., Keefe, D., Dike, S., Cheng, J., Hirsch, H. A., Sekinger, E. A., Lagarde, J., Abril, J. F., Shahab, A., Flamm, C., Fried, C., Hackermüller, J., Hertel, J., Lindemeyer, M., Missal, K., Tanzer, A., Washietl, S., Korbel, J., Emanuelsson, O., Pedersen, J. S., Holroyd, N., Taylor, R., Swarbreck, D., Matthews, N., Dickson, M. C., Thomas, D. J., Weirauch, M. T., Gilbert, J., Drenkow, J., Bell, I., Zhao, X., Srinivasan, K. G., Sung, W.-K., Ooi, H. S., Chiu, K. P., Foissac, S., Alioto, T., Brent, M., Pachter, L., Tress, M. L., Valencia, A., Choo, S. W., Choo, C. Y., Ucla, C., Manzano, C., Wyss, C., Cheung, E., Clark, T. G., Brown, J. B., Ganesh, M., Patel, S., Tammana, H., Chrast, J., Henrichsen, C. N., Kai, C., Kawai, J., Nagalakshmi, U., Wu, J., Lian, Z., Lian, J., Newburger, P., Zhang, X., Bickel, P., Mattick, J. S., Carninci, P., Hayashizaki, Y., Weissman, S., Hubbard, T., Myers, R. M., Rogers, J., Stadler, P. F., Lowe, T. M., Wei, C.-L., Ruan, Y., Struhl, K., Gerstein, M., Antonarakis, S. E., Fu, Y., Green, E. D., Karaöz, U., Siepel, A., Taylor, J., Liefer, L. A., Wetterstrand, K. A., Good, P. J., Feingold, E. A., Guyer, M. S., Cooper, G. M., Asimenos, G., Dewey, C. N., Hou, M., Nikolaev, S., Montoya-Burgos, J. I., Löytynoja, A., Whelan, S., Pardi, F., Massingham, T., Huang, H., Zhang, N. R., Holmes, I., Mullikin, J. C., Ureta-Vidal, A., Paten, B., Seringhaus, M., Church, D., Rosenbloom, K., Kent, W. J., Stone, E. A., NISC Comparative Sequencing Program, Baylor College of Medicine Human Genome Sequencing Center, Washington University Genome Sequencing Center, Broad Institute, Children’s Hospital Oakland Research Institute, Batzoglou, S., Goldman, N., Hardison, R. C., Haussler, D., Miller, W., Sidow, A., Trinklein, N. D., Zhang, Z. D., Barrera, L., Stuart, R., King, D. C., Ameur, A., Enroth, S., Bieda, M. C., Kim, J., Bhinge, A. A., Jiang, N., Liu, J., Yao, F., Vega, V. B., Lee, C. W. H., Ng, P., Shahab, A., Yang, A., Moqtaderi, Z., Zhu, Z., Xu, X., Squazzo, S., Oberley, M. J., Inman, D., Singer, M. A., Richmond, T. A., Munn, K. J., Rada-Iglesias, A., Wallerman, O., Komorowski, J., Fowler, J. C., Couttet, P., Bruce, A. W., Dovey, O. M., Ellis, P. D., Langford, C. F., Nix, D. A., Euskirchen, G., Hartman, S., Urban, A. E., Kraus, P., Van Calcar, S., Heintzman, N., Kim, T. H., Wang, K., Qu, C., Hon, G., Luna, R., Glass, C. K., Rosenfeld, M. G., Aldred, S. F., Cooper, S. J., Halees, A., Lin, J. M., Shulha, H. P., Zhang, X., Xu, M., Haidar, J. N. S., Yu, Y., Ruan, Y., Iyer, V. R., Green, R. D., Wadelius, C., Farnham, P. J., Ren, B., Harte, R. A., Hinrichs, A. S., Trumbower, H., Clawson, H., Hillman-Jackson, J., Zweig, A. S., Smith, K., Thakkapallayil, A., Barber, G., Kuhn, R. M., Karolchik, D., Armengol, L., Bird, C. P., de Bakker, P. I. W., Kern, A. D., Lopez-Bigas, N., Martin, J. D., Stranger, B. E., Woodroffe, A., Davydov, E., Dimas, A., Eyras, E., Hallgrímsdóttir, I. B., Huppert, J., Zody, M. C., Abecasis, G. R., Estivill, X., Bouffard, G. G., Guan, X., Hansen, N. F., Idol, J. R., Maduro, V. V. B., Maskeri, B., McDowell, J. C., Park, M., Thomas, P. J., Young, A. C., Blakesley, R. W., Muzny, D. M., Sodergren, E., Wheeler, D. A., Worley, K. C., Jiang, H., Weinstock, G. M., Gibbs, R. A., Graves, T., Fulton, R., Mardis, E. R., Wilson, R. K., Clamp, M., Cuff, J., Gnerre, S., Jaffe, D. B., Chang, J. L., Lindblad-Toh, K., Lander, E. S., Koriabine, M., Nefedov, M., Osoegawa, K., Yoshinaga, Y., Zhu, B. & de Jong, P. J. Identification and analysis of functional elements in 1% of the human genome by the ENCODE pilot project. Nature 447, 799–816 (2007).

115. Cooper, G. M. & Brown, C. D. Qualifying the relationship between sequence conservation and molecular function. Genome Res 18, 201–205 (2008).

116. Villar, D., Berthelot, C., Aldridge, S., Rayner, T. F., Lukk, M., Pignatelli, M., Park, T. J., Deaville, R., Erichsen, J. T., Jasinska, A. J., Turner, J. M. A., Bertelsen, M. F., Murchison, E. P., Flicek, P. & Odom, D. T. Enhancer evolution across 20 mammalian species. Cell 160, 554–566 (2015).

117. Frankel, N., Davis, G. K., Vargas, D., Wang, S., Payre, F. & Stern, D. L. Phenotypic robustness conferred by apparently redundant transcriptional enhancers. Nature 466, 490–493 (2010).

118. Dudnyk, K., Cai, D., Shi, C., Xu, J. & Zhou, J. Sequence basis of transcription initiation in the human genome. Science 384, eadj0116 (2024).

119. Kircher, M., Xiong, C., Martin, B., Schubach, M., Inoue, F., Bell, R. J. A., Costello, J. F., Shendure, J. & Ahituv, N. Saturation mutagenesis of twenty disease-associated regulatory elements at single base-pair resolution. Nat. Commun. 10, 3583 (2019).

120. Patwardhan, R. P., Lee, C., Litvin, O., Young, D. L., Pe’er, D. & Shendure, J. High-resolution analysis of DNA regulatory elements by synthetic saturation mutagenesis. Nat Biotechnol 27, 1173–1175 (2009).

121. Zamudio-Martinez, E., Herrera-Campos, A. B., Muñoz, A., Rodríguez-Vargas, J. M. & Oliver, F. J. Tankyrases as modulators of pro-tumoral functions: molecular insights and therapeutic opportunities. J Exp Clin Cancer Res 40, 144 (2021).

122. Uhlén, M., Fagerberg, L., Hallström, B. M., Lindskog, C., Oksvold, P., Mardinoglu, A., Sivertsson, Å., Kampf, C., Sjöstedt, E., Asplund, A., Olsson, I., Edlund, K., Lundberg, E., Navani, S., Szigyarto, C. A.-K., Odeberg, J., Djureinovic, D., Takanen, J. O., Hober, S., Alm, T., Edqvist, P.-H., Berling, H., Tegel, H., Mulder, J., Rockberg, J., Nilsson, P., Schwenk, J. M., Hamsten, M., von Feilitzen, K., Forsberg, M., Persson, L., Johansson, F., Zwahlen, M., von Heijne, G., Nielsen, J. & Pontén, F. Proteomics. Tissue-based map of the human proteome. Science 347, 1260419 (2015).

123. Lek, M., Exome Aggregation Consortium, Karczewski, K. J., Minikel, E. V., Samocha, K. E., Banks, E., Fennell, T., O’Donnell-Luria, A. H., Ware, J. S., Hill, A. J., Cummings, B. B., Tukiainen, T., Birnbaum, D. P., Kosmicki, J. A., Duncan, L. E., Estrada, K., Zhao, F., Zou, J., Pierce-Hoffman, E., Berghout, J., Cooper, D. N., Deflaux, N., DePristo, M., Do, R., Flannick, J., Fromer, M., Gauthier, L., Goldstein, J., Gupta, N., Howrigan, D., Kiezun, A., Kurki, M. I., Moonshine, A. L., Natarajan, P., Orozco, L., Peloso, G. M., Poplin, R., Rivas, M. A., Ruano-Rubio, V., Rose, S. A., Ruderfer, D. M., Shakir, K., Stenson, P. D., Stevens, C., Thomas, B. P., Tiao, G., Tusie-Luna, M. T., Weisburd, B., Won, H.-H., Yu, D., Altshuler, D. M., Ardissino, D., Boehnke, M., Danesh, J., Donnelly, S., Elosua, R., Florez, J. C., Gabriel, S. B., Getz, G., Glatt, S. J., Hultman, C. M., Kathiresan, S., Laakso, M., McCarroll, S., McCarthy, M. I., McGovern, D., McPherson, R., Neale, B. M., Palotie, A., Purcell, S. M., Saleheen, D., Scharf, J. M., Sklar, P., Sullivan, P. F., Tuomilehto, J., Tsuang, M. T., Watkins, H. C., Wilson, J. G., Daly, M. J. & MacArthur, D. G. Analysis of protein-coding genetic variation in 60,706 humans. Nature 536, 285–291 (2016).

124. Karczewski, K. J., Francioli, L. C., Tiao, G., Cummings, B. B., Alföldi, J., Wang, Q., Collins, R. L., Laricchia, K. M., Ganna, A., Birnbaum, D. P., Gauthier, L. D., Brand, H., Solomonson, M., Watts, N. A., Rhodes, D., Singer-Berk, M., England, E. M., Seaby, E. G., Kosmicki, J. A., Walters, R. K., Tashman, K., Farjoun, Y., Banks, E., Poterba, T., Wang, A., Seed, C., Whiffin, N., Chong, J. X., Samocha, K. E., Pierce-Hoffman, E., Zappala, Z., O’Donnell-Luria, A. H., Minikel, E. V., Weisburd, B., Lek, M., Ware, J. S., Vittal, C., Armean, I. M., Bergelson, L., Cibulskis, K., Connolly, K. M., Covarrubias, M., Donnelly, S., Ferriera, S., Gabriel, S., Gentry, J., Gupta, N., Jeandet, T., Kaplan, D., Llanwarne, C., Munshi, R., Novod, S., Petrillo, N., Roazen, D., Ruano-Rubio, V., Saltzman, A., Schleicher, M., Soto, J., Tibbetts, K., Tolonen, C., Wade, G., Talkowski, M. E., Genome Aggregation Database Consortium, Neale, B. M., Daly, M. J. & MacArthur, D. G. The mutational constraint spectrum quantified from variation in 141,456 humans. Nature 581, 434–443 (2020).

125. Zeng, T., Spence, J. P., Mostafavi, H. & Pritchard, J. K. Bayesian estimation of gene constraint from an evolutionary model with gene features. bioRxivorg 2023.05.19.541520 (2024). doi:10.1101/2023.05.19.541520

126. Cheng, J., Novati, G., Pan, J., Bycroft, C., Žemgulytė, A., Applebaum, T., Pritzel, A., Wong, L. H., Zielinski, M., Sargeant, T., Schneider, R. G., Senior, A. W., Jumper, J., Hassabis, D., Kohli, P. & Avsec, Ž. Accurate proteome-wide missense variant effect prediction with AlphaMissense. Science 381, eadg7492 (2023).

127. Esposito, D., Weile, J., Shendure, J., Starita, L. M., Papenfuss, A. T., Roth, F. P., Fowler, D. M. & Rubin, A. F. MaveDB: an open-source platform to distribute and interpret data from multiplexed assays of variant effect. Genome Biol 20, 223 (2019).

128. Gersing, S., Cagiada, M., Gebbia, M., Gjesing, A. P., Coté, A. G., Seesankar, G., Li, R., Tabet, D., Weile, J., Stein, A., Gloyn, A. L., Hansen, T., Roth, F. P., Lindorff-Larsen, K. & Hartmann-Petersen, R. A comprehensive map of human glucokinase variant activity. Genome Biol 24, 97 (2023).

129. Macdonald, C. B., Nedrud, D., Grimes, P. R., Trinidad, D., Fraser, J. S. & Coyote-Maestas, W. DIMPLE: deep insertion, deletion, and missense mutation libraries for exploring protein variation in evolution, disease, and biology. Genome Biol 24, 36 (2023).

130. Scott, A., Hernandez, F., Chamberlin, A., Smith, C., Karam, R. & Kitzman, J. O. Saturation-scale functional evidence supports clinical variant interpretation in Lynch syndrome. Genome Biol 23, 266 (2022).

131. Fowler, D. M., Araya, C. L., Fleishman, S. J., Kellogg, E. H., Stephany, J. J., Baker, D. & Fields, S. High-resolution mapping of protein sequence-function relationships. Nat Methods 7, 741–746 (2010).

132. Bulik-Sullivan, B. K., Loh, P.-R., Finucane, H. K., Ripke, S., Yang, J., Patterson, N., Daly, M. J., Price, A. L. & Neale, B. M. LD Score regression distinguishes confounding from polygenicity in genome-wide association studies. Nature Genetics 47, 291–295 (2015).

133. Weissbrod, O., Hormozdiari, F., Benner, C., Cui, R., Ulirsch, J., Gazal, S., Schoech, A. P., van de Geijn, B., Reshef, Y., Márquez-Luna, C., O’Connor, L., Pirinen, M., Finucane, H. K. & Price, A. L. Functionally informed fine-mapping and polygenic localization of complex trait heritability. Nat Genet 52, 1355–1363 (2020).

134. Avsec, Ž., Weilert, M., Shrikumar, A., Krueger, S., Alexandari, A., Dalal, K., Fropf, R., McAnany, C., Gagneur, J., Kundaje, A. & Zeitlinger, J. Base-resolution models of transcription-factor binding reveal soft motif syntax. Nat Genet 53, 354–366 (2021).

135. Nguyen, E., Poli, M., Durrant, M. G., Kang, B., Katrekar, D., Li, D. B., Bartie, L. J., Thomas, A. W., King, S. H., Brixi, G., Sullivan, J., Ng, M. Y., Lewis, A., Lou, A., Ermon, S., Baccus, S. A., Hernandez-Boussard, T., Ré, C., Hsu, P. D. & Hie, B. L. Sequence modeling and design from molecular to genome scale with Evo. Science 386, eado9336 (2024).

136. Kelley, D. R., Snoek, J. & Rinn, J. L. Basset: learning the regulatory code of the accessible genome with deep convolutional neural networks. Genome Res 26, 990–999 (2016).

137. Linder, J., Srivastava, D., Yuan, H., Agarwal, V. & Kelley, D. R. Predicting RNA-seq coverage from DNA sequence as a unifying model of gene regulation. Nat Genet (2025). doi:10.1038/s41588-024-02053-6

138. Fudenberg, G., Kelley, D. R. & Pollard, K. S. Predicting 3D genome folding from DNA sequence with Akita. Nat Methods 17, 1111–1117 (2020).

139. Tabet, D. R., Kuang, D., Lancaster, M. C., Li, R., Liu, K., Weile, J., Coté, A. G., Wu, Y., Hegele, R. A., Roden, D. M. & Roth, F. P. Benchmarking computational variant effect predictors by their ability to infer human traits. Genome Biol 25, 172 (2024).

140. Lee, D. LS-GKM: a new gkm-SVM for large-scale datasets. Bioinformatics 32, 2196–2198 (2016).

141. Pedregosa, F., Varoquaux, G., Gramfort, A., Michel, V., Thirion, B., Grisel, O., Blondel, M., Louppe, G., Prettenhofer, P., Weiss, R., Weiss, R. J., Vanderplas, J., Passos, A., Cournapeau, D., Brucher, M., Perrot, M. & Duchesnay, E. Scikit-learn: Machine Learning in Python. J. Mach. Learn. Res. 12, 2825–2830 (2011).

142. Gupta, S., Stamatoyannopoulos, J. A., Bailey, T. L. & Noble, W. S. Quantifying similarity between motifs. Genome Biol. 8, R24 (2007).

143. Grant, C. E., Bailey, T. L. & Noble, W. S. FIMO: scanning for occurrences of a given motif. Bioinformatics 27, 1017–1018 (2011).

144. Mudge, J. M., Carbonell-Sala, S., Diekhans, M., Martinez, J. G., Hunt, T., Jungreis, I., Loveland, J. E., Arnan, C., Barnes, I., Bennett, R., Berry, A., Bignell, A., Cerdán-Vélez, D., Cochran, K., Cortés, L. T., Davidson, C., Donaldson, S., Dursun, C., Fatima, R., Hardy, M., Hebbar, P., Hollis, Z., James, B. T., Jiang, Y., Johnson, R., Kaur, G., Kay, M., Mangan, R. J., Maquedano, M., Gómez, L. M., Mathlouthi, N., Merritt, R., Ni, P., Palumbo, E., Perteghella, T., Pozo, F., Raj, S., Sisu, C., Steed, E., Sumathipala, D., Suner, M.-M., Uszczynska-Ratajczak, B., Wass, E., Yang, Y. T., Zhang, D., Finn, R. D., Gerstein, M., Guigó, R., Hubbard, T. J. P., Kellis, M., Kundaje, A., Paten, B., Tress, M. L., Birney, E., Martin, F. J. & Frankish, A. GENCODE 2025: reference gene annotation for human and mouse. Nucleic Acids Res 53, D966–D975 (2025).

145. Yeo, G. & Burge, C. B. Maximum entropy modeling of short sequence motifs with applications to RNA splicing signals. J Comput Biol 11, 377–394 (2004).

146. Meuleman, W., Muratov, A., Rynes, E., Halow, J., Lee, K., Bates, D., Diegel, M., Dunn, D., Neri, F., Teodosiadis, A., Reynolds, A., Haugen, E., Nelson, J., Johnson, A., Frerker, M., Buckley, M., Sandstrom, R., Vierstra, J., Kaul, R. & Stamatoyannopoulos, J. Index and biological spectrum of human DNase I hypersensitive sites. Nature 584, 244–251 (2020).

147. Seplyarskiy, V., Koch, E. M., Lee, D. J., Lichtman, J. S., Luan, H. H. & Sunyaev, S. R. A mutation rate model at the basepair resolution identifies the mutagenic effect of polymerase III transcription. Nat Genet 55, 2235–2242 (2023).

148. McLaren, W., Gil, L., Hunt, S. E., Riat, H. S., Ritchie, G. R. S., Thormann, A., Flicek, P. & Cunningham, F. The Ensembl Variant Effect Predictor. Genome Biol 17, 122 (2016).

149. Nystrom, S. L. & McKay, D. J. Memes: A motif analysis environment in R using tools from the MEME Suite. PLoS Comput Biol 17, e1008991 (2021).

150. Rauluseviciute, I., Riudavets-Puig, R., Blanc-Mathieu, R., Castro-Mondragon, J. A., Ferenc, K., Kumar, V., Lemma, R. B., Lucas, J., Chèneby, J., Baranasic, D., Khan, A., Fornes, O., Gundersen, S., Johansen, M., Hovig, E., Lenhard, B., Sandelin, A., Wasserman, W. W., Parcy, F. & Mathelier, A. JASPAR 2024: 20th anniversary of the open-access database of transcription factor binding profiles. Nucleic Acids Res 52, D174–D182 (2024).

151. Jin, H., Zhang, C., Zwahlen, M., von Feilitzen, K., Karlsson, M., Shi, M., Yuan, M., Song, X., Li, X., Yang, H., Turkez, H., Fagerberg, L., Uhlén, M. & Mardinoglu, A. Systematic transcriptional analysis of human cell lines for gene expression landscape and tumor representation. Nat Commun 14, 5417 (2023).

152. Kuleshov, M. V., Jones, M. R., Rouillard, A. D., Fernandez, N. F., Duan, Q., Wang, Z., Koplev, S., Jenkins, S. L., Jagodnik, K. M., Lachmann, A., McDermott, M. G., Monteiro, C. D., Gundersen, G. W. & Ma’ayan, A. Enrichr: a comprehensive gene set enrichment analysis web server 2016 update. Nucleic Acids Res 44, W90–7 (2016).

