## supplementary figures and data for "Identifying non-coding variant effects at scale via machine learning models of cis-regulatory reporter assays"


**Data availability** 2

Supplementary Dataset 1: MPAC_UKBB_BBJ_GTEx_MPRA_predictions 2

Supplementary Dataset 2: MPAC_DNase_ASE_predictions 2

Supplementary Dataset 3: MPAC_ClinVar_predictions 2

Supplementary Dataset 4: MPAC_COSMIC_predictions 2

Supplementary Dataset 5: MPAC_gnomAD_predictions 2

Supplementary Dataset 6: MPAC_GWAS_tag_SNVs+LD_predictions 2

Supplementary Dataset 7: MPAC_promoter_saturation_mutagenesis

Supplementary Dataset 8: Promoter_function_constraint_correlations 2

Supplementary Dataset 9: Promoter_gene_set_enrichment_results 2

Supplementary Dataset 10: MPAC_model_hyperparameters 2

Supplementary Dataset 11: MPAC_model_artifacts 2

**Code availability** 2

**Supplementary Figures** 3

Supplementary Fig. 1: MPAC accurately predicts MPRA activity on all autosomes 3

Supplementary Fig. 2: Description of MPAC prediction windowing 4

Supplementary Fig. 3: gkm-SVM summaries of missed emVar predictions 5

Supplementary Fig. 4: TF-MoDISco motifs of MPAC missed emVar predictions 6

Supplementary Fig. 5: MPAC emVar allelic skews correlate with DNase allele-specific effects (ASE) comparably to empirical MPRA emVars 7

Supplementary Fig. 6: All Sei DNase allele-specific effects correlations 8

Supplementary Fig. 7: All Enformer DNase allele-specific effects correlations 9

Supplementary Fig. 8: All Enformer MPRA emVar allelic-skew correlations 10

Supplementary Fig. 9: All Sei MPRA emVar allelic-skew correlations 11

Supplementary Fig. 10: Precision-recall curves of fine-mapped GWAS variants 12

Supplementary Fig. 11: Precision-recall curves of fine-mapped GTEx eQTLs 13

Supplementary Fig. 12: Identification of non-coding pathogenic variants by MPAC and empirical annotations 14

Supplementary Fig. 13: Summary of MPAC activity and allelic-skew predictions for gnomAD SNVs. 15

Supplementary Fig. 14: Summary of MPAC emVar predictions 16

Supplementary Fig. 15: Enrichment of gnomAD SNVs overlapping TF binding by allelic skew 17

Supplementary Fig. 16: Recent purifying selection and mutation rates of gnomAD SNVs by allelic skew 18

Supplementary Fig. 17: Overlap between emVars and constrained bases in gnomAD 19

Supplementary Fig. 18: Evolutionary constraint of gnomAD SNVs by MPAC allelic skew vs. Ensembl VEP 20

Supplementary Fig. 19: Evolutionary constraint of gnomAD SNVs by emVar pleiotropy or activity 21
Supplementary Fig. 20: Evolutionary constraint of gnomAD SNVs by allelic skew and allele frequency 22
Supplementary Fig. 21: MPAC activity and allelic-skew predictions and evolutionary constraint for all 18,658 promoter regions 23
Supplementary Fig. 22: Correlations between evolutionary constraint and MPAC activity and allelic-skew predictions stratified by distance to TSS 24
Supplementary Fig. 23: TF motif count profiles by distance to TSS 25
Supplementary Fig. 24: Correlation between promoter activity and gene expression 26
Supplementary Fig. 25: Additional promoters in bottom 10 by their correlations between constraint and negative allelic skew 27
Supplementary Fig. 26: Additional promoters in top 10 by their sum of both their correlations between constraint and positive or negative allelic skew 28
Supplementary Fig. 28: Gene-level constraint score distributions for individual promoters classified by their correlations between allelic skew and constraint 30

### Data availability

All MPAC predictions, additional datasets and model hyperparameters are available at Zenodo (https://doi.org/10.5281/zenodo.15178434) organized as follows:

**Supplementary Dataset 1: MPAC_UKBB_BBJ_GTEx_MPRA_predictions**
**Supplementary Dataset 2: MPAC_DNase_ASE_predictions**
**Supplementary Dataset 3: MPAC_ClinVar_predictions**
**Supplementary Dataset 4: MPAC_COSMIC_predictions**
**Supplementary Dataset 5: MPAC_gnomAD_predictions**
**Supplementary Dataset 6: MPAC_GWAS_tag_SNVs+LD_predictions**
**Supplementary Dataset 7: MPAC_promoter_saturation_mutagenesis**
**Supplementary Dataset 8: Promoter_function_constraint_correlations**
**Supplementary Dataset 9: Promoter_gene_set_enrichment_results**
**Supplementary Dataset 10: MPAC_model_hyperparameters**
**Supplementary Dataset 11: MPAC_model_artifacts**

### Code availability

All code and documentation are deposited online as follows. Code used for MPAC variant effect predictions and to analyze and visualize data related to the benchmarking, ClinVar, and COSMIC work is available at https://github.com/john-c-butts/MPAC/. Code used to analyze and visualize data related to the gnomAD and promoter saturation mutagenesis work is available at https://github.com/Reilly-Lab-Yale/MPAC_gnomAD_and_satmut.

### Supplementary Figures

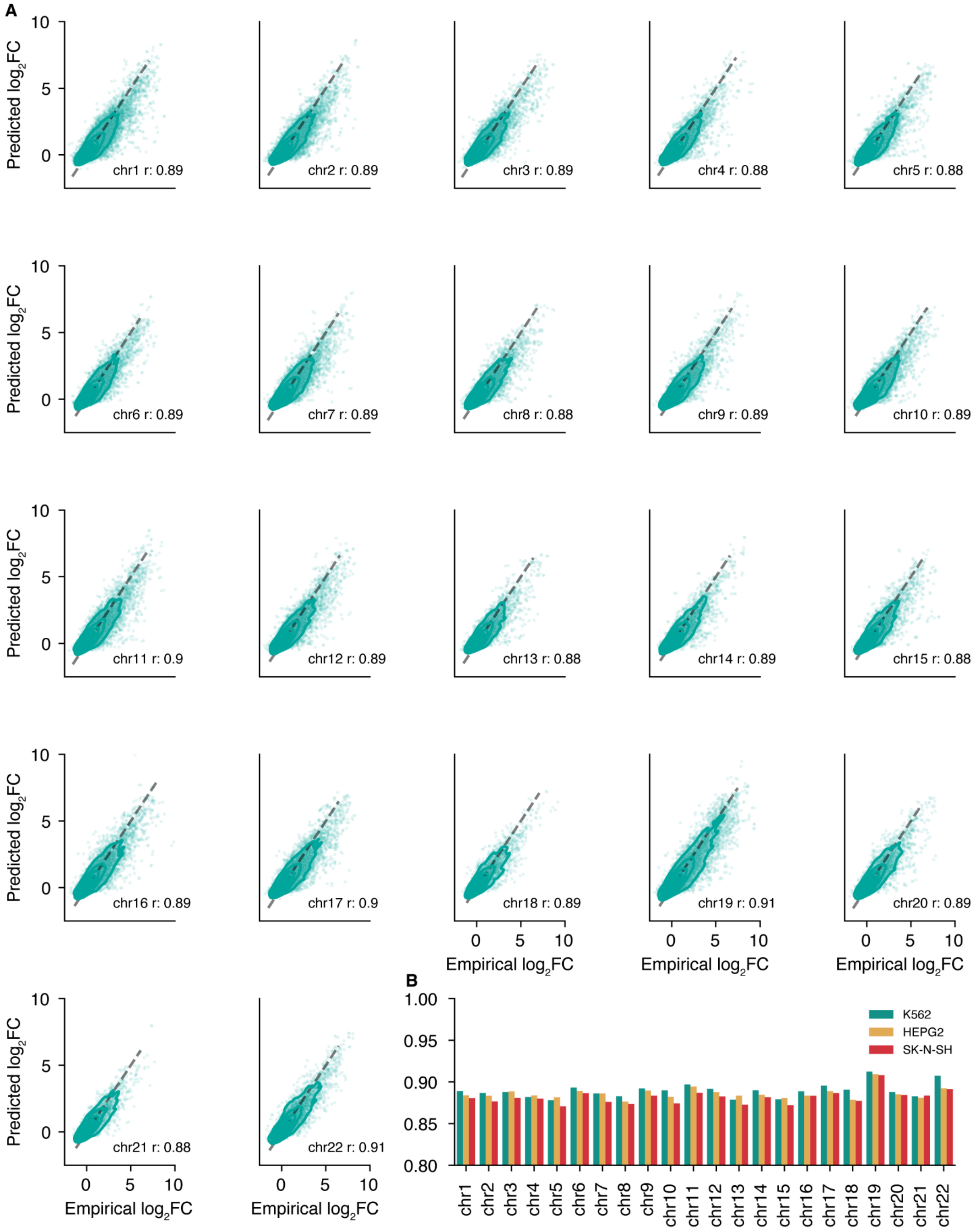


**Supplementary Fig. 1: MPAC accurately predicts MPRA activity on all autosomes**
**A)** Comparing MPAC activity predictions to MPRA shows high correlation across all autosomes.
**B)** MPAC performance is comparable across all three cell types, K562, HepG2, and SK-N-SH.

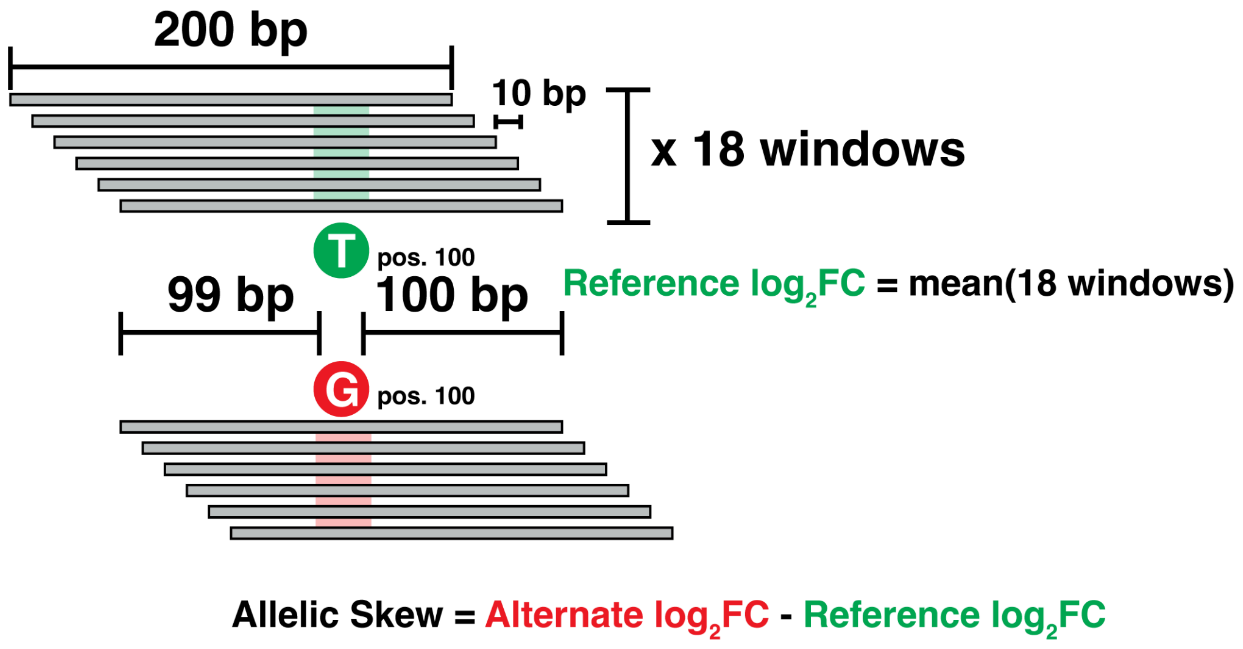


**Supplementary Fig. 2: Description of MPAC prediction windowing**
In order to capture a wider sequence context (370 bp total) the activity of each variant is tested in 18 windows where the variant's position within the 200 bp oligo is shifted by 10 bp. These 18 predictions are averaged for a final activity measurement ($log_2FC$) and allelic skew is calculated as the difference in $log_2FC$ activity between the alternate and reference alleles.

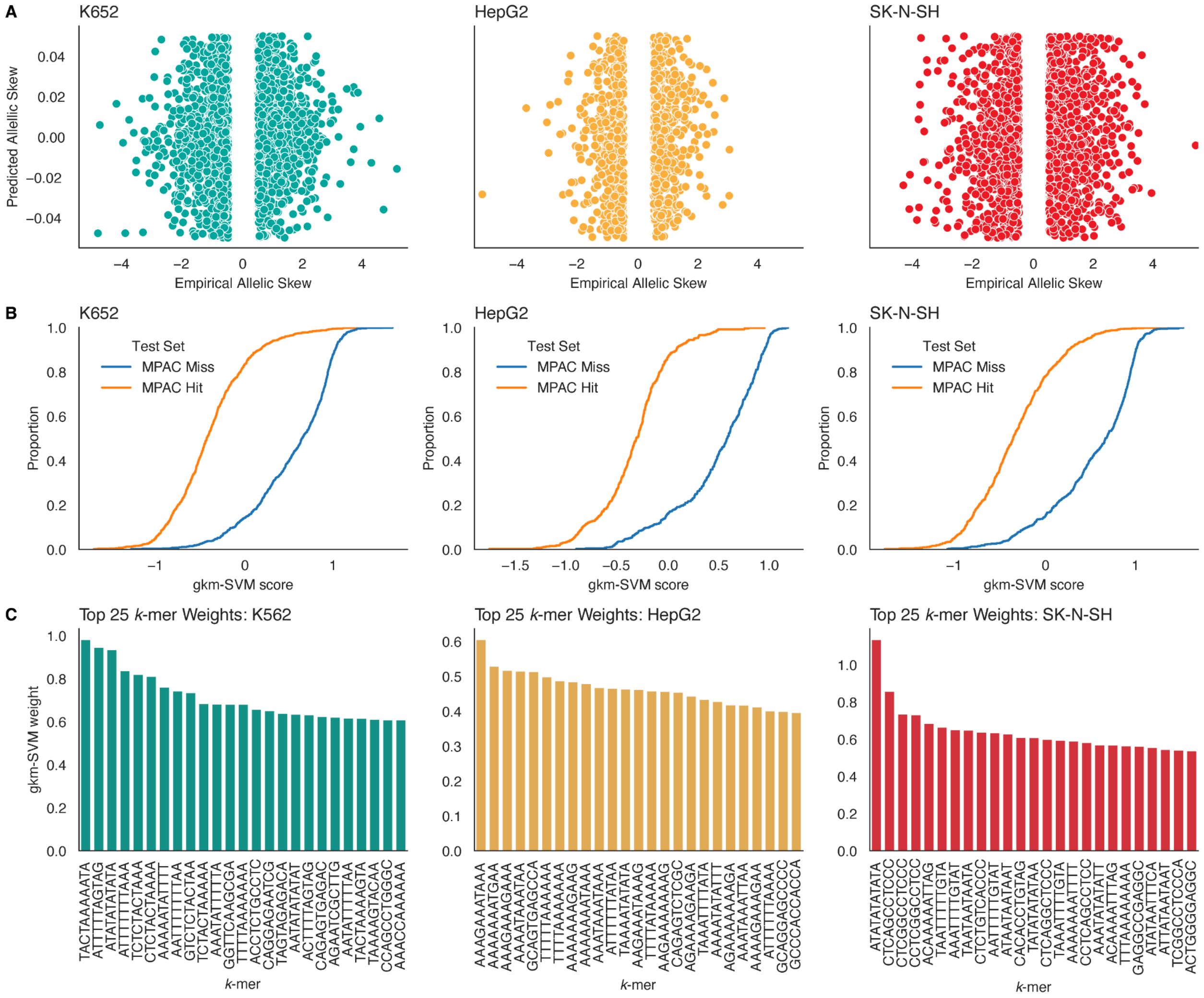


**Supplementary Fig. 3: gkm-SVM summaries of missed emVar predictions**

**A)** Scatterplots of MPAC emVar misses (|allelic skew| < 0.05) and hits (|allelic skew| > 0.5) used for LS-GKM training. **B)** Empirical cumulative density function plots of gkm-SVM model scores on test sets of MPAC hits and misses. **C)** Bar plots of weights for the top 25 *k*-mers for each LS-GKM model.

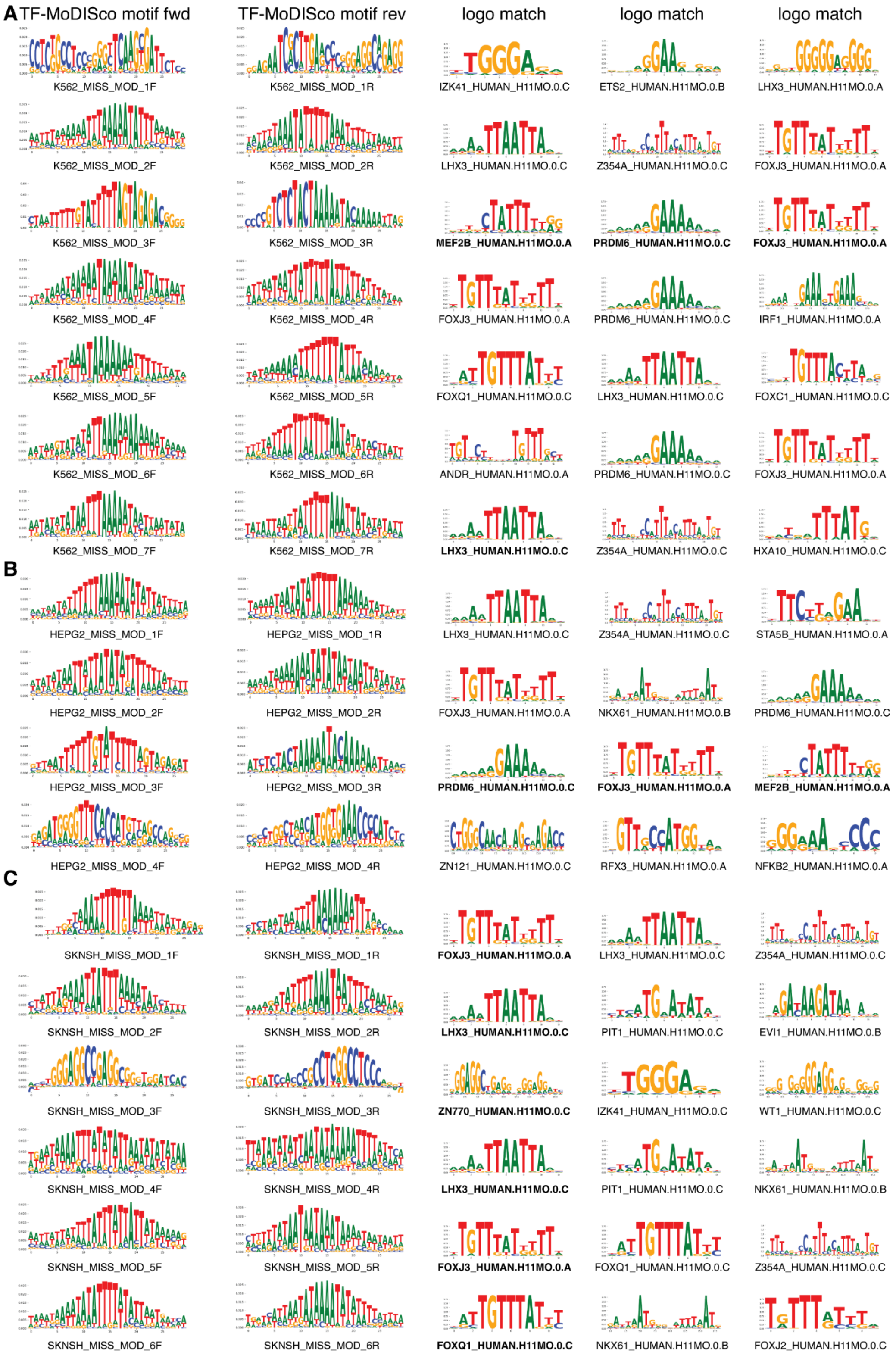


**Supplementary Fig. 4: TF-MoDISco motifs of MPAC missed emVar predictions**

Motifs for MPAC missed emVar predictions identified by TF-MoDISCO in **A)** K562, **B)** HepG2, **C)** SK-N-SH. Tomtom motif matches with FDR < 0.05 shown in bold.

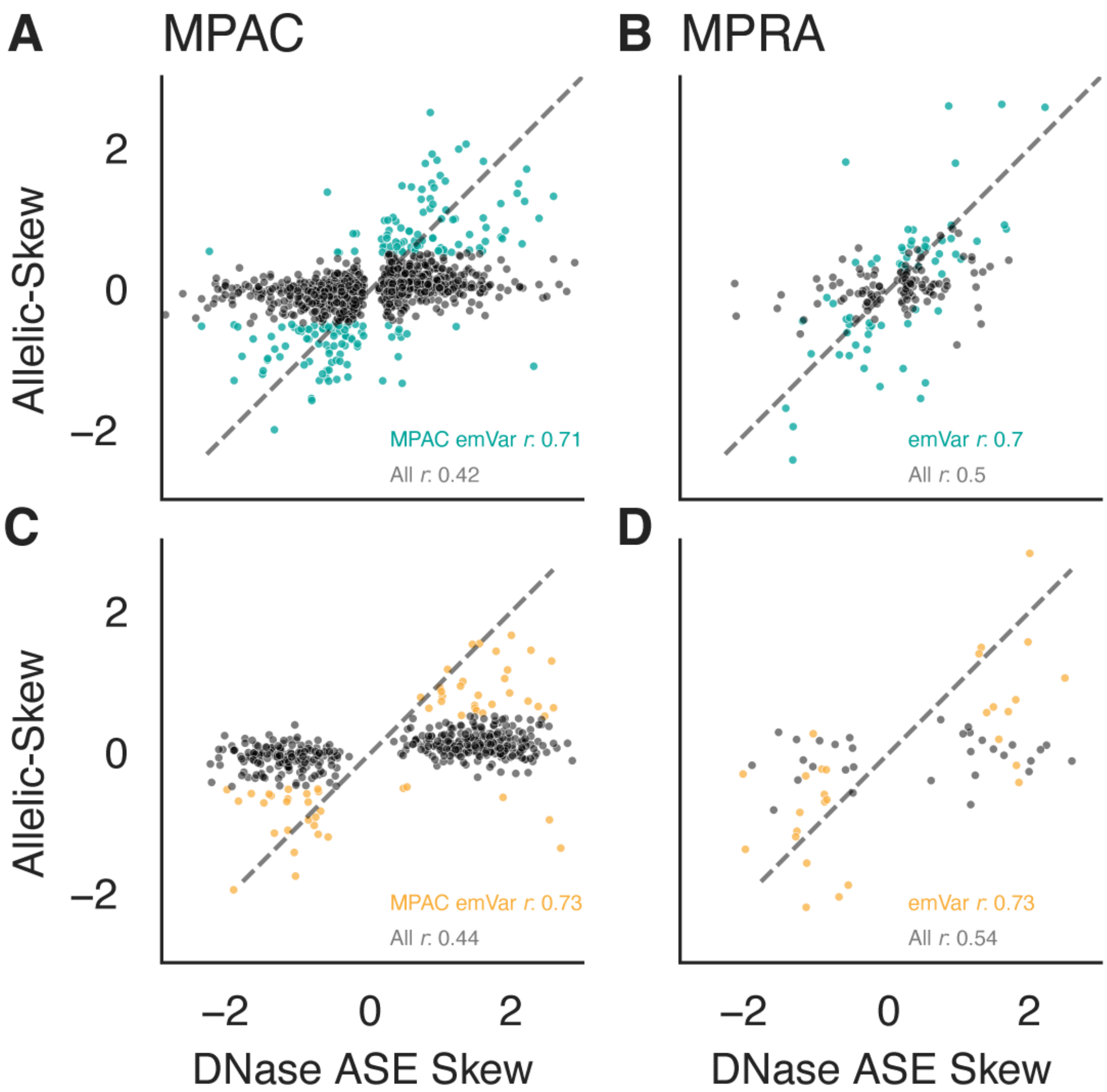


**Supplementary Fig. 5: MPAC emVar allelic skews correlate with DNase allele-specific effects (ASE) comparably to empirical MPRA emVars**
Variants with significant DNase ASE hits show moderate correlation to MPAC predictions in **A)** K562 (*n* = 1,270) while MPAC emVars (|allelic skew| > 0.5, colored blue) are highly correlated (*n* = 178). **B)** DNase ASE correlations with empirical MPRA are modestly higher than for MPAC predictions for all variants (*n* = 175) and comparable for empirically called emVars (*n* = 63). **C, D)** As in A,B for HepG2 (DNase ASE: *n* = 470, DNase ASE + MPAC emVar: *n* = 58, DNase ASE + MPRA: *n* = 59, DNase + MPRA emVar: *n* = 28).

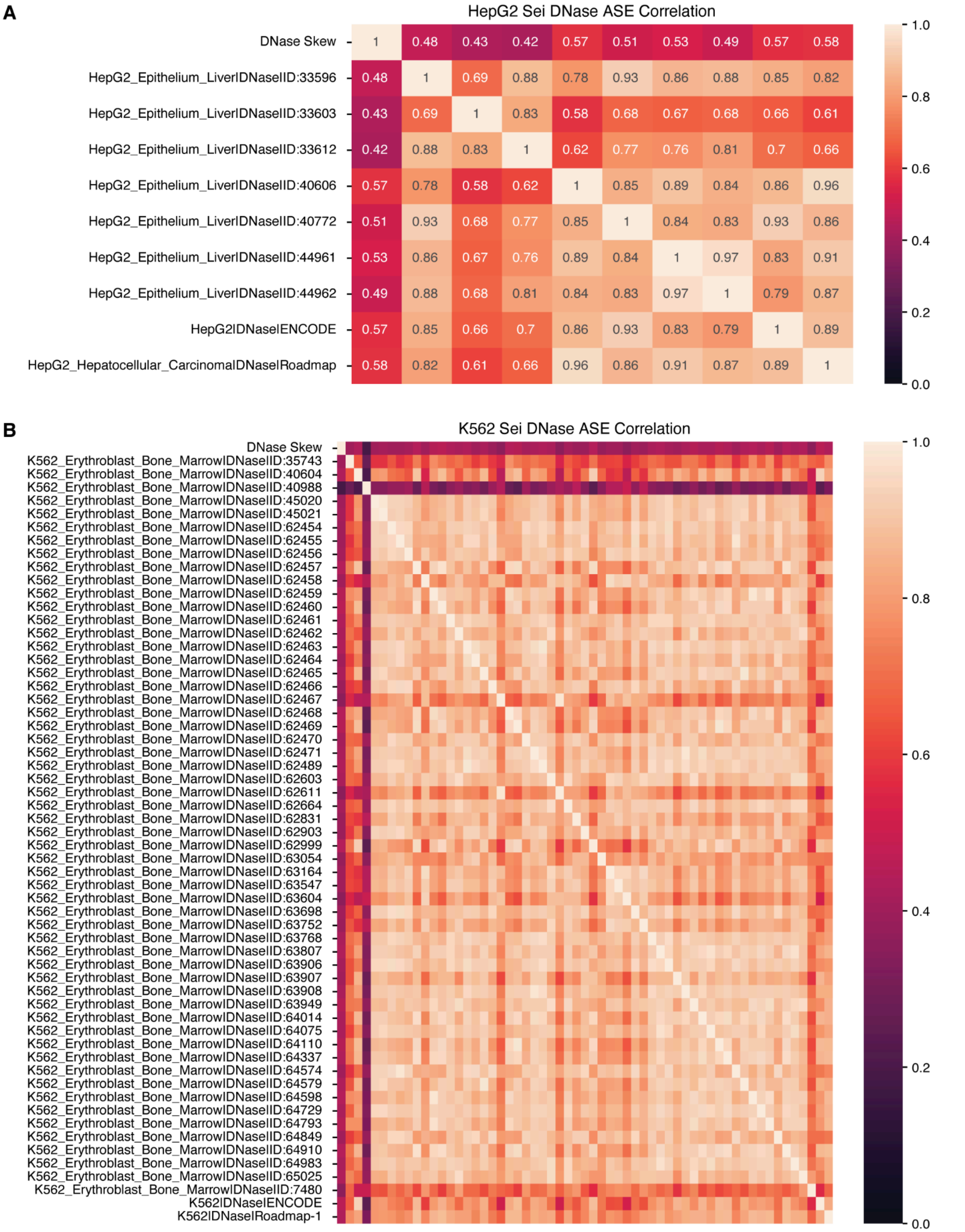


**Supplementary Fig. 6: All Sei DNase allele-specific effects correlations**

Pearson's correlations of Sei DNase predictions with empirical DNase ASE variants in **A)** HepG2 ($n$ = 331) and **B)** K562 ($n$ = 855).

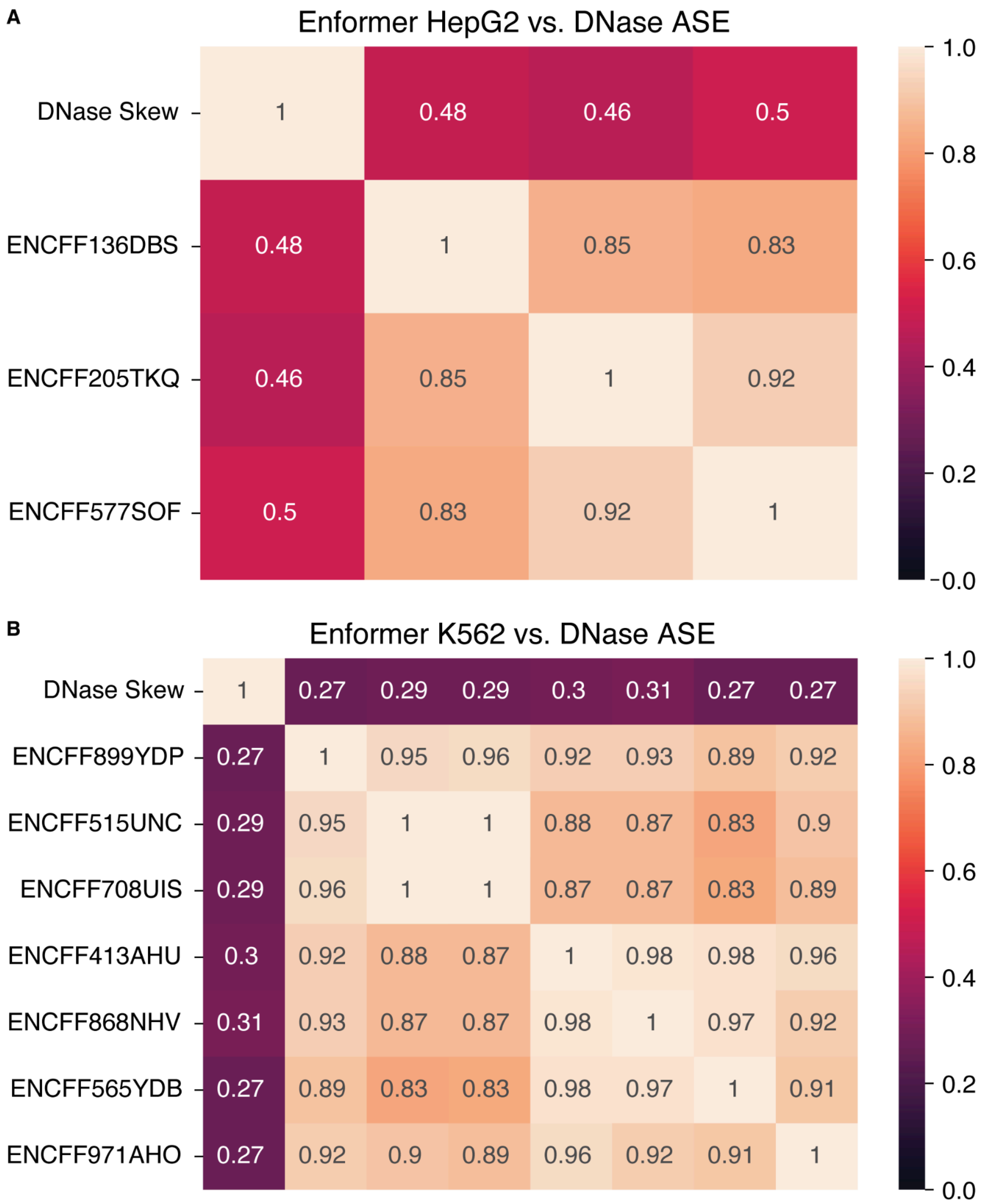


**Supplementary Fig. 7: All Enformer DNase allele-specific effects correlations**
Pearson’s correlations of Enformer DNase ASE predictions with empirical DNase ASE variants in **A)** HepG2 ($n$ = 331) and **B)** K562 ($n$ = 855).

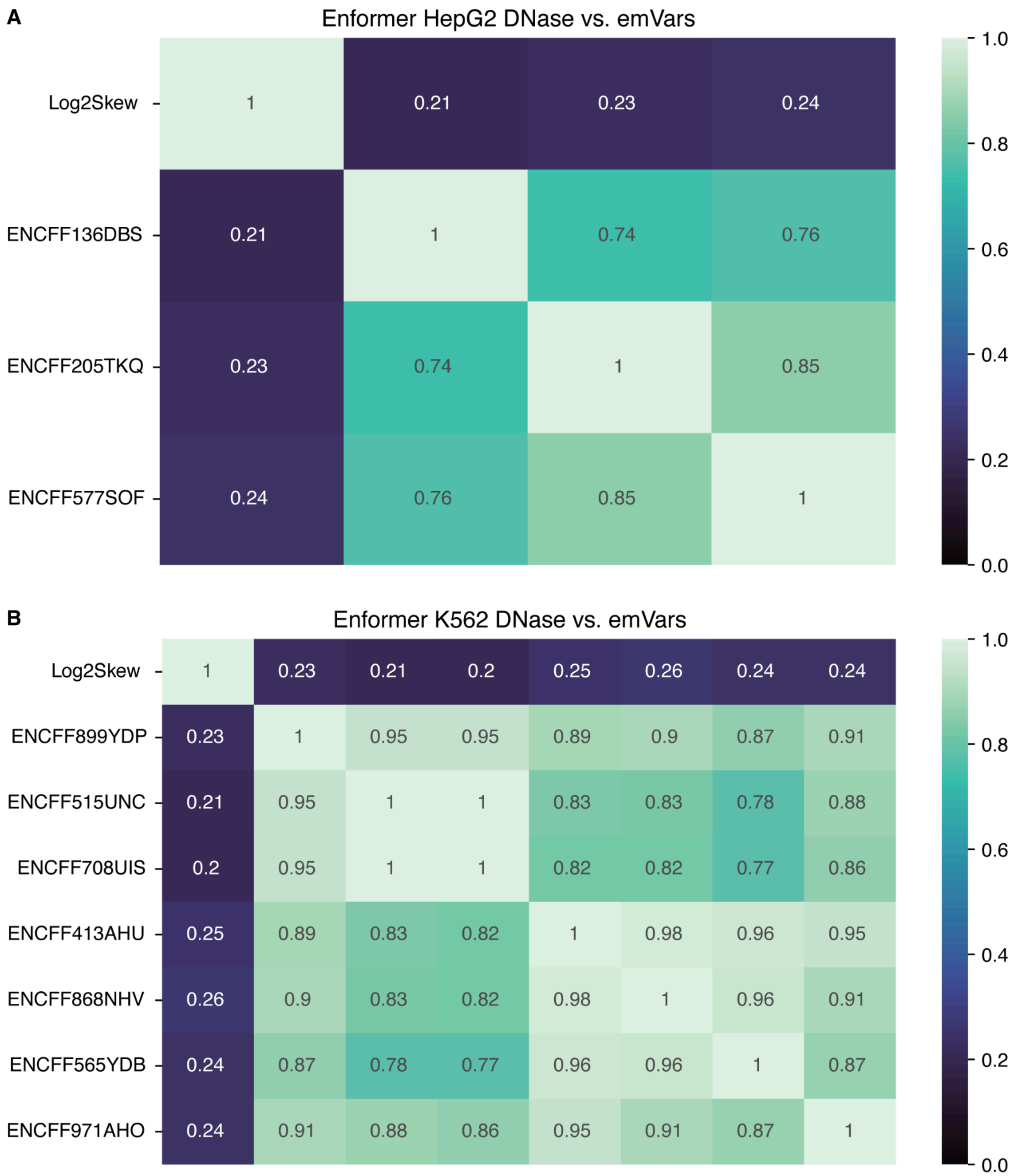


**Supplementary Fig. 8: All Enformer MPRA emVar allelic-skew correlations**
Pearson's correlations of Enformer DNase ASE predictions with empirical MPRA emVars in **A)** HepG2 (*n* = 10,407) and **B)** K562 (*n* = 11,993).

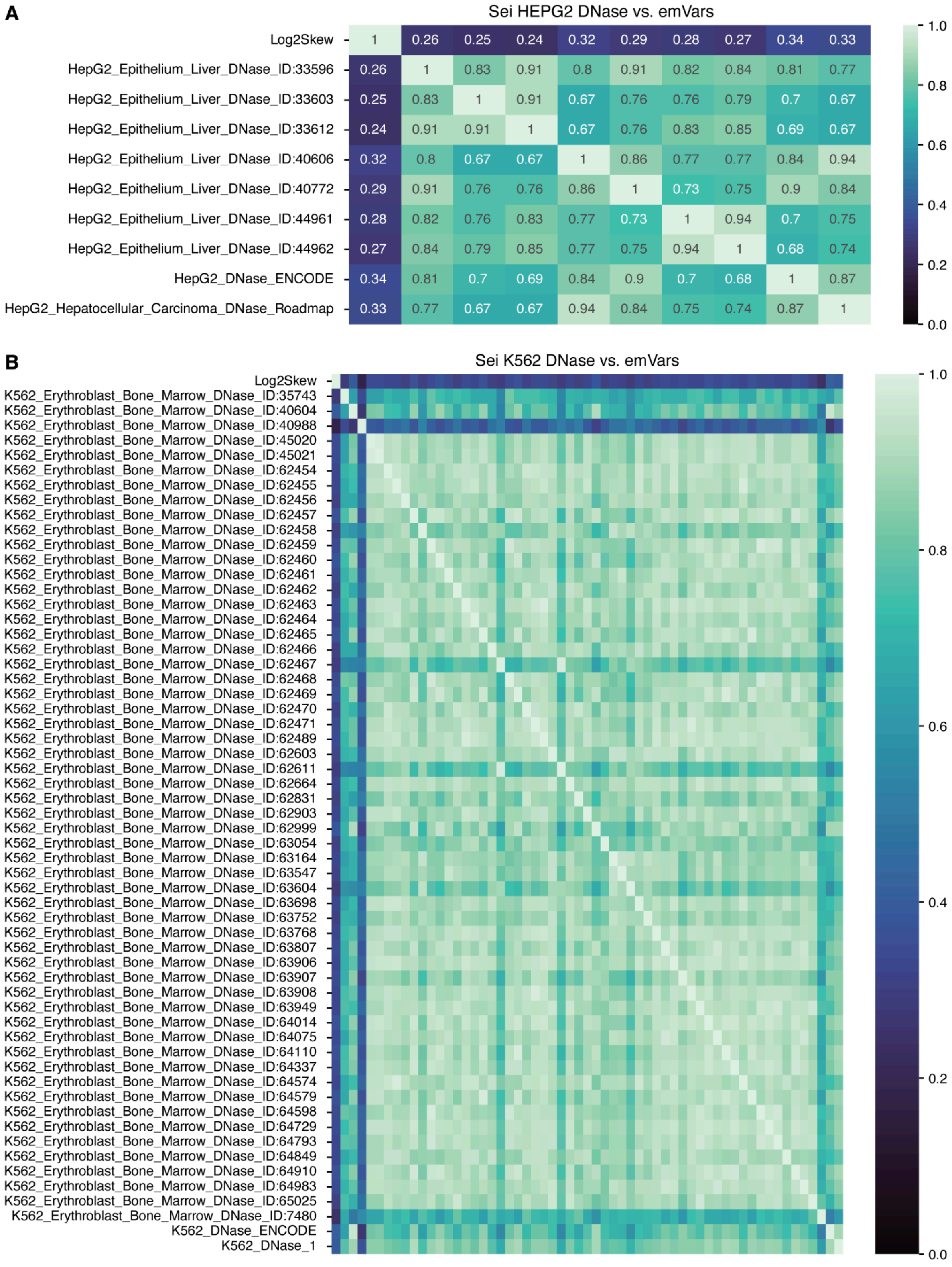


**Supplementary Fig. 9: All Sei MPRA emVar allelic-skew correlations**
Pearson’s correlations of Sei DNase ASE predictions with empirical MPRA emVars in **A)** HepG2 (*n* = 10,407) and **B)** K562 (*n* = 11,993).

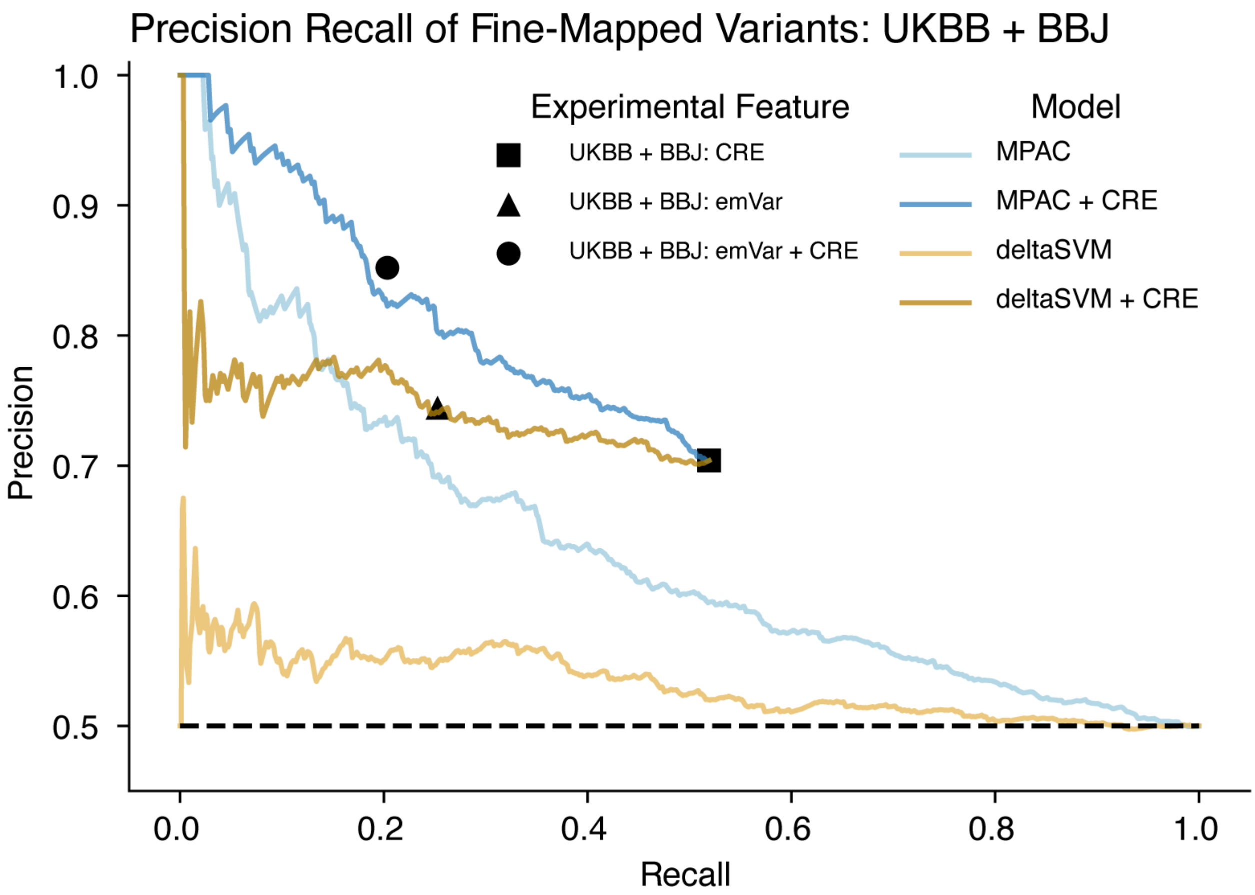


**Supplementary Fig. 10: Precision-recall curves of fine-mapped GWAS variants**
Precision-recall curves of MPAC, and deltaSVM predicted allelic skew of likely causal (high PIP > 0.9, *n* = 934) and non-causal (low PIP <0.01, *n* = 935) variants in UK BioBank (UKBB) or BioBank Japan (BBJ). See comparison between MPAC and Sei in **Fig. 1E**.

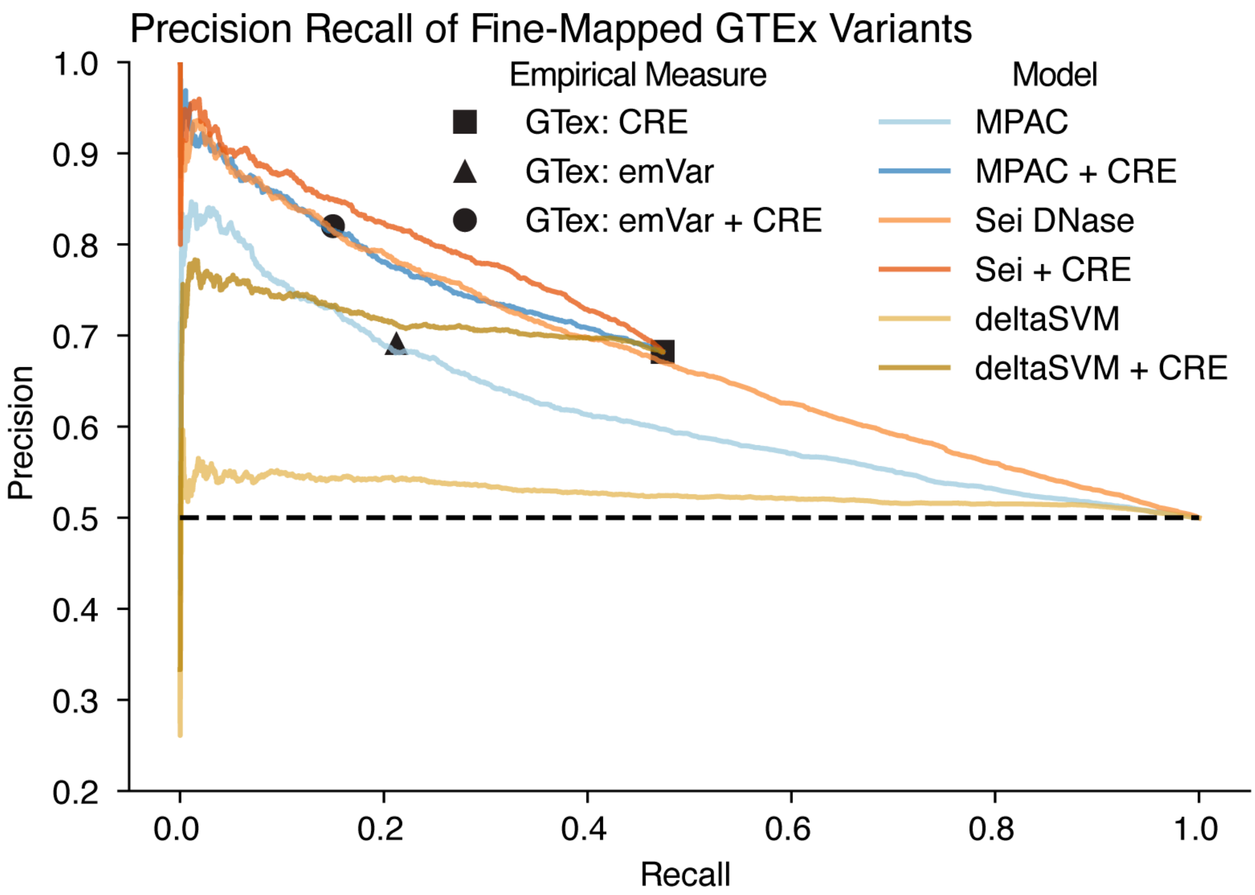


**Supplementary Fig. 11: Precision-recall curves of fine-mapped GTEx eQTLs**
Precision-recall curves of MPAC, Sei, and deltaSVM predicted allelic skew of likely causal (high PIP > 0.9, *n* = 11,354) and non-causal (low PIP <0.01, *n* = 11,366) eQTLs in GTEx**.**

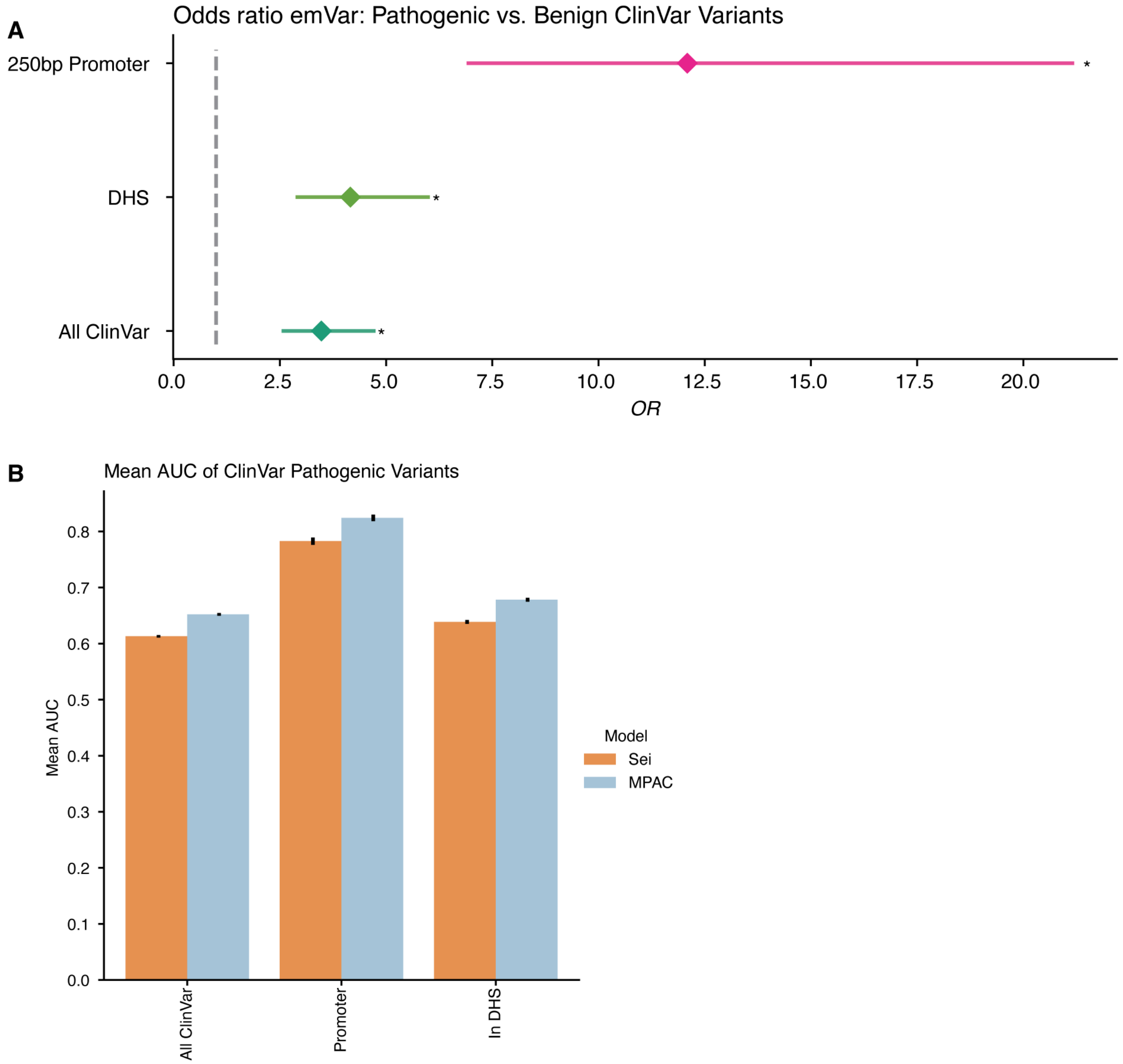


**Supplementary Fig. 12: Identification of non-coding pathogenic variants by MPAC and empirical annotations**

**A)** Odds ratio of pathogenic vs. benign variants being an emVar. All of ClinVar (*n* = 180,032) and subsets by regulatory features (250bp Promoter: *n* = 4,130, cCRE: *n* = 60,192 and DHS: *n* = 64,810). DHS: DNase I hypersensitive sites. **B)** Bar plot of mean AUPRC on classifying pathogenic vs. benign variants based on 100 randomly sampled benign variant sets equal in size to the number of pathogenic variants. Pathogenic/Benign variant counts in genomic regions: All ClinVar (573/573), 250bp Promoter (53/53), DHS (228/228). Error bars in B represent 95% CIs.

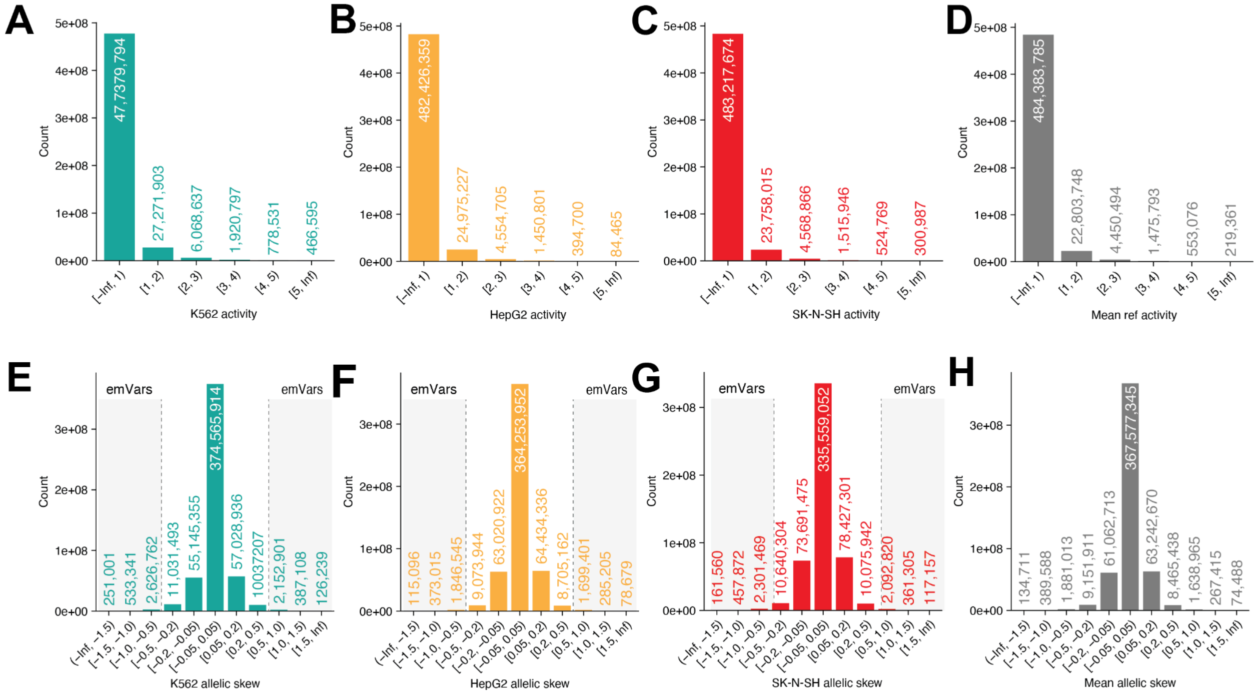


**Supplementary Fig. 13: Summary of MPAC activity and allelic-skew predictions for gnomAD SNVs.**

Counts of gnomAD SNVs in A-D) bins of activity ($log_2FC$) and E-H) bins of allelic skew ($log_2FC$) for K562, HepG2, SK-N-SH cell lines, and the means across three cell lines.

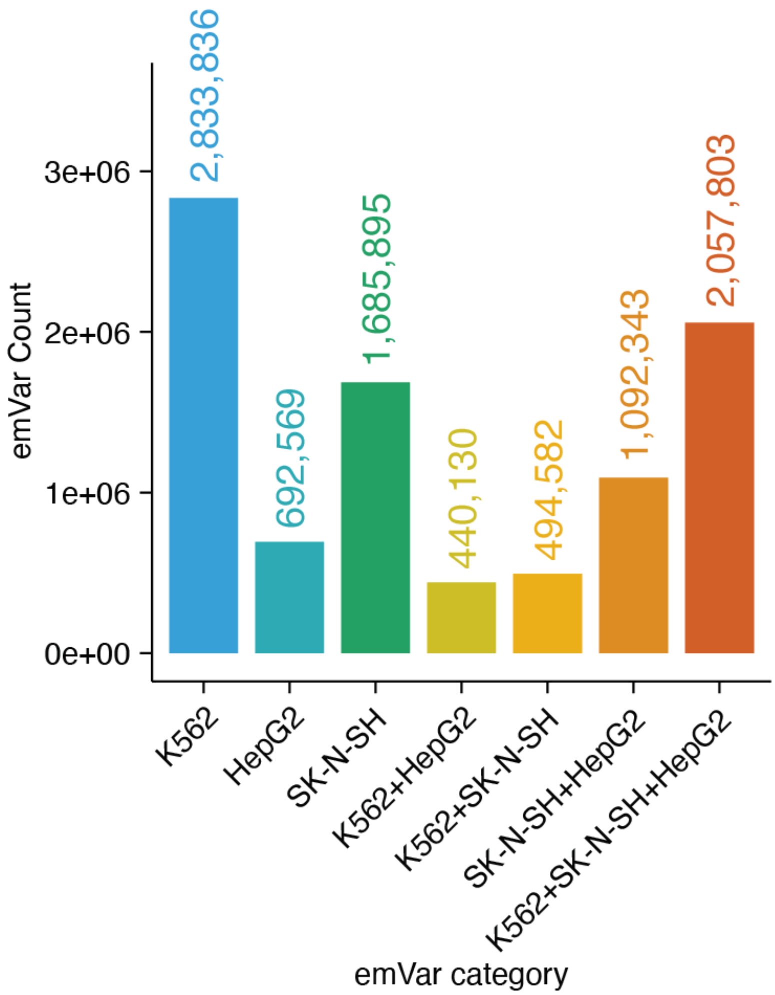


**Supplementary Fig. 14: Summary of MPAC emVar predictions**
Counts of gnomAD emVars by intersections across three cell lines.

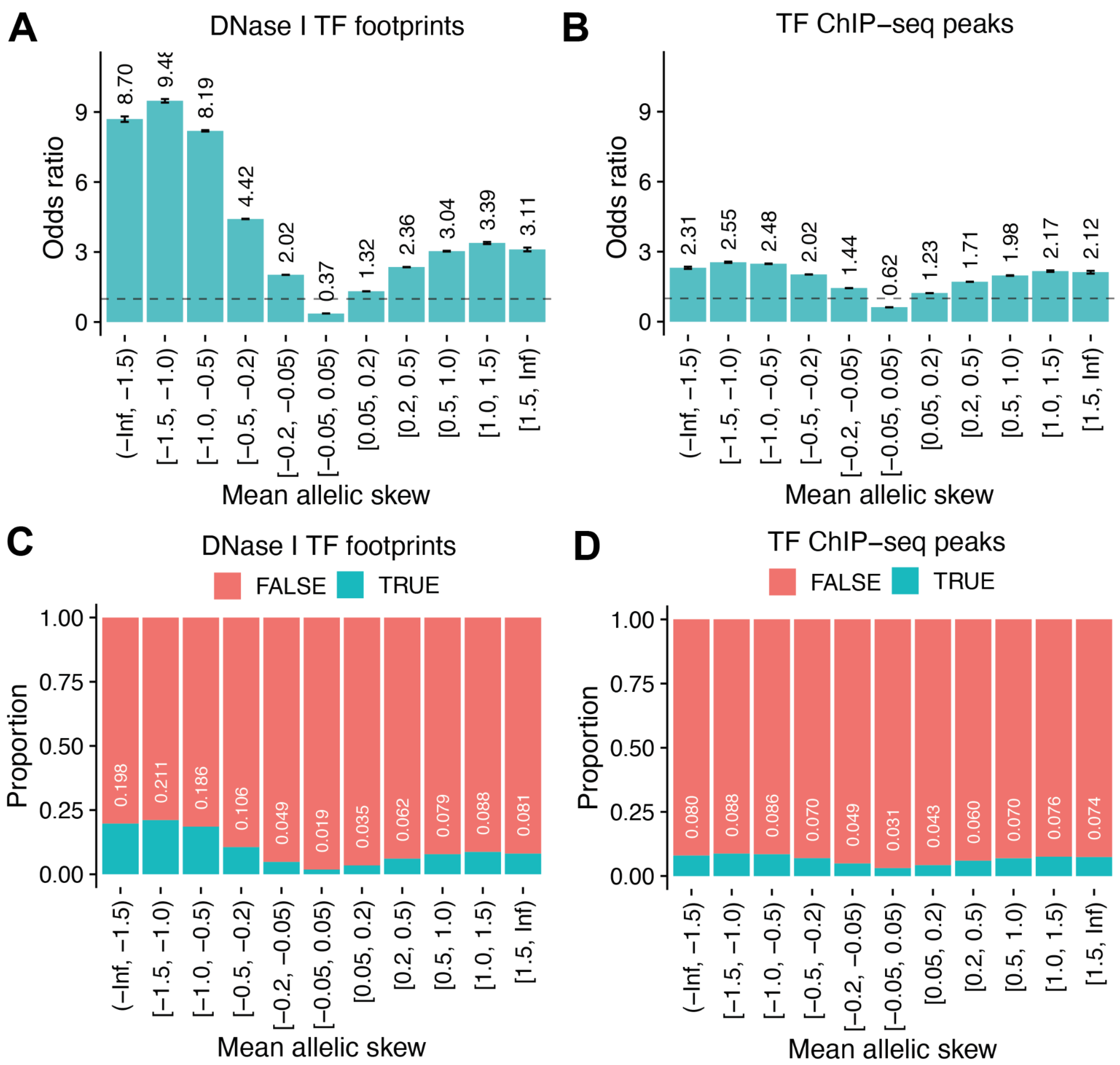


**Supplementary Fig. 15: Enrichment of gnomAD SNVs overlapping TF binding by allelic skew**

Odds ratio of SNVs overlapping vs not overlapping **A)** DNase I TF footprints or **B)** TF ChIP-seq peaks by allelic skew (mean $\log_2$FC across three cell lines). Proportion of total SNVs overlapping **C)** DNase I TF footprints or **D)** TF ChIP-seq peaks.

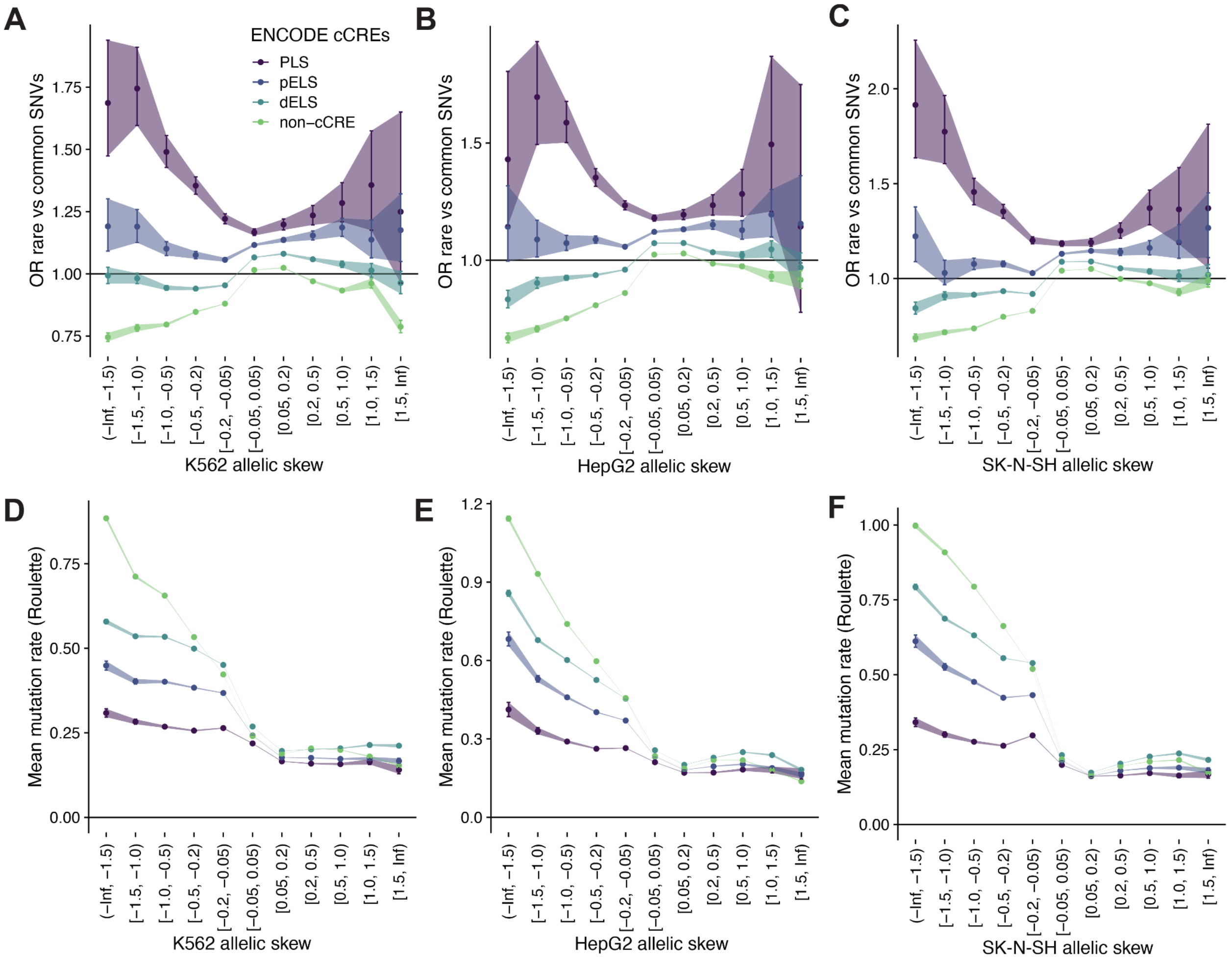


**Supplementary Fig. 16: Recent purifying selection and mutation rates of gnomAD SNVs by allelic skew**

Comparisons of within-species selective constraint as measured by the odds ratio (*OR*) of rare (allele frequency, AF < 0.1%) vs. common (AF ≥ 0.1%) SNVs in gnomAD, stratified by allelic skew ($log_2FC$) and across ENCODE cCREs classes (PLS: promoter-like signatures, dELS: proximal enhancer-like signatures, dELS: distal enhancer-like signatures, non-cCRE: regions outside of cCREs), for each of **A)** K562, **B)** HepG2, and **C)** SK-N-SH. Comparisons of *de novo* mutation rates of gnomAD SNVs as predicted by mean Roulette scores, stratified by MPAC allelic skew across ENCODE cCREs classes, for each of **D)** K562, **E)** HepG2, and **F)** SK-N-SH. Error bars represent 95% CIs.

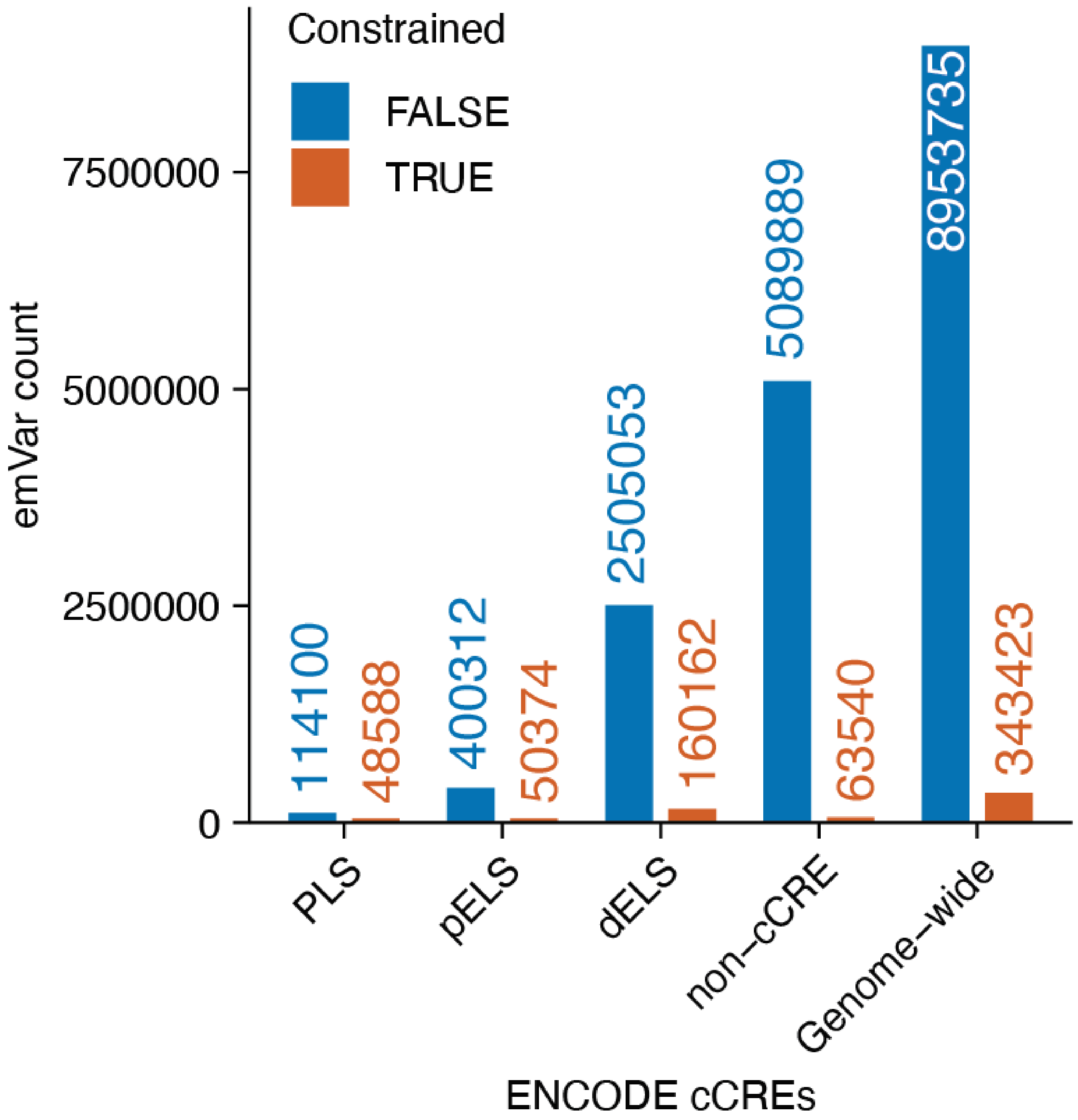


**Supplementary Fig. 17: Overlap between emVars and constrained bases in gnomAD**
Counts of gnomAD emVars overlapping evolutionarily constrained or unconstrained bases (defined as Zoonomia phyloP > 2.27 corresponding to FDR < 0.05) across ENCODE cCRE classes (PLS: promoter-like signatures, pELS: proximal enhancer-like signatures, dELS: distal enhancer-like signatures, non-cCRE: regions outside of cCREs).

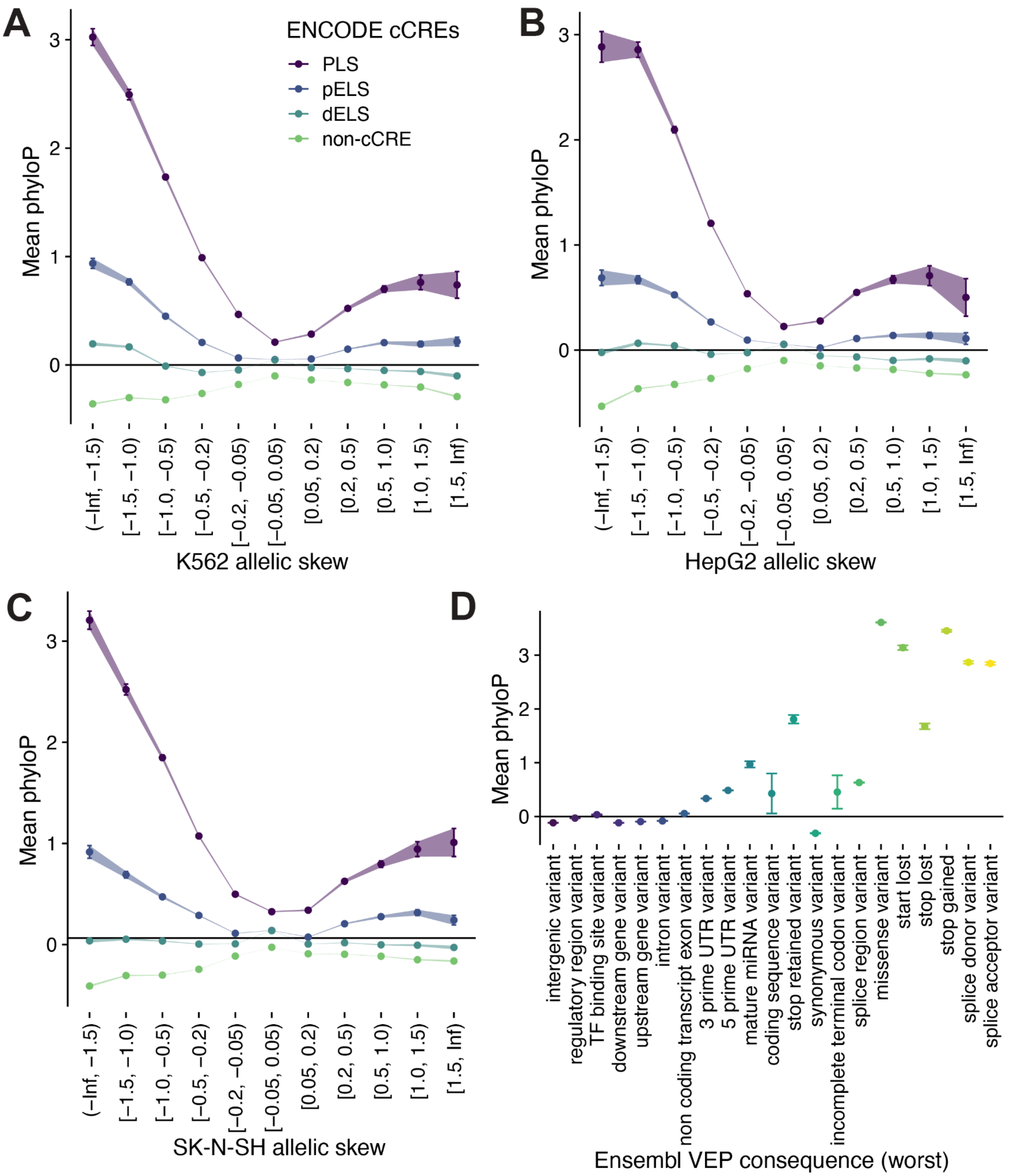


**Supplementary Fig. 18: Evolutionary constraint of gnomAD SNVs by MPAC allelic skew vs. Ensembl VEP**

Cross-species constraint (mean phyloP) of gnomAD SNVs by allelic skew ($log_2FC$) in individual cell lines across ENCODE cCREs classes (PLS: promoter-like signatures, pELS: proximal enhancer-like signatures, dELS: distal enhancer-like signatures, non-cCRE: regions outside of cCREs) for each of **A)** K562, **B)** HepG2, and **C)** SK-N-SH. **D)** Mean constraint of gnomAD SNVs grouped by Ensembl VEP consequence. Error bars represent 95% CIs.

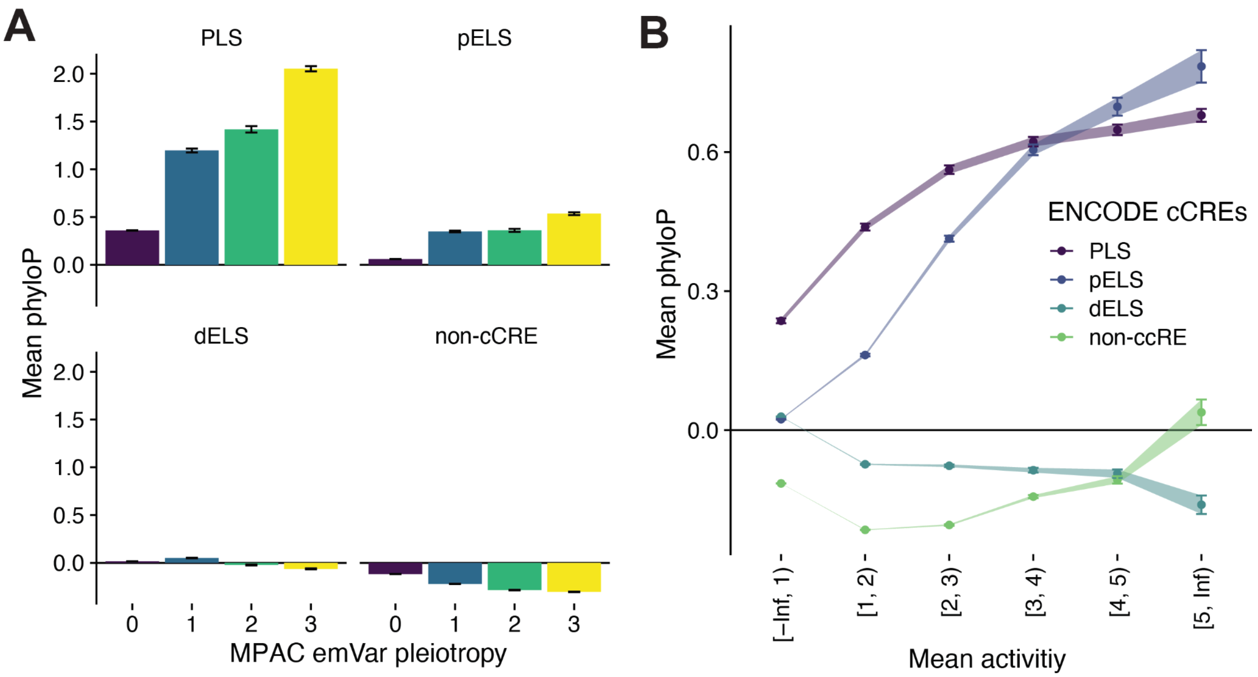


**Supplementary Fig. 19: Evolutionary constraint of gnomAD SNVs by emVar pleiotropy or activity**

Cross-species constraint (mean phyloP) of gnomAD SNVs **A)** by emVar pleiotropy (the number of cell lines where SNV is an emVar) across ENCODE cCREs classes (PLS: promoter-like signatures, pELS: proximal enhancer-like signatures, dELS: distal enhancer-like signatures, non-cCRE: regions outside of cCREs) or **B)** by activity (mean $\log_2$FC across three cell lines). Error bars represent 95% CIs.

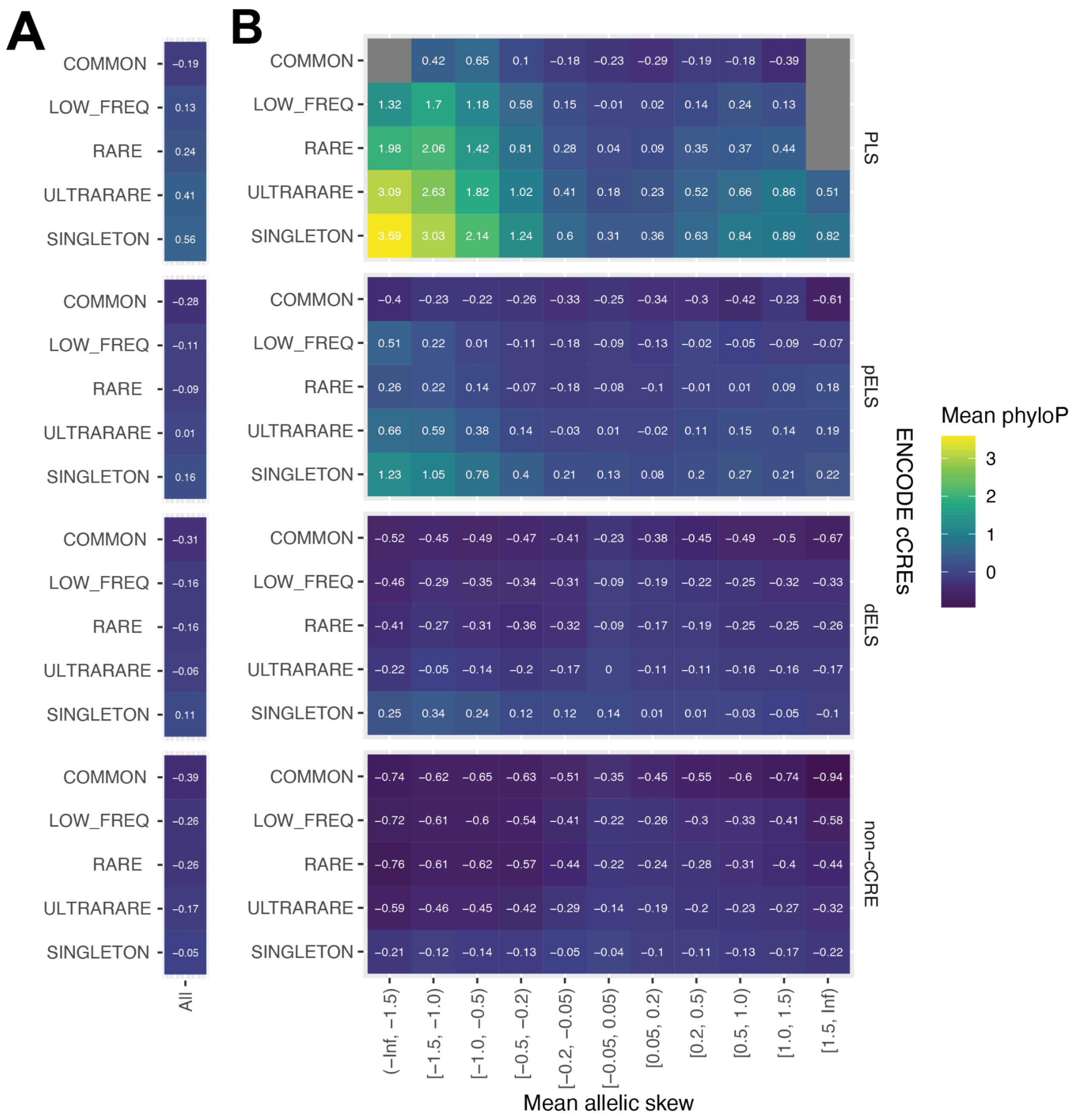


**Supplementary Fig. 20: Evolutionary constraint of gnomAD SNVs by allelic skew and allele frequency**

Cross-species constraint (mean phyloP) of gnomAD SNVs **A)** by allele frequency (AF) bins in gnomAD (SINGLETON: AF = 1, ULTRARARE: AF < 0.01%, RARE: AF < 0.1%, LOW_FREQ: AF < 1%, COMMON: AF >= 1%) and **B)** by both AF and allelic skew (mean $\log_2$FC across three cell lines) across ENCODE cCREs classes (PLS: promoter-like signatures, pELS: proximal enhancer-like signatures, dELS: distal enhancer-like signatures, non-cCRE: regions outside of cCREs).

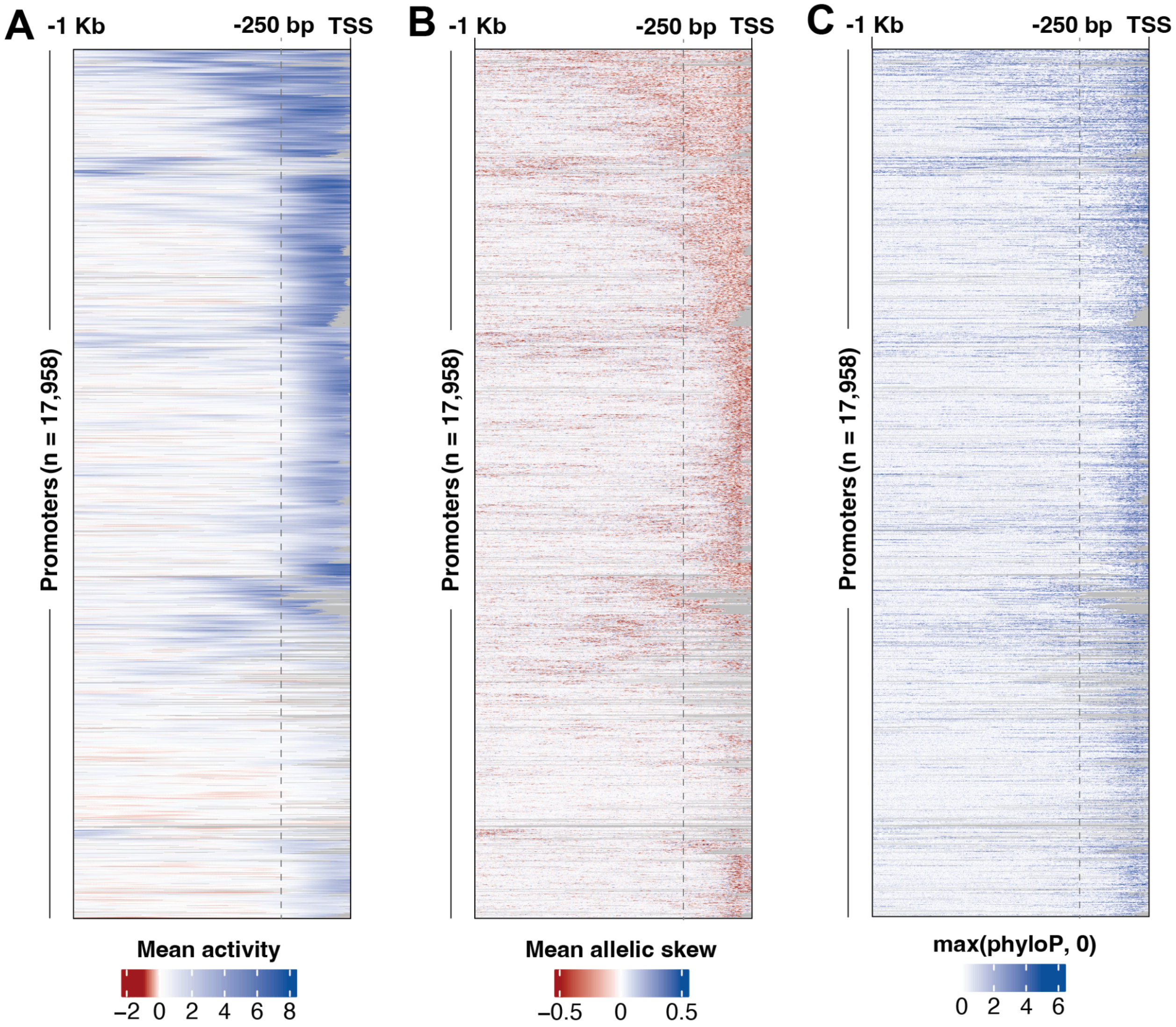


**Supplementary Fig. 21: MPAC activity and allelic-skew predictions and evolutionary constraint for all 18,658 promoter regions**

MPAC-predicted **A)** activity ($log_2FC$) and **B)** allelic-skew ($log_2FC$) predictions, and **C)** base-level evolutionary constraint (max(phyloP, 0)) at each position for all 18,658 promoters in the 1 kb upstream region of their annotated TSS. Activity and allelic skew at each position in a given promoter is the mean across all three possible SNVs and all three cell lines. The 250 bp upstream regions (right of grey dashed lines) correspond to those used in the **Fig. 4A-D** meta-promoter plots. Promoters are hierarchically clustered first by activity profiles and aligned across all three plots. Grey represents masked positions overlapping annotated exons and proximal intronic splice regions (-20 from 3' splice site or +6 from 5' splice site).

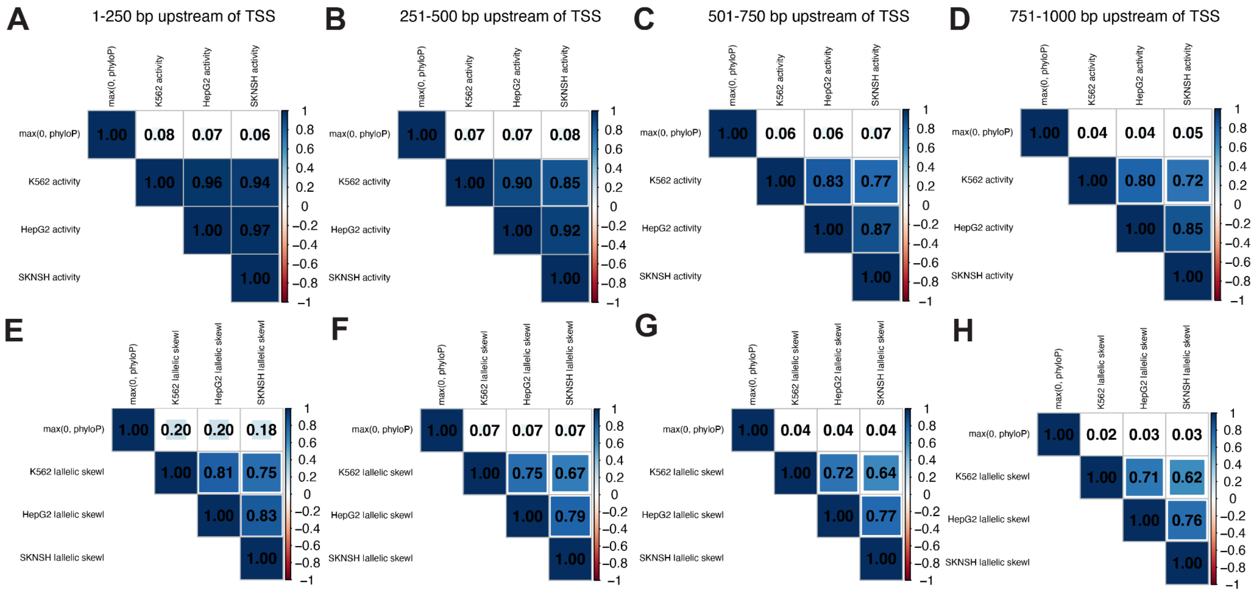


**Supplementary Fig. 22: Correlations between evolutionary constraint and MPAC activity and allelic-skew predictions stratified by distance to TSS**

Spearman's $\rho$ of constraint (max(phyloP, 0)) and **A-D)** activity or **E-H)** absolute allelic skew across three cell lines in distance bins of 1-250 bp, 251-500 bp, 501-750 bp, and 751-1000 bp upstream of the TSS.

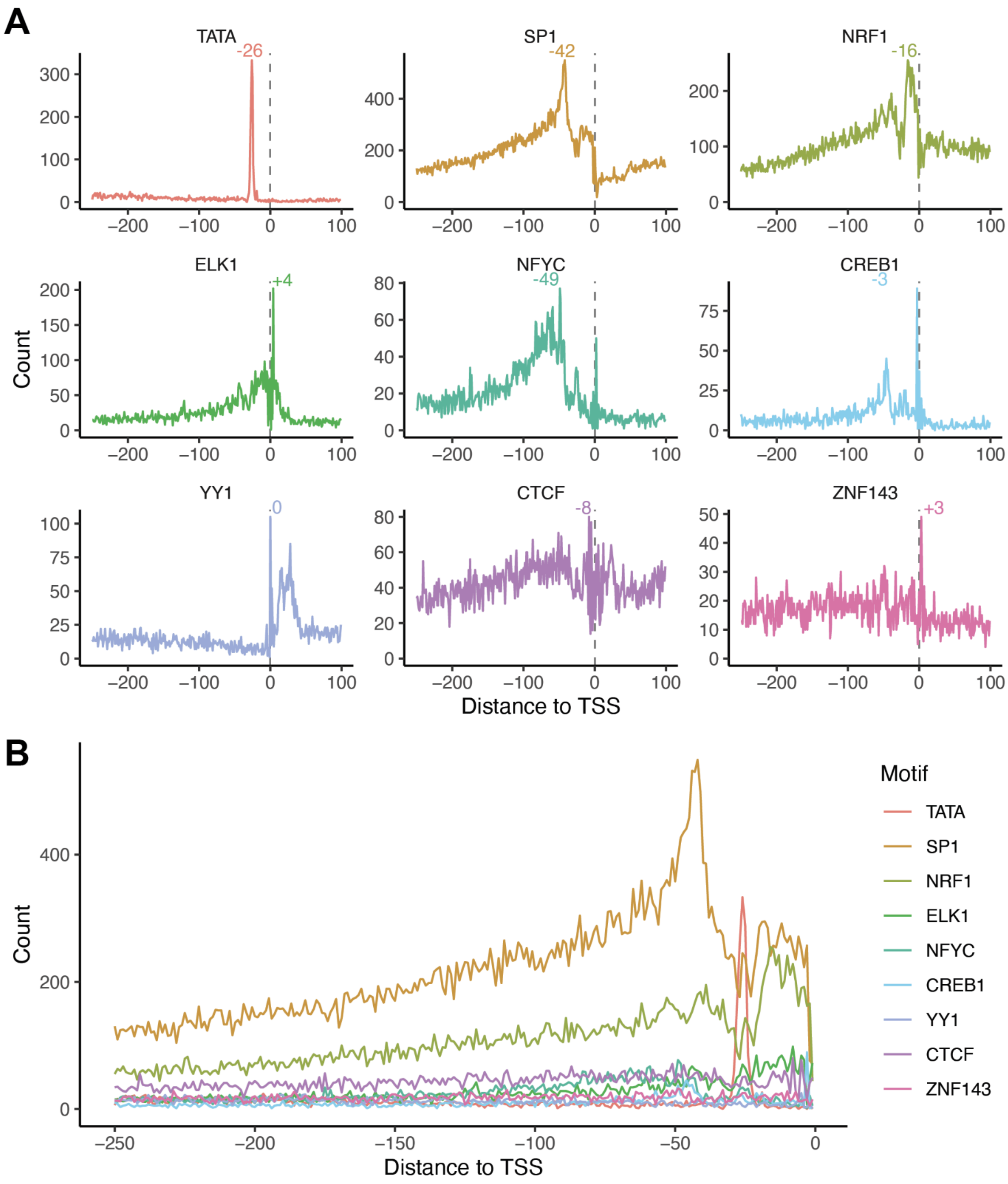


**Supplementary Fig. 23: TF motif count profiles by distance to TSS**

**A)** Motif count profiles by distance to TSS (dashed line) of promoter-binding TFs including TATA binding protein, GC binding SP1, NRF1, ETS family member ELK1, CCATT binding NFY subunit gamma NFYC, and ATF/CREB family member CREB1. We also included negative control transcription factors, YY1, which binds upstream of TSS, and CTCF and ZNF143, which are mainly involved in maintaining 3D enhancer-promoter looping. Positions of maximum motif counts are labeled. **B)** Comparison of motif count profiles across promoter-binding TFs. Motifs identified using FIMO with $p < 4 \times 10^{-4}$.

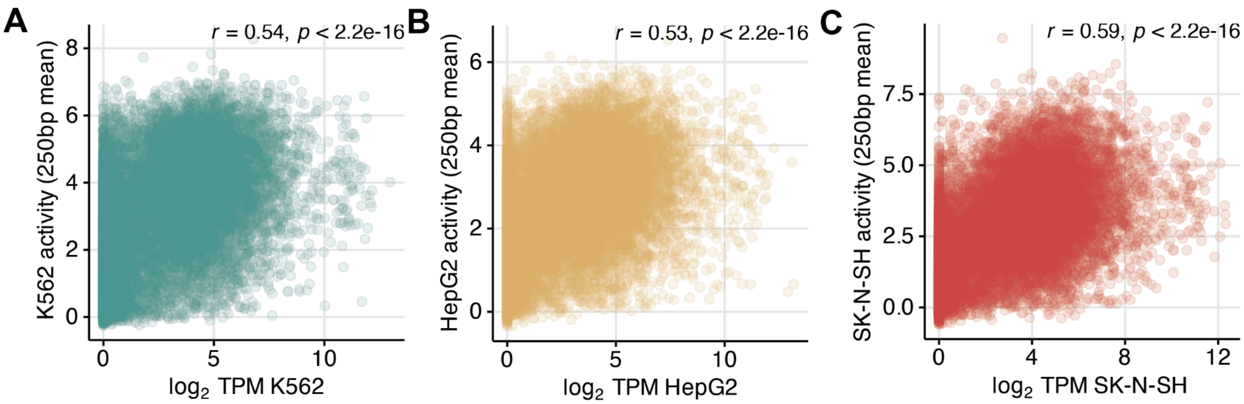


**Supplementary Fig. 24: Correlation between promoter activity and gene expression**
Pearson's *r* and *p*-values between the mean activity ($log_2FC$) in the upstream 250 bp of each promoter vs. gene expression ($log_{10}$ transcripts per million (TPM)) from the Human Protein Atlas for **A)** K562, **B)** HepG2, and **C)** SK-N-SH.

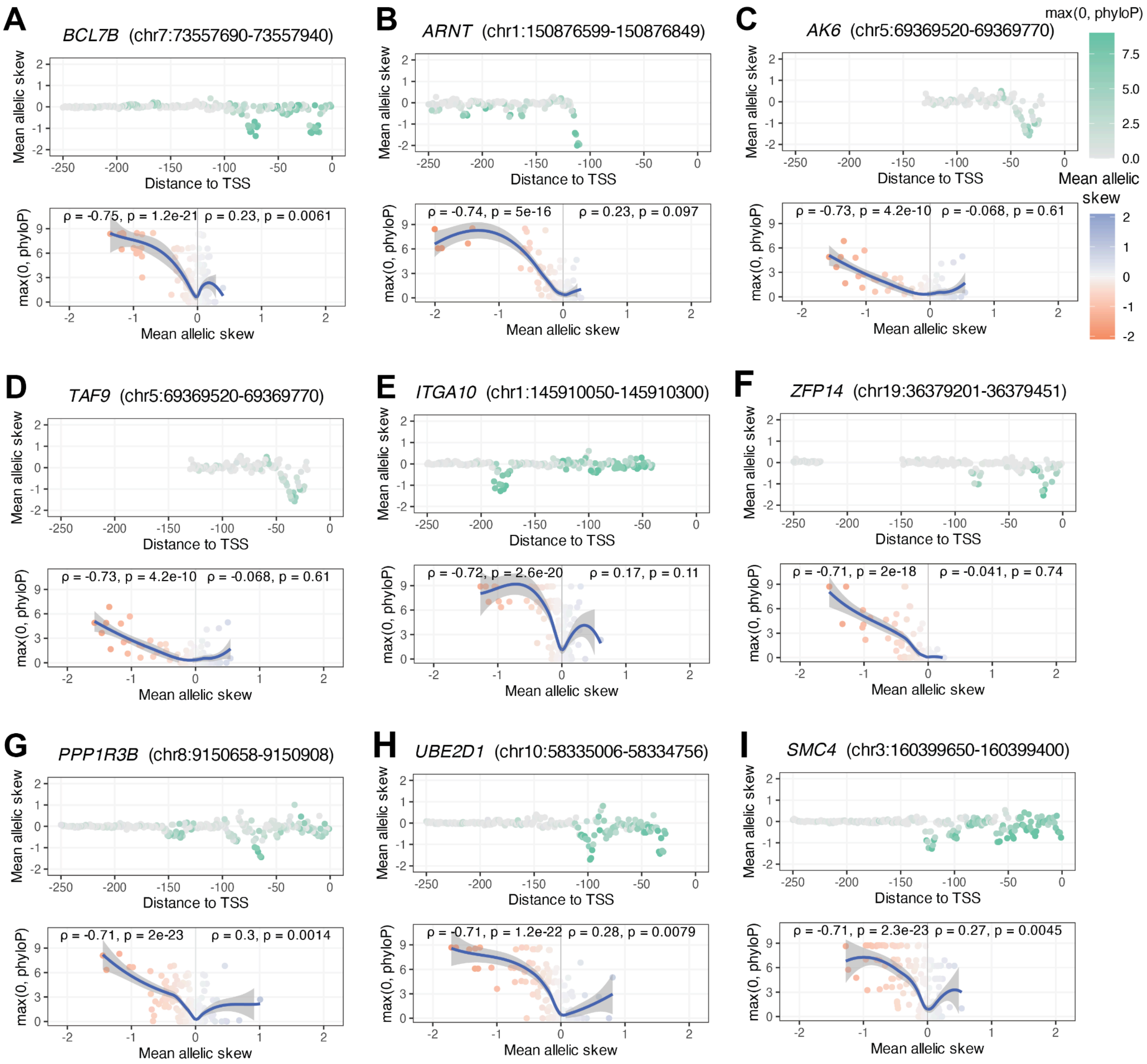


**Supplementary Fig. 25: Additional promoters in bottom 10 by their correlations between constraint and negative allelic skew**

Plots of allelic skew (mean $\log_2$FC of the three possible SNVs at each position) and constraint (max(0, phyloP)) by distance to TSS (top). Spearman correlation ($\rho$ and p-values) between constraint and allelic skew for mutations with negative allelic skew (bottom left) or mutations with positive allelic skew (bottom right). Promoters in the bottom 10 by negative allelic skew $\rho$ excluding *TNKS2* (see **Fig. 4J**). LOESS curve with 95% CI. Missing data overlap annotated exons and proximal intronic splice regions, including those of alternative transcripts and other genes.

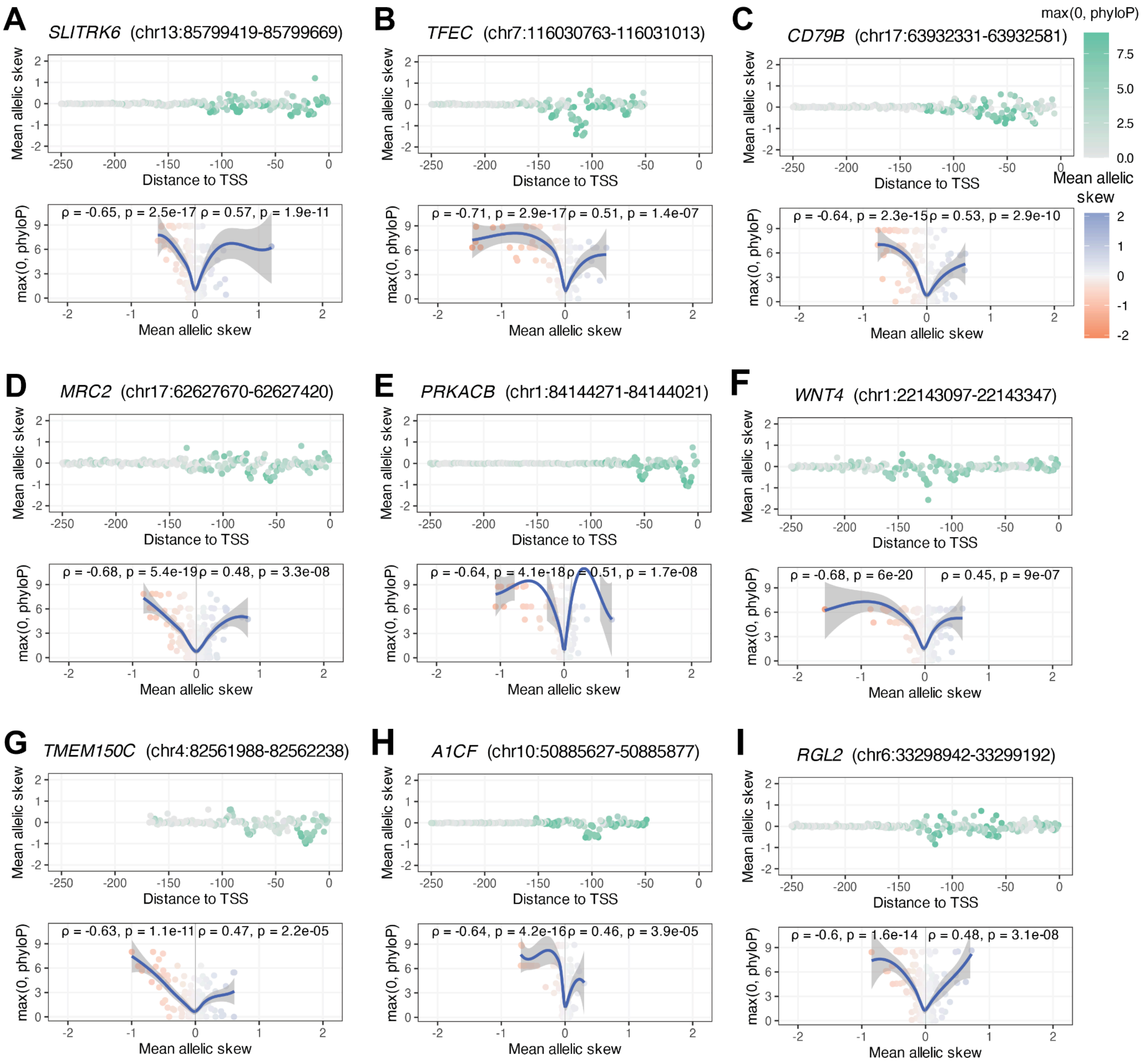


**Supplementary Fig. 26: Additional promoters in top 10 by their sum of both their correlations between constraint and positive or negative allelic skew**

Plots of allelic skew (mean $log_2$FC of the three possible SNVs at each position) and constraint (max(0, phyloP)) by distance to TSS (top). Spearman correlation ($\rho$ and p-values) between constraint and allelic skew for mutations with negative allelic skew (bottom left) or mutations with positive allelic skew (bottom right). Promoters in the top 10 by sum of absolute negative and positive allelic-skew $\rho$'s excluding *CCR10* (see **Fig. 4K**). LOESS curve with 95% CI. Missing data overlap annotated exons and proximal intronic splice regions, including those of alternative transcripts and other genes.

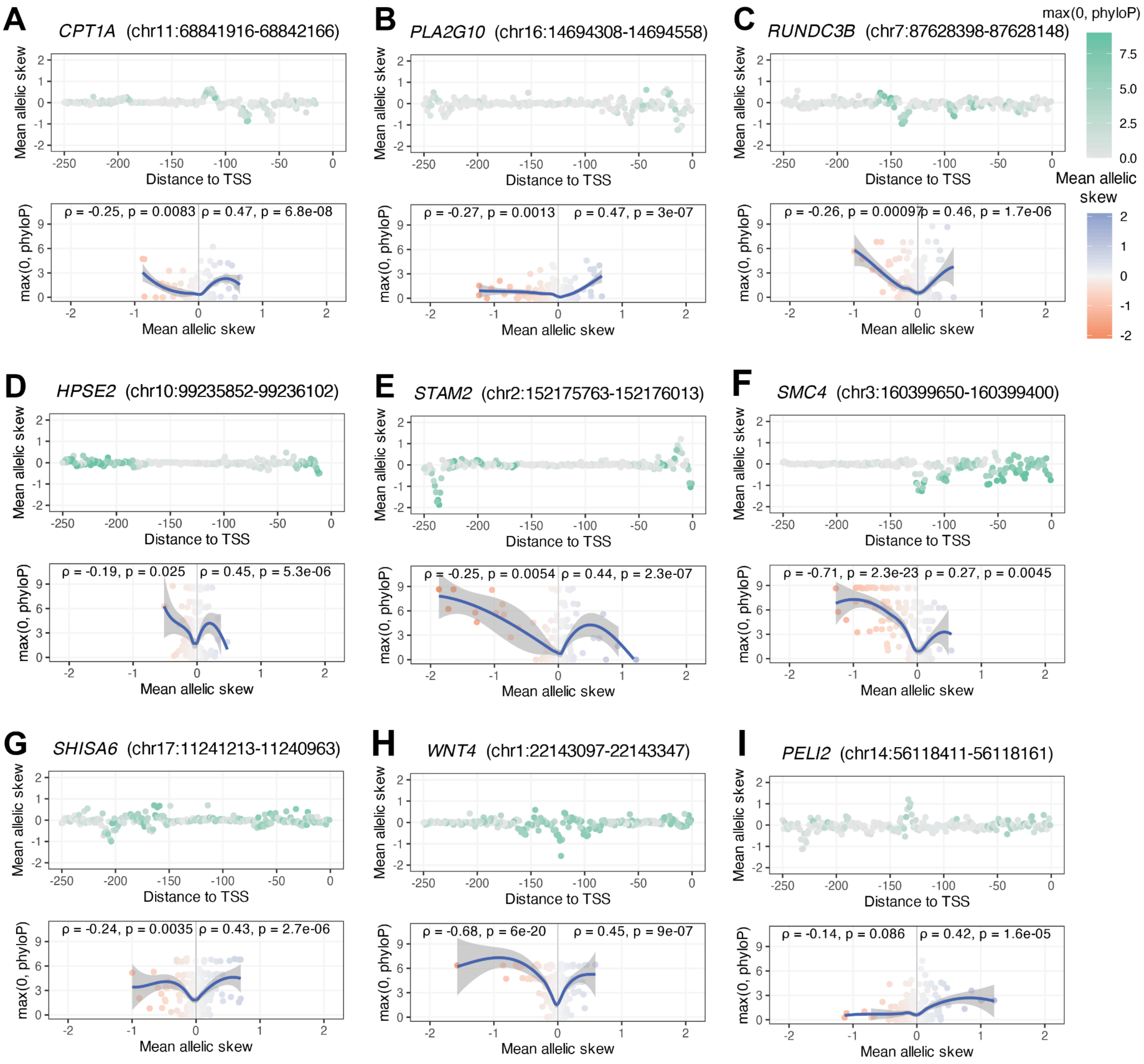


**Supplementary Fig. 27: Additional promoters in top 10 by their correlations between constraint and positive allelic skew**

Plots of allelic skew (mean $\log_2$FC of the three possible SNVs at each position) and constraint (max(0, phyloP)) by distance to TSS (top). Spearman correlation ($\rho$ and p-values) between constraint and allelic skew for mutations with positive allelic skew (bottom left) or mutations with positive allelic skew (bottom right). Promoters in the top 10 by positive allelic-skew $\rho$ excluding *EMLIN2* (see **Fig. 4I**). LOESS curve with 95% CI. Missing data overlap annotated exons and proximal intronic splice regions, including those of alternative transcripts and other genes.

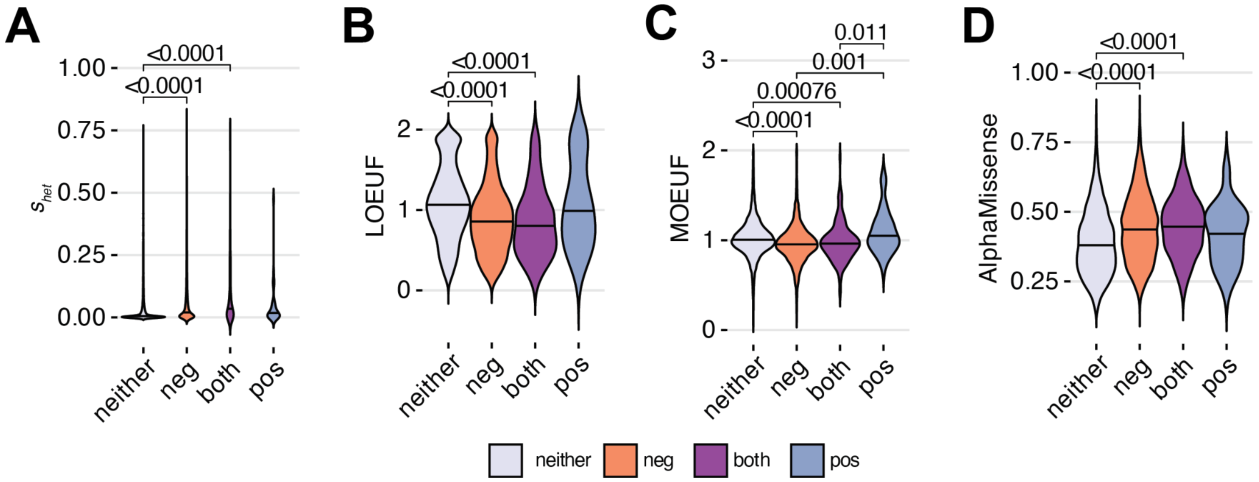


**Supplementary Fig. 28: Gene-level constraint score distributions for individual promoters classified by their correlations between allelic skew and constraint**
Distributions of gene-level scores of coding constraint for promoters with significant correlation between allelic skew and constraint (max(0, phyloP)) for negative allelic skew mutations (neg), positive allelic skew mutations (pos), neither, or both. **A)** $s_{het}$ LoF-intolerance, **B)** gnomAD LOEUF LoF-intolerance, **C)** gnomAD MOEUF missense-intolerance, and **D**) AlphaMissense average scores. Wilcoxon signed-rank test $p$-values.

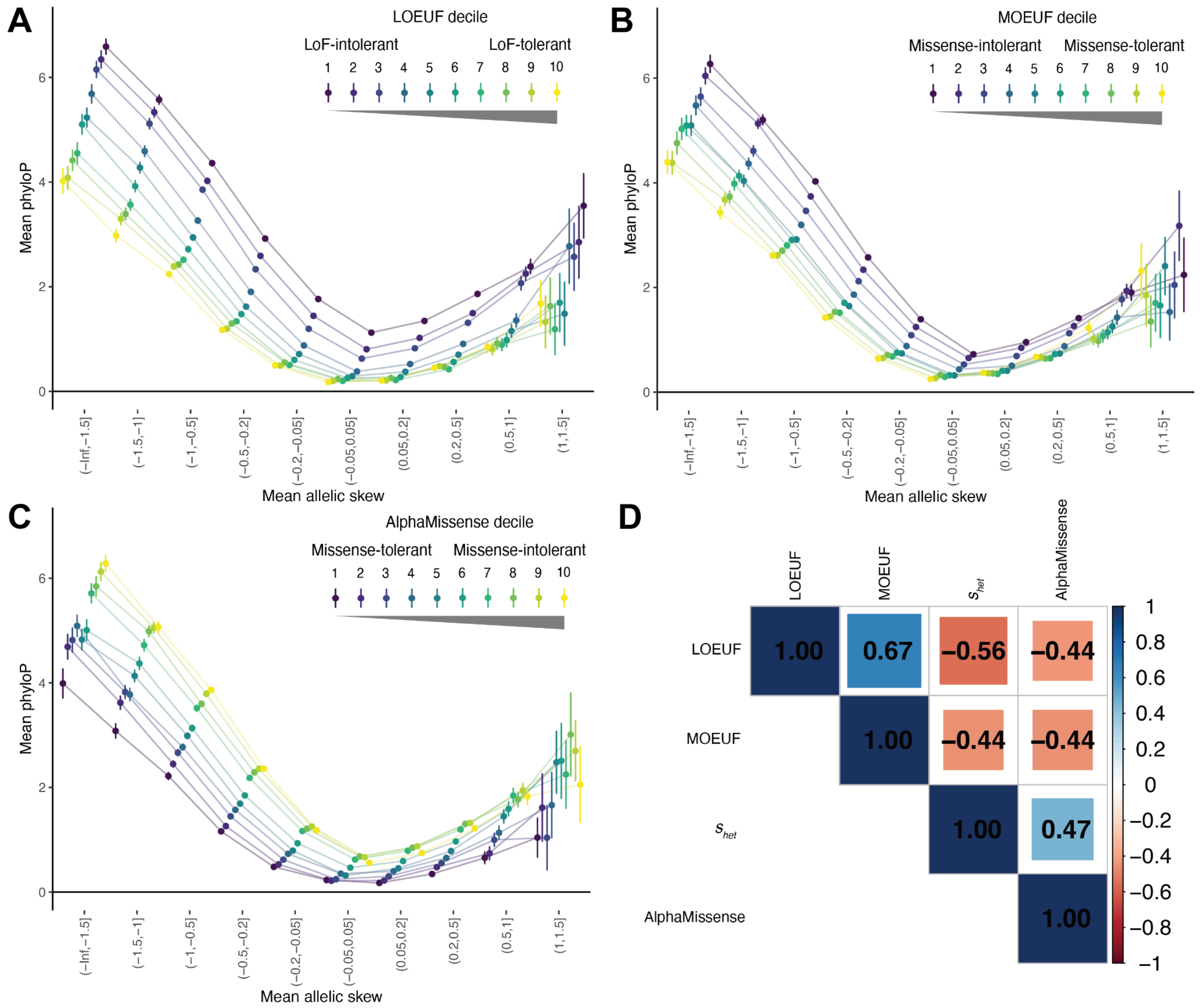


**Supplementary Fig. 29: Evolutionary constraint, gene-level coding constraint, and functional effects across alternative gene-level constraint cores**

The base-level constraint (mean phyloP) of promoter mutations by allelic skew (mean $\log_2$FC across three cell lines) stratified by alternative measures of gene-level coding constraint: **A)** gnomAD LOEUF deciles of LoF-intolerance, **B)** gnomAD MOEUF deciles of missense intolerance, and **C)** AlphaMissense average scores of all possible missense mutations. Error bars represent 95% CIs. **D)** Pairwise Spearman's $\rho$ between $s_{het}$, LOEUF, MOEUF, and AlphaMissense gene-level constraint scores.
